# Functionally diversified BiP orthologs control body growth, reproduction, stress resistance, aging, and ER-Phagy in *Caenorhabditis elegans*

**DOI:** 10.1101/2025.01.14.633073

**Authors:** Nicholas D. Urban, Shannon M. Lacy, Kate M. Van Pelt, Benedict Abdon, Zachary Mattiola, Adam Klaiss, Sarah Tabler, Matthias C. Truttmann

**Author notes:** To whom correspondence should be addressed: BSRB, 109 Zina Pitcher Place, Ann Arbor 48109, MI.

## Abstract

Cellular systems that govern protein folding rely on a delicate balance of functional redundancy and diversification to maintain protein homeostasis (proteostasis). Here, we use *Caenorhabditis elegans* to demonstrate how both overlapping and divergent activities of two homologous endoplasmic reticulum (ER)-resident HSP70 family chaperones, HSP-3 and HSP-4, orchestrate ER proteostasis and contribute to organismal physiology. We identify tissue-, age-, and stress-specific protein expression patterns and find both redundant and distinct functions for HSP-3 and HSP-4 in ER stress resistance, reproduction, and body size regulation. We show that only HSP-3 overexpression is sufficient to improve longevity and that loss of HSP-3 or HSP-4 during distinct stages of the worm cycle or specific tissues have opposing effects on worm lifespan. Furthermore, we find that loss of HSP-4, but not HSP-3, improves tolerance to protein aggregation induced-stress by activating ER-Phagy through the engagement of IRE-1 and the putative ER-Phagy receptor, C18E9.2. Mechanistically, we show that de-repression of IRE-1 via HSP-4 dissociation allows for direct inhibition of C18E9.2- mediated ER-Phagy and demonstrate that a conserved orthologous mechanism involving the respective human orthologs, BiP, Sec-62, and IRE-1, contributes to ER proteostasis regulation in human cells. Taken as a whole, our study demonstrates that functional diversification of orthologous proteins within a single organelle is an efficient mechanism to maximize stress resilience while also defining a novel link between ER-phagy and proteostasis regulation.

## Introduction

Gene duplications generate paralogs that encode distinct proteins. This process drives genetic evolution and helps ensure organismal survival^1–3^. Although structurally similar, evolutionary pressure pushes the development of distinct functions of these seemingly redundant proteins^4^. Current literature suggests only ∼25% of duplicated genes can compensate for loss of function of their respective counterpart—however, almost always in a severely reduced capacity^5,6^.

Heat shock proteins (HSPs) are highly conserved molecular chaperones which maintain protein homeostasis (proteostasis) under physiological and stress-induced conditions^7^. These proteins are categorized based on their molecular size and distinguished by subcellar compartmentalization and function. Chaperones of the 70 kilodalton heat shock protein (HSP70) family have been implicated in cardiovascular disease^8,9^, various cancers^10–12^, and numerous neurodegenerative diseases such as Alzheimer’s^13^ disease and Parkinson’s disease^14^.

As eukaryotes evolved, the proteome diversified and became more complex, accumulating proteins rich in beta folds or amino acid expansion sequences. To support proper folding of these often misfolding-prone and disease-affiliated proteins, protein quality control systems expanded and differentiated. One key adaptation included the expression of a functionally expanded panel of HSP70 family chaperones^15,16^. For example, the nematode, *Caenorhabditis elegans*, expresses at least seven, and humans at least thirteen, HSP70 family chaperones^17^, with many sharing overlapping subcellular localization^18^. Homology-based functional predictions and shared expression patterns suggest at least partial functional overlap between these distinct chaperones.

However, reliance on such predictions without experimental validation may lead to incomplete or inaccurate conclusions of paralogous protein function(s), while prohibiting the identification of newly acquired protein activities.

The endoplasmic reticulum (ER)—the main secretory organelle of the cell—is required across taxa to properly fold nascent proteins and regulate interorganellar signaling to maintain cellular proteostasis. These processes are essential requirements to sustain life^19,20^. Misfolded proteins within the ER are detected and refolded by HSPs or cleared through evolutionarily conserved mechanisms which navigate crosstalk between the ER and other organelles, such as the unfolded protein response of the ER (UPR^ER^)^21^ and ER-selective autophagy (ER-Phagy)^22^.

The UPR^ER^ is mediated by three canonical ER transmembrane proteins: Inositol-Requiring Enzyme 1 (IRE-1), Protein Kinase R-like ER Kinase (PERK), and Activating Transcription Factor 6 (ATF-6). ER stress triggers specific transcriptional responses unique to each UPR^ER^ mediator to mitigate protein misfolding burden^23^. Furthermore, increased protein misfolding increases the size of the ER through the UPR^ER^, a process critical to sufficiently alleviate proteotoxic stress^24,25^.

ER-phagy is the degradation of ER-specific cargo through canonical autophagy mechanisms^22^. ER-phagy requires transmembrane receptors to recruit cytosolic autophagy-related proteins to the ER membrane. These proteins assist in engulfing the ER-membrane (and luminal components) into autophagosomes for subsequent degradation and recycling. This process rids the ER of misfolded proteins and alleviates protein misfolding burden, as well as reduces the size of the ER to its homeostatic size following the resolution of ER stress (a process dubbed recovER-Phagy)^22,26^. ER-Phagy/recovER-phagy and the UPR^ER^ are co-regulated; however, the mechanistic details of this relationship remain elusive^27^.

In human cells, the essential ER-resident HSP70 family chaperone, Binding-Immunoglobulin Protein (BiP), contributes to protein (re)folding while regulating UPR^ER^ activation and selective forms of ER-Phagy. In unstressed cells, binding of BiP to IRE-1, PERK, or ATF-6 prevents the activation of each of the three arms of the UPR^ER^. Upon increased protein misfolding, BiP dissociates from the UPR^ER^ mediators, thus derepressing the UPR^ER^ and activating transcriptional programs aimed at restoring proteostasis such as increasing chaperone production and promoting ER expansion^28–30^. BiP also activates FAM134B-mediated ER-Phagy to help efficiently restore ER proteostasis^31,32^. Still, how a single HSP70 family chaperone fulfills this multitude of critical activities, as well as the extent of BiP’s regulation of ER-mediated signaling mechanisms, remains incompletely understood.

Of the seven canonical HSP70 family chaperones found in *C. elegans*, 4 are cytosolic (HSP-1, C12C8.1, F44E5.4, F44E5.5) and a single HSP70 chaperone is found in the mitochondria (HSP-6); but, unlike humans, two distinct BiP-like proteins are found exclusively in the ER (HSP-3, HSP-4)^17^. Although *hsp-3* is transcribed at higher levels than *hsp-4*, previous work showed *hsp-3* and *hsp-4* transcripts are both stress responsive and partially regulated by the UPR^ER^ ^33,34^. However, despite a well-described transcriptional relationship between these two chaperones suggesting potential functional compensation^34^, evidence showing HSP-4, but not HSP-3, is required for proper cuticle formation during worm development indicates currently unknown independent functions of these proteins^35^.

In this study, we define in detail the benefits of functional redundancy and diversification in HSP70-mediated proteostasis regulation by assessing the individual contributions of the two *C. elegans*’ BiP orthologs, HSP-3 and HSP-4, as a model case. We demonstrate both divergent and convergent functions of HSP-3 and HSP-4 in ER and non-ER-stress resilience, body growth, reproduction, lifespan, and ER-Phagy regulation. Finally, we identify a novel mechanistic link between HSP-4’s regulation of IRE-1 and C18E9.2-mediated recovER-phagy which is conserved in human cells.

Taken as a whole, our findings showcase how functionally diversified HSP70 chaperones within the ER efficiently co-operate to optimize stress resilience and maintain proteostasis.

## Methods

### C. elegans maintenance and strain

Worms were maintained at 20 °C on peptone rich (8P) nematode growth media (NGM) plates spotted with OP50-1 *E. coli* (Caenorhabditis Genetics Center (CGC)). All experiments were performed at 20 °C unless otherwise stated. Worms were fed for at least two generations before being used for experiments. A detailed list of strain information can be found in Table S1.

### Cloning and generation of transgenic strains

*hsp-3* and *hsp-4* full-length transcripts were amplified from *C. elegans* cDNA by PCR, purified, and cloned into pMT686^36^. The following primers were used: *hsp-3*: (fwd 5-acgtcccagactacgctggcATGAAGACCTTATTCTTGTTGGGCTTGATC-3’; rev: 5’- ttactcattttttctaccggTTAGAGCTCGTCCTTGTCGTCAGATC-3’); *hsp-4*: (fwd: 5’- acgtcccagactacgctggcATGAAAGTTTTCTCGTTGATTTTGATTGCC-3’; rev: 5’- ttactcattttttctaccggTTACAGTTCATCATGATCCTCCGATGG-3’. Plasmid sequences were verified by Sanger sequencing. *hsp-3* and *hsp-4* overexpression plasmids were injected into wild-type (N2) hermaphrodite worms alongside the co-injection marker, *Pmyo-2::GFP,* to generate strains carrying extrachromosomal arrays (Table S1). Strains PHX4377 (*hsp-3::wrmScarlet*) and PHX4415 (*hsp-4::wrmScarlet*) were generated using CRISPR/Cas9 technology^37^. All microinjections were performed by SunyBiotech (Fujian, China). Extrachromosomal arrays were integrated by UV irradiation. We were unsuccessful in integrating *mtmEx74* after 5 attempts. N2 worms (Bristol, CGC) were used as wild type controls unless otherwise stated. All strains were backcrossed at least 3-5x to N2 worms prior to use in experiments.

### Gene knockdown via RNA interference

HT115 *E. coli* expressing siRNA against the gene of interest are from either the Vidal^38^ or Ahringer siRNA library^39^ and described in Table S2. To prepare siRNA, single *E. coli* colonies were inoculated into 5x cultures of Lauria broth (LB) media (Genesee Scientific, Cat #11-118) supplemented with 100 mg/mL carbenicillin (diluted 1:1000, GoldBio, Cat #C-103-25) and shaken overnight (37 °C). The following day, the 5x culture was spun down at 2,633 g for 10-15 minutes. The supernatant was removed, and the bacterial pellet was resuspended in 1x fresh LB media (e.g., 5 mL of overnight culture resuspended in 1 mL of fresh LB). 100 mg/mL carbenicillin (diluted 1:1000) and 1M isopropyl β-D-1-thiogalactopyranoside (IPTG) (diluted 1:200; Dot Scientific, Cat #DSI5600-25) were added to the 1x culture. Bacteria were spotted on standard nematode growth media (NGM) plates supplemented with 1M IPTG (1:1000, induces siRNA expression), 100 mg/mL carbenicillin (1:1000, antibiotic) and 10 mg/mL nystatin (1:1000, antifungal; Dot Scientific, Cat #DSN82020-10). Once dry, plates were used immediately or kept at 4 °C for no more than 2-3 days. For all double knockdown experiments (e.g., *hsp-4/C18E9.2*), equal amounts of siRNA with similar OD were mixed in a 1:1 ratio to account for any potential dosage effects on siRNA efficacy and single knockdown controls were mixed with equal volumes of *pos-1* siRNA as indicated in the figure. Knockdown of all siRNA used in this study were validated previously (including *hsp-3* and *hsp-4* siRNA)^36^ or via RT-qPCR (see *RT-qPCR* and Figs. S11/S12) using the primers listed in Table S2.

### Synchronization via hypochlorite treatment

Heterogeneously aged worms were washed in 1-2 mL of M9 buffer and collected in a 1.7 mL Eppendorf tube. Worms were gravity separated for 3-5 minutes and washed with fresh M9 (repeated 3-4x). M9 was then removed and replaced with 1 mL of hypochlorite solution (56 mL ddH_2_O, 14.4 mL 5N NaOH, 7.2 mL 7.55% bleach). Tubes were shaken at 1200 rpm in a tabletop heat block at 20 °C for 7-10 minutes until worm bodies were not observed. Eggs were spun at 1600 rcf for 30 seconds, bleach was removed, and eggs were washed with 1 mL of M9. Washing was repeated twice. Eggs were washed a final time with M9 and spun at 2900 rcf for 3 minutes, resuspended in ∼100 μL of M9, and plated.

### C. elegans development assays

HT115 *E. coli* expressing the designated siRNA were prepared and spotted onto NGM/RNA interference plates as described above (see *Gene knockdown via RNA interference*). Worms were synchronized and eggs were spotted onto NGM/RNA interference plates. Total eggs were counted, and plates were incubated in a 20 °C incubator for the time described. The number of larval stage 4 worms or older were then scored and compared to the number of eggs on the plate.

### RT-qPCR analysis

N2 Worms were synchronized and grown on the described siRNA for 72 hours unless otherwise stated. For RNA extraction, ∼500-1000 worms were washed 3x via gravity separation in M9 and lysed using the Direct-zol^TM^ RNA MiniPrep Plus Kit (Zymo Research, Cat #R2070) following the manufacturers instruction in a Qiagen TissueLyser III (5 minutes, 30 Hz). cDNA generation was completed using Applied Biosystems™ High-Capacity cDNA Reverse Transcription Kit (Fisher Scientific, Cat #43-688-14) following the manufacturer’s instructions. RT-qPCR analysis was done using PowerUp™ SYBR™ Green Master Mix for RT-qPCR (Applied Biosystems™, Cat #A25742) master mix. Primers used for RT-qPCR analysis are listed in Table S2.

### C. elegans fluorescent image acquisition and analysis

Worms were immobilized in 10 μL of 25 mM tetramisole hydrochloride (Santa Cruz, Cat #sc-215963) dissolved in M9 buffer, placed on a 2% agarose pad on a glass microscope slide, and covered with a round glass coverslip. For quantification analysis, worms were imaged using a Keyonce BZ-X700 fluorescent microscope at 10x magnification. Image analysis was done using a custom generated semiautomatic pipeline in Cell Profiler (version 4.2.1). This pipeline is available upon request.

Integrated intensity per body area for each worm was reported. Representative confocal images were taken using a Zeiss LSM 800 confocal laser scanning microscope. Representative worms were straightened using the “Straighten” feature in Fiji (version 2.14).

### Thapsigargin and tunicamycin stress and survival assays

Worms were synchronized and grown on the described siRNA for 72 hours. Thapsigargin (Invitrogen, Cat #T7458) was dissolved in DMSO to a final concentration of 1mM. Tunicamycin (Tocris Biosciences, Cat #35-161-0) was dissolved in basic M9 (pH 9.5-10) to a final concentration of 1.0 mg/mL. 48 hours post-synchronization, fresh siRNA-spotted plates were treated ectopically with 0.1 μg of thapsigargin or 10 μg of tunicamycin per mL of NGM agar or equivalent volume of buffer and stored at 4 °C overnight. For both florescence imaging and survival assays, day 1 adult worms were transferred to a chemical or solvent treated plate and kept at 20 °C.

### Bulk RNA-sequencing and analysis

Synchronized N2 worms were exposed to siRNAs as described above and maintained for 48 hours (∼larval stage 4). Animals were collected, washed repeatedly with M9 as described above, snap frozen in liquid nitrogen, and stored at-80 °C until processing. RNA extraction was done as described above (see *RT-qPCR analysis*). Three biological replicates were collected and then processed by the University of Michigan Advanced Geonomics Core. The library was prepared using the NEBNext Poly(A) mRNA Magnetic Isolation Module (NEB, Cat #E7490L) and NEBNext Ultra II RNA Library Prep Kit (NEB, Cat #E7770L). Sample sequencing was done using an Illumina NovaSeq 6000 S4. The reference genome, WBcel235, was used to map reads using STAR v2.7.8a^40^ and count estimates were assigned using RSEM (v1.3.3)^41^. Differential expression analysis was performed with edgeR (v3.19, Bioconductor)^42^. ShinyGO (v0.80)^43^ was used to generate graphical representations of enriched gene ontology (GO) terms. Volcano plots were generated using the EnhancedVolcano package (v.1.22.0)^44^. 1.5-fold change was used as a cutoff for differential expression. Raw data of RNA-sequencing experiments were disclosed in *Van Pelt et al.*^36^ and reanalyzed for this publication. RNA-sequencing data can be found in Tables S6-S8.

### Timed egg laying (fecundity) assays

Worms were synchronized onto HT115 *E. coli* expressing the described siRNA and maintained for 72 hours. Worms were transferred to 35mm NGM-IPTG plates spotted with 150 μL of the described siRNA-expressing bacteria and maintained at 20 °C for 3.5 hours (1 worm per plate). Following the timed egg lay, worms on the same siRNA were then transferred to a shared 60mm NGM-IPTG plate of the described siRNA and maintained at 20 °C until the next timepoint. Total number of eggs laid on each plate were then counted. Timed egg lays began 72 hours (Day 1), 144 hours (Day 3), and 192 hours (Day 5) after synchronization.

### Body area assays

Worms were synchronized onto HT115 *E. coli* expressing the described siRNA. Animals were imaged 72 hours (Day 1), 144 hours (Day 3), and 192 hours (Day 5) after synchronization. Worms were transferred to fresh siRNA plates after imaging was complete on Day 1 and Day 3 to maintain cohort synchronization. Image analysis was done using a custom generated semiautomatic pipeline in Cell Profiler (version 4.2.1) described above (see C. elegans *florescent image acquisition and analysis*).

### C. elegans lifespan assays

Worms were synchronized and transferred to assay plates 72 hours. For knockdowns “From Birth”, eggs were spotted directly on HT115 *E. coli* expressing siRNA and remained on that siRNA for the duration of the experiment. For knockdowns during “Adulthood Only”, worms were synchronized onto *pos-1* siRNA against and transferred after 72 hours to the indicated siRNA. For knockdowns during the “Larval Only” stages, worms were synchronized onto the described siRNA and maintained for 72 hours, then transferred to *pos-1* siRNA for the remainder of the experiment. Worms were transferred 5-10 minutes to an intermediate plate without bacteria to rid worms of previous siRNA before being transferred to fresh plates. Worms were transferred every ∼48 hours to fresh plates for the first 6-7 days as needed to maintain a synchronized population as needed. Worms were scored as alive or dead every 1-2 days; worms which failed to produce movement spontaneously or after being gently prodded with a platinum worm pick were considered dead and removed. Unless otherwise stated, all siRNA experiments were “From Birth”. Each biological replica is the aggregate of 3 technical replicas (individual NGM/siRNA plate). Number of animals and median lifespan data per experiment is in Table S4.

### GMC101 paralysis assays

GMC101 were synchronized onto the indicated siRNA and grown at 20 °C for 72 hours. Worms were then moved to 60mm plates containing fresh siRNA and transferred to either a 20 °C (non-inducive) or 25 °C (inducive) incubator. Animals were scored for paralysis every 1-2 days, and paralyzed worms were removed from the plate. Paralysis was defined as the inability to complete a full body movement in either direction spontaneously or when prodded with a platinum wire worm pick. If needed, worms were transferred to fresh plates after 2 and 4 days of adulthood to maintain a synchronized cohort and ensure adequate food availability. After all control worms at the inducive temperature paralyzed, the remaining non-paralyzed worms were counted and censored. Number of animals paralyzed/censored and day of median paralysis data per experiment is in Table S5.

### C. elegans protein lysate preparation

Synchronized worms were washed with sterile M9 into a 1.7mL Eppendorf tube and washed 2-3x using gravity separation. Thereafter, the remaining M9 was removed after brief centrifugation. Worms were snap frozen in liquid nitrogen and stored at-80 °C. Frozen samples were resuspended in 150-200 μL of sterile filtered worm lysis buffer (HEPES (20mM, 7.4pH), NaCl (20mM), MgCl_2_ (200mM), and Nonidet P-40 (0.5%)) spiked with protease inhibitor cocktail (Pierce^TM^ Protease and Phosphatase Inhibitor Mini Tablets, EDTA-free, ThermoFisher, Cat #A32961), transferred to reinforced tubes containing a steel lysis ball, and mechanically lysed using a Qiagen TissueLyser III (7.5 minutes, 30 Hz). Lysate was cleared at 16,100 rcf (4 °C, 15 minutes) twice, transferring to a precooled tube after each transfer. The soluble fraction was collected, and protein lysate concentration was determined using the Pierce^TM^ BCA Protein Assay Kit (ThermoFisher, Cat #23227) following the manufacture’s instruction. For western blot, equal amount of protein lysate was brought to equal volume with worm lysis buffer and combined with 4x Laemmli Protein Sample Buffer (Bio-Rad, Cat #1610747) prepared per the manufacturer’s instruction. Samples were boiled for 5 minutes at 100 °C, allowed to cool, and excess moisture was spun down briefly using a tabletop centrifuge. Samples were then used immediately or stored at-20 °C.

### Immunoblotting

Prepared protein lysates were run on 8% (PHX4377/PHX4415), 13.5% (DA2123), or 10% (A549 cells) SDS polyacrylamide gels and transferred to PVDF membranes using the Bio-Rad Trans-Blot^®^ Turbo System (Bio-Rad, Cat #1704150) and Trans-Blot^®^ Turbo RTA Transfer Kit, PVDF (Bio-Rad, Cat #1704272). Membranes were then trimmed and blocked at least 1 hour at room temperature with gentle shaking. The blocking buffer was then removed, and membranes were incubated with primary antibody overnight at 4 °C with gentle rocking. Primary antibody was then removed, and membranes were washed 3x (5-10 minutes each) with 0.1% TBS-T. Membranes were then incubated and gently rocked with HRP-conjugated secondaries one hour. Antibody was removed and membranes were washed 3x with 0.1% TBS-T (5 minutes each). HRP-conjugates were detected using ProSignal Dura ECL Reagent (Prometheus Protein Biology Products, Genesee Scientific, Cat #20-301) following the manufacturer’s instructions and imaged using an Invitrogen iBright1500. If required, membranes were stripped using *OneMinute^®^* Western Blot Stripping Buffer (GM Biosciences, Cat #GM6001) following the manufacturers instruction, washed vigorously with ddH_2_O, then rehydrated in 0.1% TBS-T for 10 minutes. Membranes were then treated as described above.

Immunoblot quantification was done using Fiji (version 2.14). A list of blocking buffers, antibodies, and dilutions used in this study are described in Table S3.

### A549 cell maintenance and tunicamycin treatment

A549 cells were obtained from ATCC and cultured in DMEM media supplemented with 10% FBS and 100 μg/mL penicillin and streptomycin mixture. Cells were maintained at 37 °C and 5% CO_2_. For ER stress assays, cells were plated in 10 cm plates overnight and subsequently treated for the listed amount of time with 10 μg/mL of tunicamycin dissolved in DMSO (Tocris Biosciences, Cat #35-161-0) before harvesting for immunoprecipitation assays.

### Co-immunoprecipitation assays

A549 cells were lysed with RIPA buffer (Sigma-Aldrich, Cat #R0278) containing protease inhibitor (Pierce^TM^ Protease and Phosphatase Inhibitor Mini Tablets, EDTA-free, ThermoFisher, Cat #A32961) for 15 minutes on ice and sonicated at 20% for 30 seconds on ice. Cell debris was spun down 16,100 rcf (4 °C, 15 minutes), and protein quantification was performed using the DC^TM^ Protein Assay kit (Bio-Rad, Ca #5000112). Cell lysates were pre-cleared using Pierce^TM^ Protein A Magnetic beads (ThermoFisher, Cat #88846) for 1 hour at 4 °C, and the beads were removed using a magnetic rack.

200 μg of cell lysate was incubated with 3 μL of IRE-1 antibody (Cell Signaling, 14C10, Cat #3294) overnight at 4 °C. 20 μL of Protein A Magnetic beads were added to the mixture for 2 hours at 4°C with gentle rocking. The immunoprecipitated proteins were isolated from the cell lysate mixture using a magnetic rack and isolated beads were washed three times with the cell lysis buffer. The bound proteins were released from the beads by adding SDS loading buffer and boiled for 10 minutes. The membranes were probed using the antibodies listed in Table S3 and imaged as described above (see *Immunoblotting*). The tunicamycin treatment and immunoprecipitation assay was performed three separate times and the bands were quantified using the online Image Lab software from Bio-Rad (v 6.1.0).

### Statistical analyses

All statistical analysis was done in Graphpad Prism (v10.4.0) and described in the figure legend. For super plots, the average of each biological replicate was used for analysis. Error bars represent standard deviation unless otherwise stated in the figure legend.

## Results

### HSP-3 and HSP-4 proteins have distinct temporal and spatial expression profiles

*C. elegans* is equipped with two ER-resident HSP70 family chaperones, HSP-3 and HSP-4. In contrast, human cells express a single ER-resident HSP70 family chaperone, BiP, to orchestrate ER protein folding and maintain ER proteostasis. To better understand the roles of HSP-3 and HSP-4 *in vivo*, we generated HSP- 3::wrmScarlet (PHX4377) and HSP-4::wrmScarlet (PHX4415) fusion protein reporters using CRISPR/Cas9. The wrmScarlet sequence was inserted at the endogenous C-termini of the respective gene immediately upstream of the XDEL motif. Using functional assays and RT-qPCR, we confirmed HSP-3::wrmScarlet and HSP-4::wrmScarlet were functional, expressed similarly to HSP-3 and HSP-4 in wild type (WT) animals, and did not interfere with worm development (Fig S1A-C).

Using these reporter strains, we first determined the temporal expression of both HSP-3 and HSP-4 via immunoblotting. We found HSP-3::wrmScarlet protein levels were greater than HSP-4::wrmScarlet throughout development and in day 1 adult worms. Relative HSP-3::wrmScarlet abundance peaked 48 hours after development during the L3/L4 stages and was reduced upon entering adulthood; in contrast, HSP-4::wrmScarlet protein levels were lowest throughout larval development and highest during early adulthood (Fig. 1A-B, Fig. S1D). HSP-3::wrmScarlet and HSP-4::wrmScarlet levels increased throughout the worm body with increased age; maximum HSP-3::wrmScarlet expression was observed in day 5 adults, whereas peak HSP-4::wrmScarlet protein levels occurred in day 10 adults (Fig. S2A-D).

**Fig. 1.**
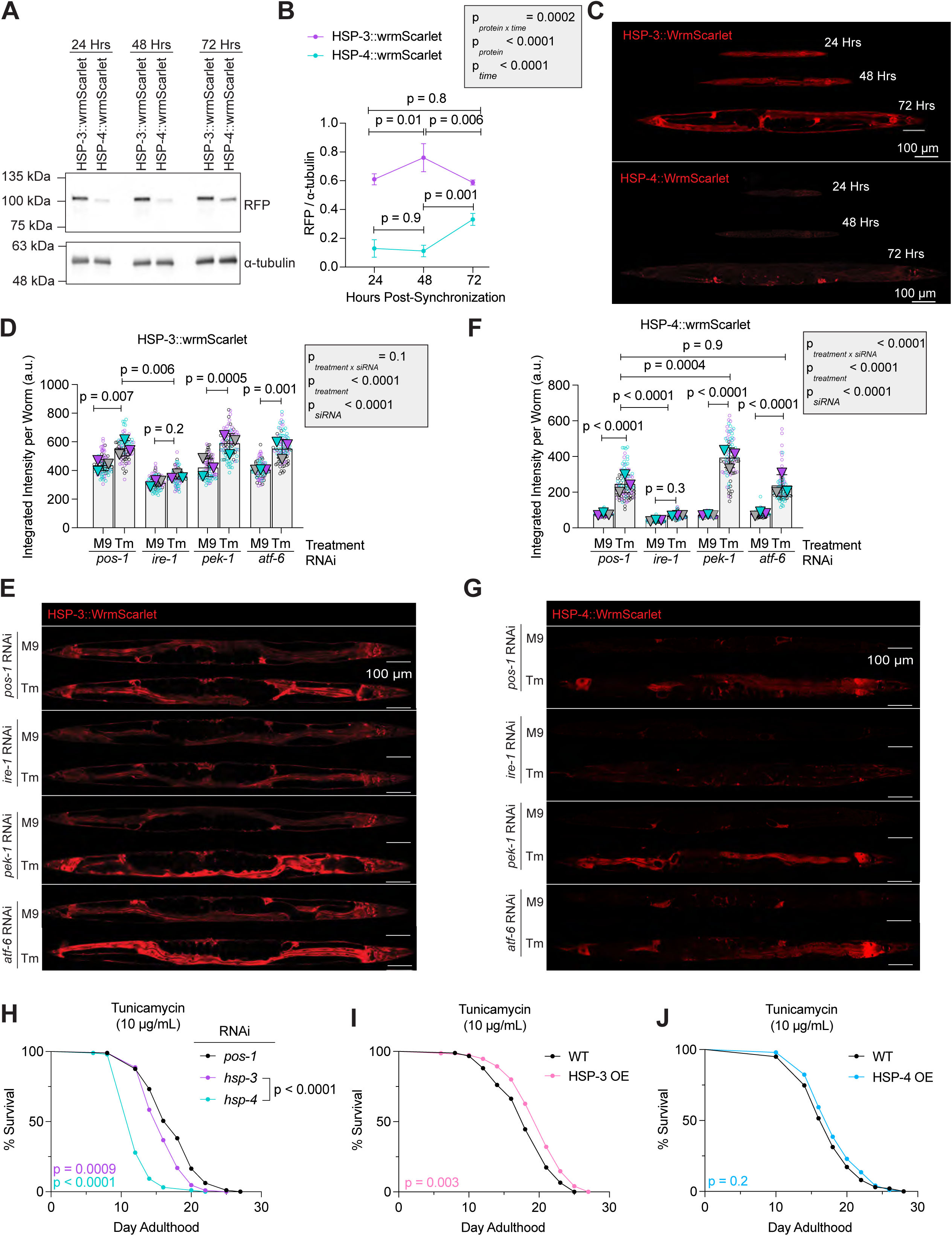
HSP-3 and HSP-4 show distinct spatial-temporal expression patterns and responses to ER-stress. (A-B). Representative western blot (A) and quantification (B) of PHX4377 (HSP-3::wrmScarlet) or PHX4415 (HSP-4::wrmScarlet) grown on *pos-1* siRNA at 20 °C and collected 24’, 48’, and 72’ after synchronization. Membranes were probed with an anti-RFP (Chromotek, Cat #6g6) and anti-alpha tubulin (DSHB, Cat #12G10) antibody. (C). Confocal images of PHX4377 and PHX4415 grown on *pos-1* siRNA collected 24’, 48’, and 72’ after synchronization. Worms were kept at at 20 °C. (D-E). Quantification (D) and representative confocal images (E) of PHX4377 grown on indicated siRNA and treated with M9 or Tm. (F-G). Quantification (F) and example confocal images (G) of PHX4415 grown on indicated siRNA and treated with M9 or Tm. (H). Chronic Tm stress survival assay of WT (N2) worms grown on indicated siRNA. (I- J). Chronic Tm stress survival assays of WT and worms overexpressing HSP-3 (MTX203) (I) or HSP-4 (MTX291) (J). For C, E, and G, images were taken at the midline. *(B, D-F).* For H-L, the number of animals and median lifespan data per experiment is in Table S4. *Two-way ANOVA followed by Tukey’s multiple comparisons test. (H-J). Log-rank Mantel-Cox test. p < 0.05 is considered statistically significant*.

Next, we used confocal microscopy to determine the spatial distribution of HSP- 3::wrmScarlet and HSP-4::wrmScarlet. Tissue distribution of HSP-3::wrmScarlet and HSP-4::wrmScarlet were ubiquitous throughout the body during larval development (Fig. 1C). In the early adult hermaphrodite, HSP-3::wrmScarlet was highly expressed throughout the worm body, with peak expression in the intestine and neurons. HSP- 4::wrmScarlet was found in all tissues as well but at lower levels, with maximal expression observed in the spermatheca, the site of spermatogenesis in the adult hermaphroditic worm (Fig. 1C). These results show HSP-3 and HSP-4 protein expression is both distinct and dynamic, adapting to age-dependent needs.

### HSP-3 and HSP-4 show varied responses to ER stressors and the UPR^ER^

To define how ER stress controls HSP-3::wrmScarlet and HSP-4::wrmScarlet expression and tissue distribution, we treated day one adult worms with the SERCA inhibitor, thapsigargin (Tg), or the N-linked glycosylation inhibitor, tunicamycin (Tm) for 24 hours. Tg treatment increased HSP-4::wrmScarlet, but not HSP-3::wrmScarlet levels (Fig. S2E-H), whereas Tm exposure increased both HSP-3::wrmScarlet and HSP- 4::wrmScarlet protein (Fig. 1D-G). Expression increases were predominantly observed in the intestine and neurons for both proteins. We next assessed the involvement of the UPR^ER^ on regulating HSP-3::wrmScarlet or HSP-4::wrmScarlet protein levels in basal and ER stress conditions. Knockdown of *ire-1* reduced HSP-3::wrmScarlet and HSP- 4::wrmScarlet protein levels in controls; however, loss of *pek-1* or *atf-6* did not alter protein levels of buffer-treated worms of either genotype (Fig. 1D-G, Fig. S2E-H).

Knockdown of any of the UPR^ER^ sensors did not alter HSP-3::wrmScarlet levels after Tg treatment (Fig. S2E-F), but IRE-1 was required to upregulate HSP-3::wrmScarlet following Tm exposure (Fig. 1D-E). Loss of *ire-1* or *atf-6* restricted the upregulation of HSP-4::wrmScarlet in response to Tg treatment (Fig. S2G-H), whereas only IRE-1 was required to upregulate HSP-4::wrmScarlet following Tm exposure (Fig. 1F-G). However, *pek-1* knockdown further increased the upregulation of HSP-4::wrmScarlet in response to Tm compared to control siRNA, with strong expression observed in the intestine (Fig. 1F-G).

These data indicate that IRE-1 is required for HSP-3 and HSP-4 protein expression in both stress-free and ER-stress environments. Our results further confirm divergent ER stress-and UPR^ER^-specific responses of HSP-3 and HSP-4.

### HSP-3 *and* HSP-4 are required to tolerate ER stress, but only HSP-3 is sufficient to increase ER stress tolerance

Based on our results showing HSP-3 and HSP-4 exhibit significant differences in spatial expression, we hypothesized both proteins may have adopted unique roles in proteostasis maintenance and worm physiology. To test this prediction, we assessed if HSP-3 or HSP-4 are required for worm survival following sustained ER stress induced by chronic Tm exposure. We measured survival of synchronized WT worms exposed to control (*pos-1*), *hsp-3* or *hsp-4* siRNA and treated with Tm starting on day 1 of adulthood. Knockdown of *hsp-3* or *hsp-4* reduced survival after Tm treatment compared to control siRNA, but worms lacking *hsp-4* were significantly shorter lived than worms lacking *hsp-3* when treated with Tm (Fig. IH, Fig. S3A). Next, we overexpressed HSP-3 and HSP-4 to determine if presence of either chaperone in excess is sufficient to buffer against Tm-induced ER stress (Table S1). HSP-3 overexpression (OE) increased ER stress tolerance (Fig. 1I, Fig. S4A), whereas HSP-4 OE did not affect survival following Tm exposure (Fig. 1J, Fig. S5A). These results confirm that HSP-3 and HSP-4 are required to buffer against prolonged (chronic) ER stress, but only HSP-3 OE is sufficient to increase ER stress resilience.

### Loss of HSP-3 or HSP-4 elicits different transcriptional responses

Similar to BiP in human cells, HSP-3 and HSP-4 are proposed to regulate UPR^ER^-mediated gene transcription by repressing IRE-1, PEK-1, and ATF-6 in *C*. *elegans* ^45^. To test this hypothesis, we performed bulk RNA-sequencing of larval stage 4 (L4) animals depleted of either *hsp-3* or *hsp-4* expression via RNA interference (RNAi). Principal component analysis showed loss of either *hsp-3* or *hsp-4* resulted in distinct and consistent clustering (Fig. 2A). We further confirmed that *hsp-3* or *hsp-4* knockdown did not reduce transcription of the sibling chaperone (Fig. 2B-C). Knockdown of *hsp-3* and *hsp-4* upregulated 83 and 240 distinct genes, respectively, while 126 common genes were upregulated in both siRNA conditions. Loss of HSP-3 downregulated 65 unique genes and knockdown of *hsp-4* downregulated 236 different genes (Fig. 2D-E). 65 genes were downregulated in both conditions, including numerous vitellogenin genes (*vit-1*, *vit-3*, and *vit-4*), which encode for proteins that provide nutrients to embryos during development^46^. A complete list of genes altered by each condition is available in Table S6-8.

**Fig. 2.**
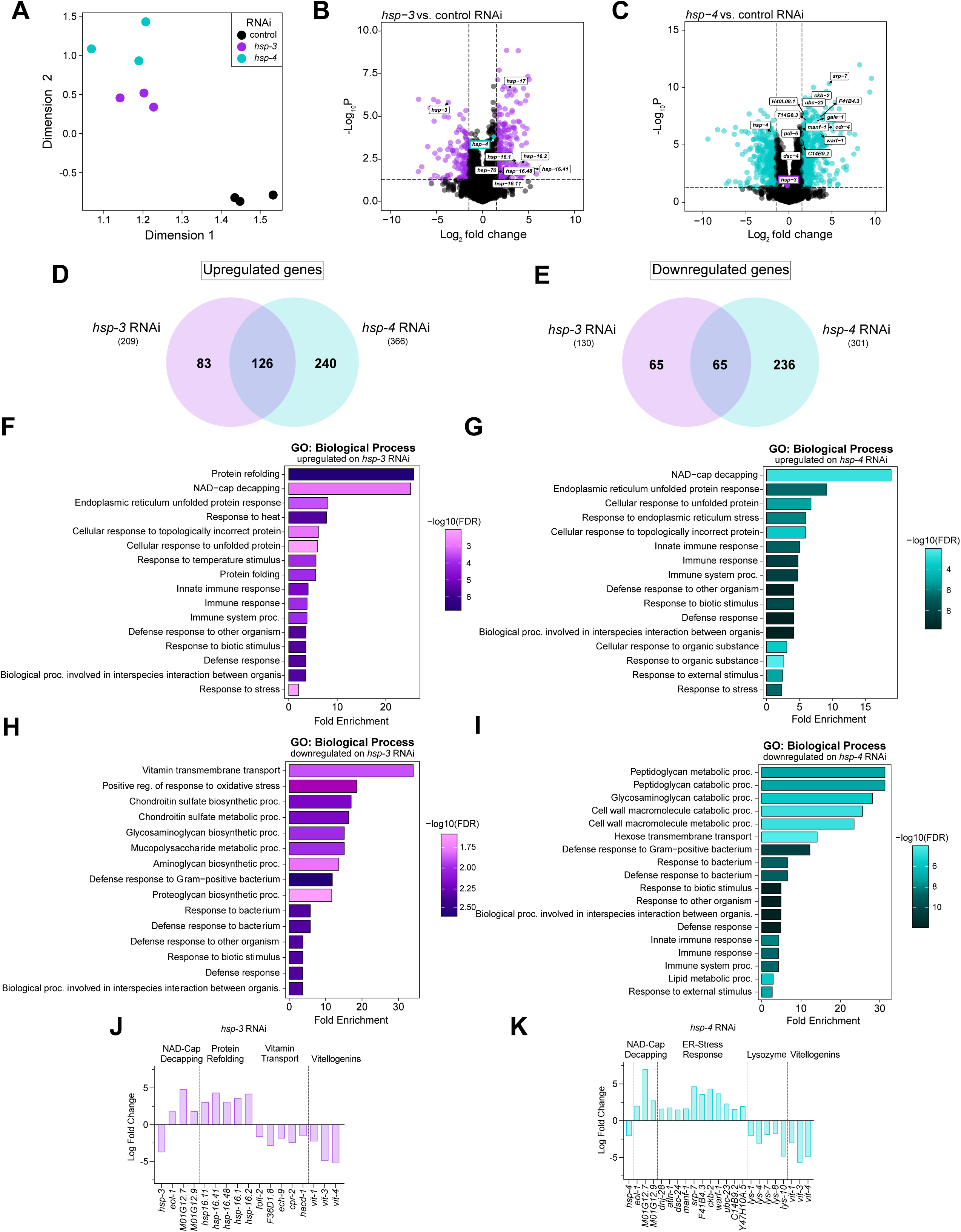
Loss of HSP-3 or HSP-4 triggers distinct transcriptional changes in *C. elegans*. (A). Principal components analysis of L4 WT (N2) worms grown on control (*pos-1*), *hsp-3*, or *hsp-4* siRNA. (B-C). Volcano plots of gene transcripts on *hsp-3* (B) or *hsp-4* (C) siRNA relative to control. (D-E). Venn diagrams depicting the number distinct and overlapping genes upregulated (D) or downregulated (E) on *hsp-3* and/or *hsp-4* siRNA relative to control. (F-G). Gene ontology (GO) analysis of biological processes upregulated on *hsp-3* (F) or *hsp-4* (G) siRNA relative to control. (H-I). Gene ontology (GO) analysis of biological processes downregulated on *hsp-3* (H) or *hsp-4* (I) siRNA relative to control. (J-K). Plot of the relative log-fold change of gene transcripts related to indicated biological processes which are altered on *hsp-3* or *hsp-4* siRNA. For all plots a cutoff of 1.5-fold change was used.

Gene ontology (GO) analysis revealed worms lacking HSP-3 significantly upregulate genes related to protein refolding and chaperone production (e.g., small heat shock proteins like *hsp-16.2*), whereas loss of *hsp-4* increased transcripts related to ER-signaling and cellular responses to stress, including the UPR^ER^ (Fig. 2F-G, J-K). Loss of either *hsp-3* or *hsp-4* also resulted in significant upregulation of genes related to NAD-cap decapping, which regulates RNA transcription^47^(Fig. 2 H-K). Knockdown of *hsp-3* reduced transcripts involved in vitamin transport and various metabolic processes, while loss of *hsp-4* reduced transcripts linked to peptidoglycan metabolism and catabolism— specifically 5 lysozyme genes (*lys-1*, *lys-4*, *lys-7*, *lys-8, and lys-10*)—as well as lipid metabolism. (Fig. 2 H-K). These results suggest HSP-3 and HSP-4 engage in both overlapping and distinct biological pathways. Our data further imply that loss of HSP-4, but not HSP-3, promotes a transcriptional response resembling the UPR^ER^.

### HSP-4 is required for fecundity *and* body growth

Our transcriptomic analysis showed numerous vitellogenin genes are downregulated in both HSP-3*-* and HSP-4*-*deficient animals (Fig. 2J-K). Furthermore, HSP-4::wrmScarlet expression in unstressed animals peaks in the spermatheca (Fig. 1C). We thus hypothesized that HSP3 and HSP-4 are required for proper *C. elegans* reproduction. To test this prediction, we assessed the expression of vitellogenins VIT-1, VIT-2, and VIT-3 using established reporter strains^48^. We observed a reduction in VIT-1 and VIT-3 expression in developing eggs of day 1 adults upon loss of *hsp-3* or *hsp-4*, consistent with the results of our RNA-seq experiments. Co-localization of VIT-1 or VIT-3 with VIT-2 was also severely diminished in developing eggs (Fig. 3A, Fig. S6). In fecundity experiments, worms lacking *hsp-4* laid significantly less eggs than those lacking *hsp-3* or controls throughout early adulthood (Fig. 3B). Animals deficient in *hsp-3* laid less eggs on day 1 but more eggs on day 3 of adulthood compared to controls, indicating a delay in egg laying (Fig. 3B). However, neither HSP-3 nor HSP-4 OE altered egg laying (Fig. 3C-D). These results suggest HSP-3 and HSP-4 are required for proper yolk assembly and affect reproductive aging in *C. elegans*.

**Fig. 3.**
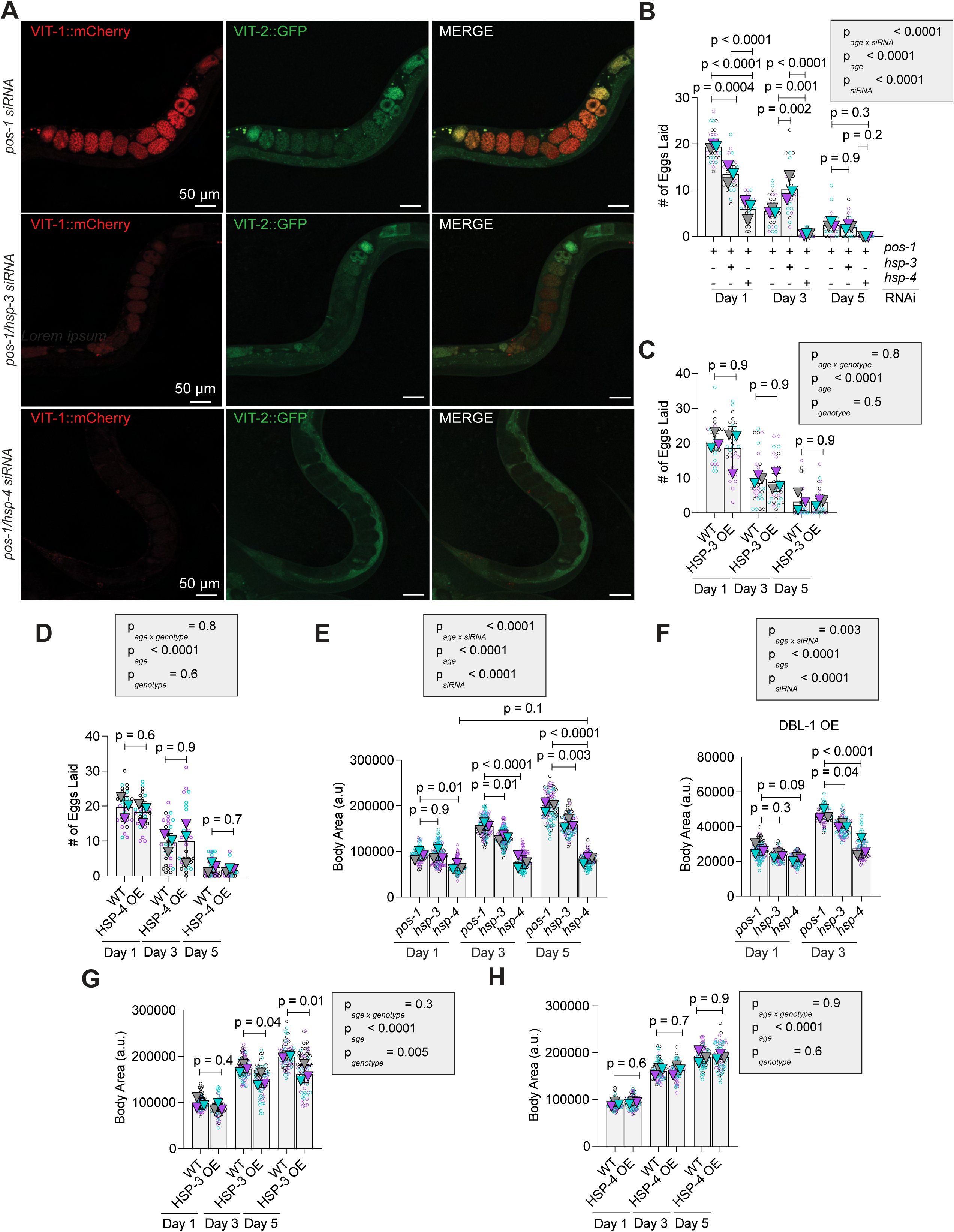
HSP-3 and HSP-4 affect reproductive behavior and body size growth in *C. elegans*. (A). Representative confocal images of developing eggs from MQD2798 (day 1 adults) on indicated siRNA. (B). Timed egg laying (TEL) assay of WT (N2) worms grown on indicated siRNA and day of adulthood. (C). TEL assay of WT and HSP-3 OE (MTX203) worms on *pos-1* siRNA. (D). TEL assay of WT and HSP-4 OE (MTX291) worms on *pos-1* siRNA. For B-D, each circle represents an individual worm, and each triangle represents the average of eggs laid by all worms. (E). Body area quantification of WT worms grown on indicated siRNA and day of adulthood. (F). Body area quantification of DBL-1 OE worms (BW1940) on indicated siRNA and day of adulthood. (G-H). Quantification of WT and HSP-3 OE (G) or HSP-4 OE (H) worms grown on *pos-1* siRNA at indicated days of adulthood. For E-H, each circle represents an individual worm, and each triangle represents the average body size of that given biological replica. *(B-H). Two-way ANOVA followed by Tukey’s multiple comparisons test. p < 0.05 is considered statistically significant*.

Since HSP-4 is required for proper cuticle formation^35^, we next assessed if HSP-3 or HSP-4 are involved in regulating worm body growth. Worms lacking *hsp-3* were similar in size to controls through early adulthood but showed a reduction in body size with age (Fig. 3E). In contrast, worms lacking *hsp-4* failed to increase in size with age (Fig. 3E). DBL-1, the TGF-β ligand required for enlarging body size in *C. elegans*^49^, OE was not sufficient to rescue the reduction in body size of worms lacking *hsp-3* or *hsp-4* (Fig. 3F). HSP-4 OE did not alter body size, but worms overexpressing HSP-3 were smaller compared to controls with age (Fig. 3G-H). These data suggest a proper balance of HSP-3 and HSP-4 is required to regulate body size in *C. elegans* through the engagement of non-canonical body-growth mechanisms.

### Age-and-tissue specific expression of HSP-3 and HSP-4 regulate lifespan

HSP-3 and HSP-4 expression patterns show different tissue distributions and dynamic age-dependent changes (Fig. 2A-C, Fig. S2A-D). We thus tested how either the loss or enhanced expression of either chaperone affects worm lifespan. We found that OE of HSP-3, but not HSP-4, increased lifespan compared to WT controls (Fig. 4A-B, Fig. S4B, 5B). In contrast, life-long depletion of either *hsp-3* or *hsp-4* reduced lifespan compared to control animals; however, worms lacking *hsp-3* were significantly shorter lived compared to worms lacking *hsp-4* (Fig. 4C, Fig. S3B). Neither OE of HSP- 3 in *hsp-4* ablated animals nor OE of HSP-4 in *hsp-3* deficient worms was sufficient to prevent lifespan shortening and in fact further reduced lifespan, indicating neither chaperone can compensate for loss of the other paralog (Figs. 4D-E, Supp. Figs. 4C, 5C).

**Fig. 4.**
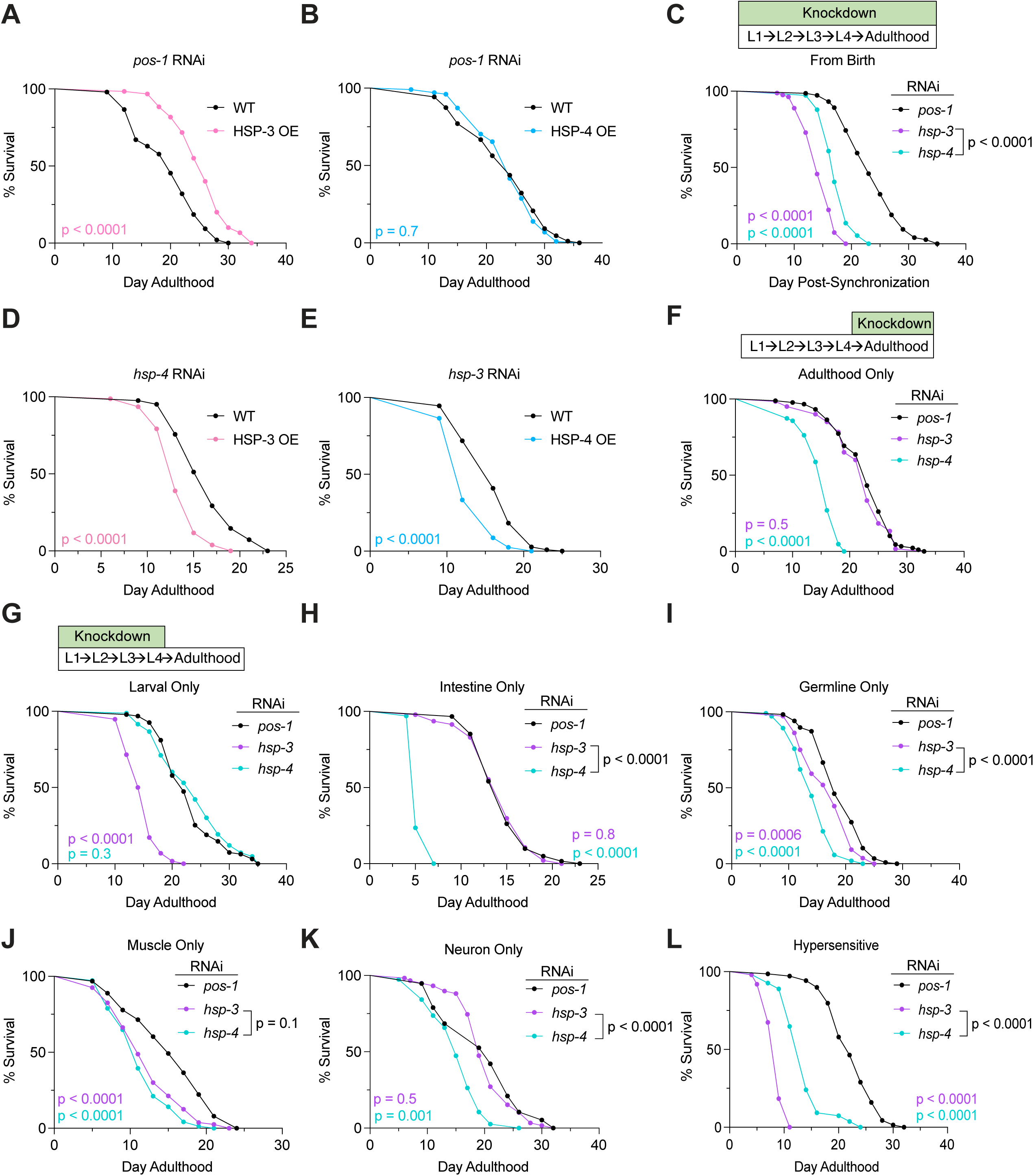
HSP-3 and HSP-4 are required in different tissues and stages of the worm life cycle which contributes to worm lifespan. (A-B). Survival curves of WT (N2) and worms overexpressing HSP-3 (A, MTX203) or HSP-4 (B, MTX291) on *pos-1* siRNA. (C). Survival curve of WT worms grown on *hsp-3* or *hsp-4* siRNA for their entire lives. (D). Survival curve of WT and MTX203 grown on *hsp-4* siRNA. (E). Survival curve of WT and MTX291 grown on *hsp-3* siRNA. (F-H). Survival curves of WT worms grown on *hsp-3* or *hsp-4* siRNA after larval development (F) or only during larval development. (H-K). Survival curves of worms sensitive to siRNA only in the intestines (H, IG1839), germline (I, DCL569), muscle (J, WM118), or neurons (K, MAH677) on indicated siRNA. (L). Survival curve of WT worms hypersensitive to siRNA in neurons on indicated siRNA (L, TU3311). The number of animals and median lifespan data per experiment is in Table S4. *Log-rank Mantel-Cox test. p < 0.05 is considered statistically significant*.

To identify the time(s) of life when HSP-3 and HSP-4 are most important to maintain health, we performed experiments in which *hsp-3* or *hsp-4* expression was only ablated during specific life stages. Knockdown of *hsp-4*, but not *hsp-3*, during adulthood shortened worm lifespan akin to life-long *hsp-4* ablation (Fig. 4F, Fig. S3C). Conversely, *hsp-3* loss only during larval development was sufficient to reduce lifespan, whereas knockdown of *hsp-4* during the larval stages did not affect worm longevity (Fig. 4G, Fig. S3D). RT-qPCR analysis confirmed that worms treated with *hsp-3* siRNA only during development had regained normal *hsp-3* transcript levels within 48 hours after completion of RNAi treatment (Fig. S3E).

Next, we sought to define which tissues are most dependent on HSP-3 and HSP- 4 expression. Using established worm strains that enable tissue-restricted RNAi-mediated gene knockdown^50,51,52^, we found that life-long loss of *hsp-3* in the intestine did not shorten lifespan, but intestine-specific loss of *hsp-4* lead to matricide and early worm death (Fig. 4H, Fig. S7C). Animals with *hsp-3* or *hsp-4* ablation in the germline only were short lived, with *hsp-4* deficient worms showing a significantly more severe lifespan shortening than those lacking HSP-3 (Fig. 4I, Fig. S7D). Muscle-restricted knockdown of *hsp-3* and *hsp-4* decreased lifespan compared to controls to a similar extent (Fig. 4J, Fig. S7E). Animals with neuron-specific loss of *hsp-4* were short lived, but lifespan of worms with pan-neuronal loss of *hsp-3* was similar to controls (Fig. 4K, Fig. S7B). However, knockdown of *hsp*-*3* or *hsp-4* in worms with neurons hypersensitized to RNAi treatment in a WT (i.e. RNAi-sensitive) background^53^ showed similar lifespan patterns as WT (Fig. 4L, Fig. S7A).

Taken as a whole, our results show that HSP-3 loss is most impactful during development but tolerated in adulthood whereas HSP-4 ablation in adult worms, even when restricted to specific tissues, significantly decreases worm health.

### Loss of HSP-4 protects against a***β***_1-42_-associated paralysis

Given the roles of HSP70 chaperones in maintaining proteostasis, we next tested if HSP-3 or HSP-4 were required to buffer against protein aggregation-induced ER stress. As previous work showed *hsp-3* and *hsp-4* transcripts are upregulated in animals expressing human amyloid beta (aβ_1-42_)in the body wall muscle^54,55^, we assessed aβ_1-42_-mediated paralysis in HSP-3- and HSP-4-deficient worms at aggregation-permissive (20 °C, “non-inducive”) and inducive (25 °C, “inducive”) temperatures. At permissive temperatures, *hsp-3* and *hsp-4* loss significantly increased the rate of paralysis in aβ_1-42_-expressing worms compared to controls (Fig. 5A, Fig. S8A). Consistent with trends observed in lifespan assays (Fig. 4C), loss of *hsp-3* increased the rate of worm paralysis compared to worms lacking *hsp-4* (Fig. 5A, Fig. S8A). These data suggest loss of both HSP-3 and HSP-4 disrupt proteostasis and compound exogenous protein misfolding stress.

**Fig. 5.**
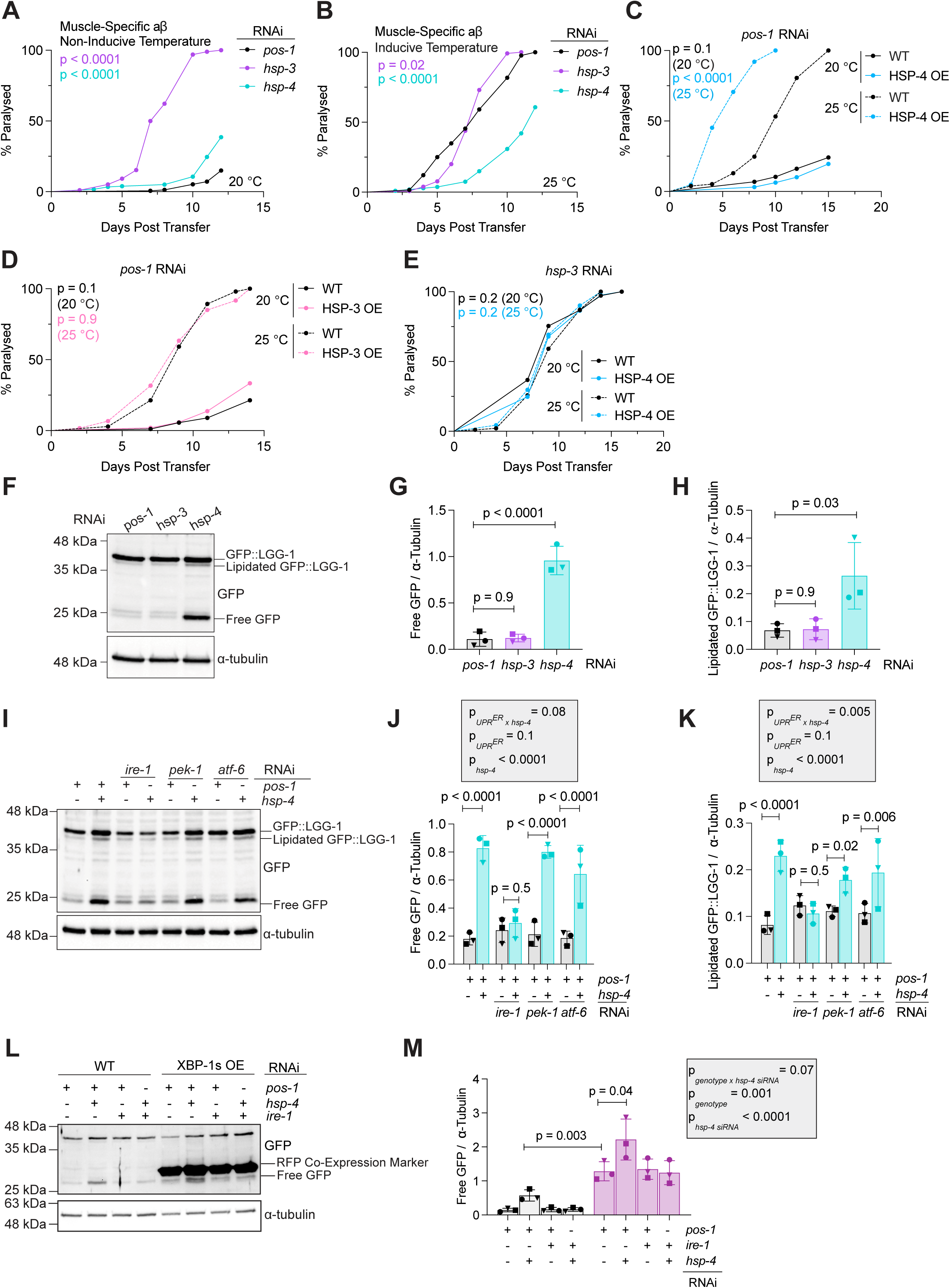
Loss of HSP-4 delays aβ*_1-42_-induced paralysis and triggers an increase in autophagy through IRE-1 independent of spliced XBP-1 signaling.* (A-B). Paralysis assay of GMC101 on indicated siRNA at the non-inducive (A, 20 °C) and inducive (B, 25 °C) temperatures. (C). Paralysis assay of GMC101 and MTX300 (GMC101; MTX291) on *pos-1* siRNA. (D). Paralysis assays of GMC101 and MTX328 (GMC101; MTX203) on *pos-1* siRNA. (E). Paralysis assay of GMC101 and MTX300 on *hsp-3* siRNA. (F-K). Representative western blot (F, I) and quantification (G-H, J-K) of GFP::LGG-1 expressing worms (DA2123) grown on indicated siRNA. (L-M). Representative western blot (L) and processed GFP quantification (M) of DA2123 (“WT”) and MTX339 (DA2123; AGD928 “XBP-1s OE”) on indicated siRNA. For F-M, worms were collected as day 1 adults. Membranes were probed with an anti-GFP (Roche, Cat #11814460001) and anti-alpha tubulin antibody (DSHB, Cat #12G10). The number of animals and median paralysis data per experiment is in Table S5. *(A-E). Log-rank Mantel-Cox test. (G-H). One-way ANOVA followed by Tukey’s multiple comparisons test. (J-K, M). Two-way ANOVA followed by Tukey’s multiple comparisons test. p < 0.05 is considered statistically significant*.

Worms lacking *hsp-3* showed similar rates of paralysis at both the inducive and non-inducive temperatures (Fig. S8A). However, loss of *hsp-4* at the inducive temperature reduced the rate of paralysis compared to controls (Fig. 5B, Fig. S8A). In agreement with these results, HSP-4 OE significantly increased paralysis of aβ_1-42_-expressing worms (Fig. 5C, Fig. S9A). However, HSP-3 OE did not alter the rate of paralysis rate compared to controls (Fig. 5D, Fig. S9B). HSP-4 OE was unable to compensate for the increase in paralysis caused by loss of *hsp-3* (Fig. 5E, Fig. S9C). Altogether, these data suggest HSP-3, but not HSP-4, is required to buffer against aβ_1-42_ aggregation-associated proteotoxic stress.

### Loss of HSP-4 increases autophagy through engagement of IRE-1 but is independent of XBP-1 splicing

The finding that the loss of an ER-chaperone (HSP-4) improved tolerance to cytosolic aβ_1-42_ aggregation was unexpected. We hypothesized that loss of HSP-4 triggers an increase in cytosolic protein turnover through an ER-dependent protein degradation mechanism, such as ER-selective autophagy or recovER-Phagy, accompanied by an increase in autophagy.

In the cytosol, autophagy maintains proteostasis by delivering cargo in autophagosomes to lysosomes for protein degradation and recycling^56^. Autophagosome formation requires cytosolic ATG8/LC3, which is recruited to and subsequently incorporated into autophagosome membranes (i.e. lipidated); upon fusion with the lysosome, contents of the newly-formed autolysosome are then degraded^57^. In *C. elegans*, relative autophagic activity (autophagic flux) can be assessed by probing lysate of worms expressing a GFP::LGG-1 (LC3 ortholog) fusion protein via immunoblotting. This method allows for delineation of the total abundance of GFP::LGG-1, lipidated GFP::LGG-1 (reflective of total autophagosomes), and processed GFP (reflective of autolysosomal degradation)^58,59^.

To test our hypothesis, we knocked down *hsp-3* or *hsp-4* in worms expressing GFP::LGG-1^60^. We observed loss of *hsp-3* did not affect GFP::LGG-1 processing or lipidation; however, *hsp-4* knockdown significantly increased the amount of both lipidated and processed GFP::LGG-1 (Fig. 5F-H, Fig. S10A). These results suggest that loss of HSP-4, but not HSP-3, increases autophagy in *C. elegans*.

To test if the upregulation of autophagy upon HSP-4 loss was mediated through the UPR^ER^, we simultaneously knocked down *hsp-4* in combination with *ire-1*, *pek-1*, or *atf-6* in GFP::LGG-1 expressing animals. We observed that *ire-1*;*hsp-4* double-knockdown prevented increased GFP::LGG-1 lipidation and processing compared to controls, whereas the simultaneous knockdown of *pek-1* or *atf-6* with *hsp-4* still resulted in increased autophagy (Fig. 5I-K, Fig. S10B). Using RT-qPCR, we confirmed *hsp-4* knockdown efficiency was unaffected by combinations of siRNA in all conditions (Fig. S10C-E).

To determine if canonical IRE-1 signaling was sufficient to increase autophagy, we overexpressed spliced XBP-1 in GFP::LGG-1 animals using established alleles^61^. We found that spliced XBP-1 OE increased autophagy compared to WT controls, but *hsp-4* knockdown still boosted GFP::LGG-1 processing compared to control siRNA in spliced XBP-1 OE animals (Fig. 5L-M, Fig. S11A). These results suggest the upregulation of autophagy upon HSP-4 ablation requires IRE-1 but is at least partially independent of canonical IRE-1/XBP-1 signaling.

### Loss of HSP-4 triggers increased autophagy through the ER-Phagy receptor, C18E9.2

Given that IRE-1, but not spliced XBP-1 signaling, was required to upregulate autophagy upon *hsp-4* knockdown, we next tested if loss of HSP-4 triggered an increase in autophagy via ER-Phagy. As HSP-4 is significantly upregulated during times of ER stress and ER expansion, we hypothesized HSP-4 could inhibit ER-phagy to allow for stress-induced ER-expansion. Thus, loss of HSP-4 could relieve this repression and consequently enhance ER-phagy.

In human cells, the ER-phagy receptor, Sec-62, is required to reduce the size of the ER after stress resolution in a specific form of ER-phagy called recovER-Phagy^62^. We thus assessed if the *C. elegans* Sec-62 ortholog, C18E9.2, is required to upregulate autophagy triggered by HSP-4 loss. Double knockdown of *C18E9.2* and *hsp-4* inhibited the upregulation of GFP::LGG-1 lipidation and processing (Fig. 6A-C, Fig. S11B). *hsp-4* knockdown efficacy was unaffected by combinatory knockdown with *C18E9.2* siRNA, though knockdown of *C18E9.2* slightly but significantly increased HSP-4 protein and transcript abundance compared to controls; this is consistent with findings in human cells showing that Sec-62 is required to reduce BiP levels following ER-stress^26^ (Fig. S11C-F).

**Fig. 6.**
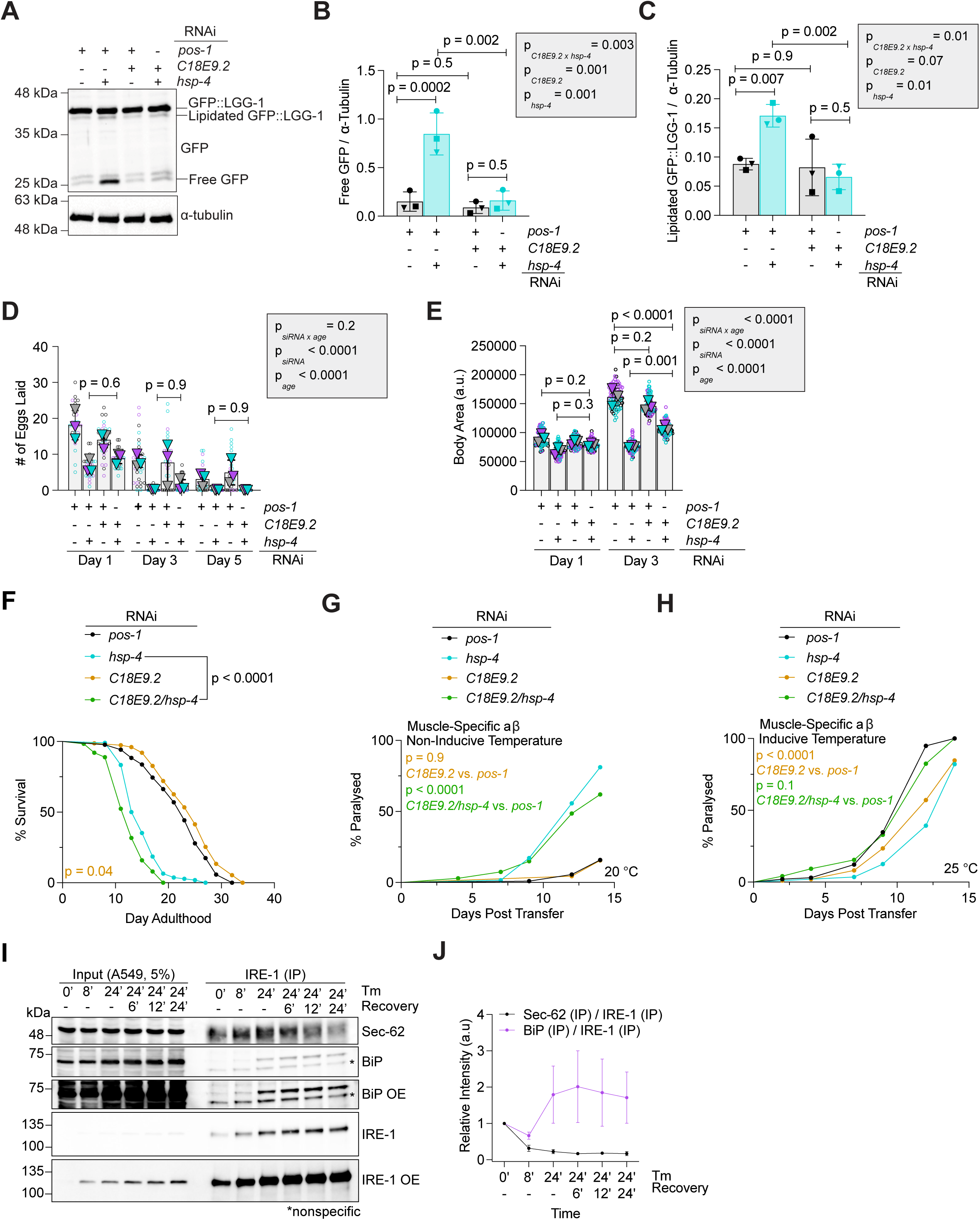
**BiP/*HSP-4 and IRE-1 regulate ER-Phagy through Sec-62/C18E9.2*** (A-C). Representative western blot (A) and quantification (B-C) of DA2123 grown on indicated siRNA. Worms were collected as Day 1 adults and probed with an anti-GFP (Roche, Cat #11814460001) and anti-alpha tubulin antibody (DSHB, Cat #12G10). (D). TEL assay of WT (N2) worms grown on indicated siRNA and days of adulthood. (E). Body area quantification of WT worms grown on indicated siRNA and day of adulthood. (F). Survival curve of WT worms grown on indicated siRNA. (G-H). Paralysis assay of GMC101 grown on the indicated siRNA at the non-inducive (G) and inducive (H) temperatures. (I-J). Representative western blots (I) and relative quantification (J) of co-immunoprecipitation assay of Tm-treated A549 cells pulling on and probed for IRE-1 (Cell Signaling, Cat #3294S; Rabbit IgG, Cell Signaling, Cat #7074S), Sec-62 (Abcam, Cat #EPR9212; Light Chain-Specific Rabbit IgG (Cell Signaling, Cat #D4W3E), and BiP (Proteintech, Cat #66574; Mouse IgG, Cell Signaling Cat #7076S). For F-H, the number of animals and median lifespan/paralysis data per experiment is in Tables S4/S5. *(B-E). Two-way ANOVA followed by Tukey’s multiple comparisons test. (F-G). Log-rank Mantel-Cox test. p < 0.05 is considered statistically significant*.

We then assessed the roles of the other three conserved ER-Phagy receptors (RET-1/RTN3, CDKR-3/CDK5, and ATLN-1/ATL3) in the same assay to determine if other ER-Phagy receptors may be involved in this upregulation. Combinatorial knockdown of *hsp-4* with *ret-1* or *cdkr-3* did not prevent HSP-4-loss-induced upregulation of LGG-1 processing and lipidation (Fig. S12A-F). Furthermore, upon *hsp-4* knockdown, *atln-1* KO animals showed a similar upregulation of GFP::LGG-1 processing compared to WT controls but reduced GFP::LGG-1 lipidation (Fig. S12G-I). Altogether, these results suggest the primary upregulation of autophagy triggered by *hsp-4* knockdown is mediated through C18E9.2.

Given C18E9.2 lacks a luminal HSP70-binding J-domain, a direct interaction of HSP-4 with C18E9.2 is unlikely. We thus asked if HSP-4’s regulation of C18E9.2 was mediated by HSP-4 potentially binding to the J-domain containing Sec-63 ortholog and Sec-62/C18E9.2 binding partner, DNJ-29. In human cells, BiP binds to the J-domain of Sec-63 to assist in protein refolding through the Sec-61/Sec-2/Sec-63 protein translocation complex^63–66^. Interestingly, worms deficient in HSP-4 and DNJ-29 still exhibited increased autophagy (Fig. S12J-L).

Taken as a whole, our work supports a model in which HSP-4, but not HSP-3, regulates ER-cytosolic crosstalk and autophagy through engagement of the ER-Phagy receptor C18E9.2/Sec-62 and IRE-1, but is independent of canonical IRE-1 signaling.

### C18E9.2-mediated ER phagy contributes to some, but not all, phenotypes associated with HSP-4 loss

Having established direct links between HSP-4, C18E9.2, and ER-phagy, we repeated the previously introduced fecundity, body size, lifespan, and paralysis assays with worms deficient in *C18E9.2* to determine if C18E9.2 contributed to phenotypes mediated by loss of HSP-4. Knockdown of *C18E9.2* was not sufficient to rescue the egg laying deficiency of *hsp-4* knockdown worms (Fig. 6D). However, worms lacking both *C18E9.2* and *hsp-4* were significantly larger than worms lacking *hsp-4* alone (Fig. 6E). Interestingly, loss of *C18E9.2* alone slightly extended lifespan, but animals with knockdown of *hsp-4* and *C18E9.2* together were shorter lived than those lacking HSP-4 and controls (Fig. 6F, Fig. S13A).

Double knockdown of *hsp-4* and *C18E9.2* did not reduce the rate of paralysis of aβ_1-42_-expressing worms compared to *hsp-4* knockdown alone at the non-inducive temperature (Fig. 6G, Fig. S13B). *C18E9.2* knockdown alone reduced paralysis compared to controls at the inducive temperature; however, double knockdown of *C18E9.2* and *hsp-4* at the inducive temperature increased the rate of paralysis back to that of controls (Fig. 6H, Fig. S13B).

These data suggest that HSP-4 loss increases C18E9.2-mediated ER-Phagy, thus reducing aβ_1-42_-dependent paralysis and limiting body growth. However, the observed decreases in fecundity and lifespan upon HSP-4 ablation are not linked to C18E9.2-dependent ER-phagy.

### ER-stress increases interaction of IRE-1 and Sec-62 in human cells

Given our worm data suggesting a relationship between worm orthologs of BiP, Sec-62, and IRE-1 independent of spliced XBP-1 signaling, we hypothesized that in mammalian cells a fraction of IRE-1 binds to Sec-62 to limit ER-phagy when BiP is released from IRE-1 during ER stress. This model further predicts that BiP binding to IRE-1 would limit the interaction of Sec-62 and IRE-1. To test this, we stressed A549 cells for 24 hours using tunicamycin followed by a 24 hour recovery period, then immunoprecipitated IRE-1. In line with our hypothesis, Sec-62 co-immunoprecipitated with IRE-1; however, as co-immunoprecipitation of BiP with IRE-1 increased, pulldown of Sec-62 was reduced (Fig. 6I-J, Fig. S14). Combined, these data suggest a conserved model in which HSP-4/BiP’s regulation of IRE-1 limits the direct interaction of IRE-1 and C18E9.2/Sec-62. (Fig. 7). Cumulatively, these data highlight the importance of understanding the roles of individual paralogs present in model organisms to best interpret the conserved functions of these proteins in more complex organisms.

**Fig. 7.**
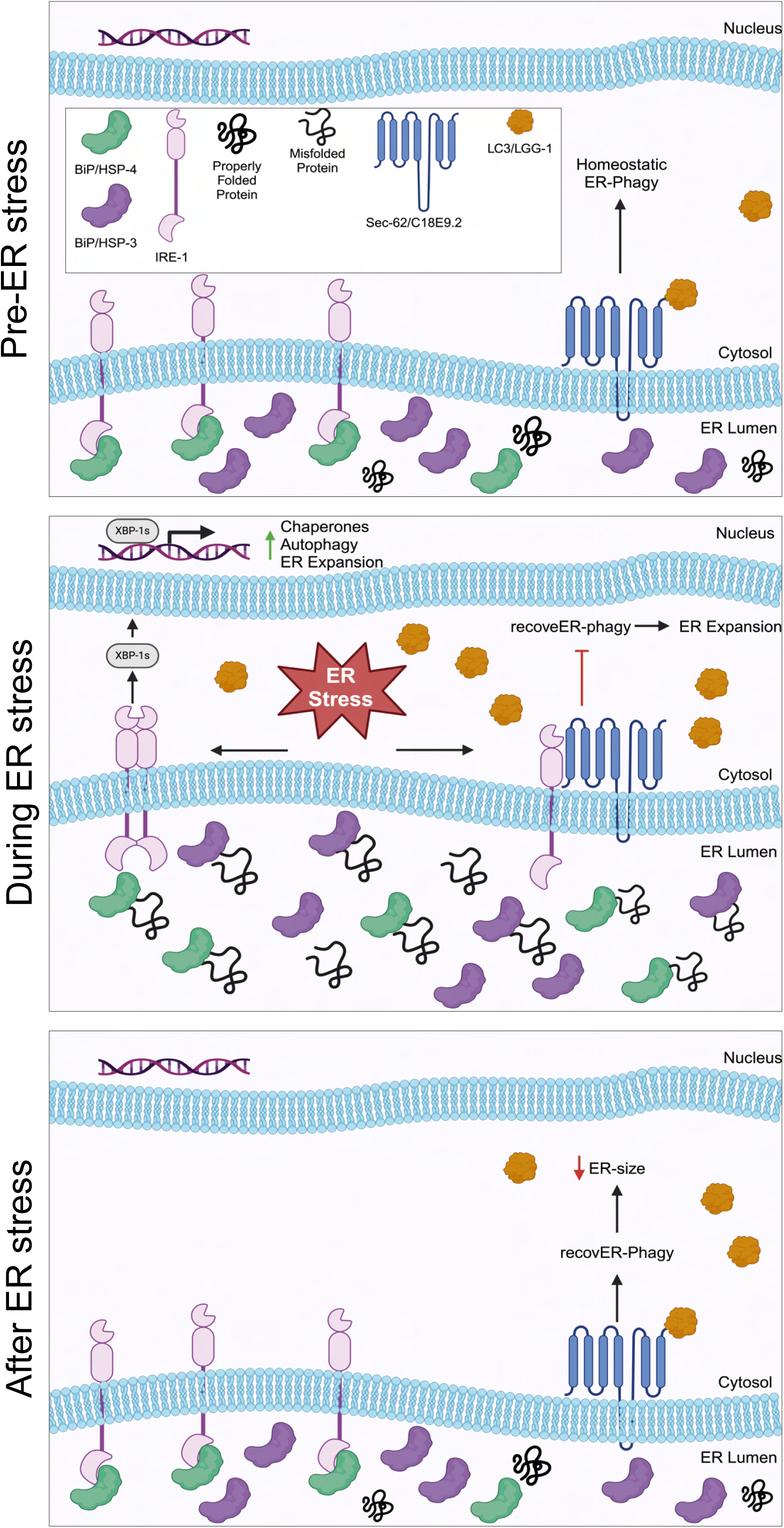
Working model as to how BiP and IRE-1 regulate Sec-62-mediated recovER-Phagy.

## Discussion

In this study, we define how the two *C. elegans*’ BiP orthologs, HSP-3 and HSP- 4, functionally overlap and diverge to maximize ER stress resilience and contribute to worm physiology. We define distinctive roles for both paralogs in ER and non-ER stress resistance, reproduction, body size growth, lifespan, and ER-Phagy regulation. Our results further describe a novel conserved mechanism linking HSP-4/BiP-mediated regulation of IRE-1 to C18E9.2/Sec-62-associated recovER-Phagy.

Under stress conditions, UPR^ER^ induction leads to the adaptive expansion of the ER lumen^67^. Once the stress is resolved, recovER-phagy, a select form of ER-phagy requiring Sec-62, removes excessive ER membrane and luminal content. This process decreases the abundance of BiP and other stress-induced ER proteins to pre-stress levels^26^. Our findings suggest a direct relationship between IRE-1, Sec-62, and ER stress, supporting a model in which BiP/HSP-4’s de-repression of IRE-1 allows for a direct interaction between Sec-62/C18E9.2 and IRE-1 in times of ER stress. We postulate this interaction leads to ER-Phagy inhibition primarily to allow for ER expansion, and the repression of IRE-1 upon BiP binding following ER stress resolution allows for Sec-62/C18E9.2-medidated recovER-Phagy (Figs. 6 and 7).

This model contrasts previously described mechanisms of how BiP regulates other known ER-Phagy receptors. For example, ER-stress induced binding of BiP to FAM134B promotes ER-Phagy and cell survival in breast cancer and cardiac cells^31,32^. Thus, our data suggest a novel mechanism involving the upstream regulation of the UPR^ER^ and Sec-62-mediated ER-Phagy—adding to BiP’s multi-faceted repertoire of ER homeostasis regulation.

Previous work showed a transcriptional relationship between *hsp-3* and *hsp-4*, implying compensatory regulation between the two proteins^34^. However, studies by *Zha et al.* indicated HSP-4, but not HSP-3, is required for proper cuticle functionality in the worm, suggesting the possibility of functional diversity in *C. elegans*^35^. Our work builds upon these findings, showing HSP-3 and HSP-4 are unable to compensate for loss of the other protein in lifespan and protein misfolding stress resistance assays (Fig. 4D-E, Fig. 5E). These OE experiments further imply HSP-3 and HSP-4 may have partially opposing functions. Furthermore, our temporal and tissue-specific analysis of HSP-3 and HSP-4 in the context of lifespan (Fig. 4F-L) and clear tissue-and age-specific differences in protein expression (Fig. 2, Fig. S2) suggest a greater diversification of HSP-3 and HSP-4 function within individual tissues at distinct life stages. We thus conclude that HSP-3 and HSP-4 evolved divergent roles supporting distinct aspects of the worm’s life cycle. Our RNA-sequencing (Fig. 2), ER-stress resistance (Fig. 1), and ER-Phagy regulation data (Figs. 5-6) are consistent with a model in which HSP-3 represents the canonical ER-resident HSP70 chaperone responsible for *de novo* and misfolded protein (re)folding, while HSP-4 is primarily engaged in orchestrating interorganellar signaling and ER stress mitigation pathways (Fig. 7).

This division of chaperone function is supported by the findings that HSP-4 OE increases paralysis of aβ_1-42_-expressing worms, whereas HSP-3 OE has no effect (Figs. 5C-D). It is plausible that ER-restricted HSP-3 lacks direct access to cytosolically expressed aβ_1-42_, whereas ER-mediated proteotoxic stress response mechanisms which temporarily alleviate proteotoxic burden, such as ER-Phagy, may be limited by the excess presence of HSP-4. This notion of compartmentalized function is further supported by our lifespan data. Molting events during the larval stages require significant protein production, which dramatically increases demand on protein folding and quality control systems such as the UPR^ER^ ^68^. Thus, elevating HSP-3 levels while restricting HSP-4 levels during development (Fig. 1A-C) would allow for protein quality control mechanisms to remain active and functional. Conversely, losing HSP-3 during this stage of life may reduce quality protein production and maintenance which has longstanding effects on worm health.

Furthermore, our tissue-specific (Fig. 4H-L) and adulthood-restricted (Fig. 4F) knockdown experiments suggest there are tissue and aging-specific demands which may require a switch in protein quality control mechanisms from direct chaperone-mediated protein (re)folding to broader transcriptional regulation with increased age and in specific tissues. However, our understanding of the tissue-and-age-specific mechanism(s) by which HSP-3 and HSP-4 regulate proteostasis is incompletely understood.

Taken as a whole, our work identifies both redundancies and functional diversifications of HSP-3 and HSP-4 in *C. elegans* and describes a novel role for BiP-mediated regulation of ER-proteostasis (i.e., Sec-62-mediated recovER-Phagy).

### Study Limitations

Our approach to assessing autophagy in a total worm lysate, while efficient, reproducible, and high-throughput, prevents us from identifying tissue-specific contributions driving changes in autophagy. Furthermore, the identification of total autophagosomes can be particularly difficult to interpret if other florescent proteins are present (e.g. Fig. 5 L-M, Fig. S12) which do not allow for overexposure of the membranes. Finally, to the best of our knowledge, a chemical or genetic approach to stimulate autophagy in *C. elegans* has yet to be described, which prevented us to determine if increased autophagy is sufficient to increase resistance to aβ_1-42_-induced paralysis.

## Acknowledgments

We thank the members of the Truttmann lab for their helpful comments and discussions. We would like to thank SUNY Biotech for generation of numerous *C. elegans* strains. Thank you to the lab of Dr. Pierre Coloumbe for access to their Zeiss LSM 800 confocal laser scanning microscope. And finally, thank you to the lab of Dr. Andrew Dillin for providing us strain AGD925. Dr. William Giblin is acknowledged for his assistance in integrating numerous *C. elegans* strains. NDU was supported by Training Grants GM008322-30 and AG000114-37, as well as award 1F31AG085891-01A1. MCT is supported by an Alzheimer’s foundation Young Investigator Award, a Ruth K Broad foundation award and grant 1R35GM142561.

## Author contributions

MCT supervised the project. NDU and MCT planned and designed all experiments. NDU, SML, KVP, BA, AK, ST, MCT, and ZM performed all experiments. NDU wrote the first draft of the manuscript and subsequent drafts were edited by MCT and NDU.

## Data availability statement

The unprocessed raw datasets (images) generated and analyzed during the current study are available from the corresponding author upon reasonable request.

## Additional information

The authors declare no competing interests.

**Fig. S1. HSP-3::wrmScarlet and HSP-4::wrmScarlet proteins are functional, transcribed like wild type proteins, and have distinct temporal protein distributions.** (A) Development assays of WT and reporter strains grown on indicated siRNA. (B-C). RT-qPCR of adult N2 and PHX4377 (B) or PHX4415 (C) worms for indicated gene relative to control gene (*cdc42*). Worms were grown on *pos-1* siRNA and collected 72 hours after synchronization at 20 °C (n = 3). (D) Three biological replicates of western blots of worms expressing HSP-3::wrmScarlet (PHX4377) or HSP- 4::wrmScarlet (PHX4415) grown on *pos-1* siRNA at 20 °C and collected 24’, 48’, and 72’ after synchronization. Membranes were probed with an anti-RFP (Chromotek, Cat #6g6) and anti-alpha tubulin (DSHB, Cat #12G10) antibody. Red box indicates what is shown in main text *(A) Two-way ANOVA followed by Tukey’s multiple comparisons test. (B-C). Student’s t-test. p < 0.05 is considered statistically significant*.

**Fig. S2. *HSP-3::wrmScarlet and HSP-4::wrmScarlet show distinct age and thapsigargin-induced expression responses.*** (A-D). Representative images (A-B) and quantification (C-D) of PHX4377 and PHX4415 grown on *pos-1* siRNA at denoted ages. (E-F). Representative images (E) and quantification (F) of PHX4337 grown on indicated siRNA and treated with 0.1 µg/mL thapsigargin (Tg) or equivalent volume of solvent (DMSO) as day one adults for 24 hours. (G-H). Representative images (G) and quantification (H) of PHX4415 grown on indicated siRNA and treated with 0.1 µg/mL Tg or DMSO as day one adults for 24 hours. Images were taken using a Keyonce BZ-X700 fluorescent microscope at 10x magnification. *(B,D). Ordinary one-way ANOVA followed by Tukey’s multiple comparisons test. (F,H). Two-way ANOVA followed by Tukey’s multiple comparisons test. p < 0.05 is considered statistically significant*.

**Fig. S3. HSP-3 and HSP-4 have distinct effects on prolonged ER stress resistance lifespan in WT worms.** (A). Three biological replicas of survival curves of WT (N2) worms treated with 10 µg/mL of Tm or equivalent volume of M9 as day one adults grown on indicated siRNA. (B). Survival curves of three biological replicas of WT worms synchronized on *pos-1*, *hsp-3,* or *hsp-4* siRNA. (C) Survival curves of three biological replicas of WT worms synchronized on *pos-1* siRNA then transferred to *hsp-3* or *hsp-4* siRNA 72 hours after development. (D). Survival curves of three biological replicas of WT worms synchronized on *pos-1*, *hsp-3*, or *hsp-4* and transferred to pos-1 siRNA 72 hours after development. (E). RT-qPCR of *hsp-3* transcript levels treated with indicated siRNA relative to control (*cdc42*). Worms were grown on *pos-1* or *hsp-3* siRNA for 72 hours at 20 °C prior to being transferred. The number of animals and median lifespan data per experiment is in Table S4. *(A-D). Log-rank Mantel-Cox test. (E). Ordinary one-way ANOVA followed by Dunnett’s multiple comparisons test. p < 0.05 is considered statistically significant*.

**Fig. S4. *HSP-3 overexpression increases lifespan and improves ER-stress resistance but cannot compensate for loss of HSP-4.*** (A). Survival curves of three biological replicas of WT (N2) and HSP-3 overexpresing (MTX203) worms treated with 10 µg/mL of Tm or solvent (M9) control. (B). Survival curves of three biological replicas of N2 and MTX203 on *pos-1* siRNA. (C). Survival curves of three biological replicas of N2 and MTX203 on *hsp-4* siRNA. The number of animals and median lifespan data per experiment is in Table S4. *Log-rank Mantel-Cox test. p < 0.05 is considered statistically significant*.

**Fig. S5. *HSP-4 overexpression does not affect lifespan, ER-stress resistance, or compensate for loss of HSP-3.*** (A). Survival curves of three biological replicas of WT (N2) and HSP-4 overexpressing (MTX291) worms treated with 10 µg/mL Tm or solvent control. (B). Survival curves of three biological replicas of N2 and MTX291 grown on *pos-1* siRNA. (C). Survival curves of three biological replicas of N2 and MTX291 grown on *hsp-3* siRNA. The number of animals and median lifespan data per experiment is in Table S4. *Log-rank Mantel-Cox test. p < 0.05 is considered statistically significant*.

**Fig. S6. Loss of HSP-3 and HSP-4 reduces vitellogenin protein levels in developing eggs.** (A) Representative confocal images of MQD2775 on indicated siRNA.

**Fig. S7. *Tissue-specific loss of HSP-3 or HSP-4 have diverse effects on worm lifespan.*** (A-E). Survival curves of three biological replicas of TU3311 (A), MAH677 (B), IG1839 (C), DCL569 (D), and WM118 (E) grown on *pos-1*, *hsp-3*, or *hsp-4* siRNA. The number of animals and median lifespan data per experiment is in Table S4. *Log-rank Mantel-Cox test. p < 0.05 is considered statistically significant*.

**Fig. S8. *HSP-4 knockdown reduces paralysis of GMC101.*** (A). Three biological replicas of paralysis assays of GMC101 grown on indicated siRNA. Worms were transferred to the inducive temperature as day 1 adults. The number of animals and median paralysis data per experiment is in Table S5. *Log-rank Mantel-Cox test. p < 0.05 is considered statistically significant*.

**Fig. S9. *HSP-4 OE increases paralysis of GMC101, but HSP-3 OE does not affect paralysis.*** (A). Three biological replicas of paralysis assays of GMC101 and MTX300 (GMC101; MTX291) on *pos-1* siRNA. (B). Three biological replicas of paralysis assays of GMC101 and MTX328 (GMC101; MTX203) on *pos-1* siRNA. (C). Three biological replicas of paralysis assays of GMC101 and MTX300 on *hsp-3* siRNA. The number of animals and median paralysis data per experiment is in Table S5. *Log-rank Mantel-Cox test. p < 0.05 is considered statistically significant*.

**Fig. S10. *Loss of* HSP-4 *increases autophagy and requires IRE-1.*** (A-B). Three biological replicas of western blots of DA2123 grown on indicated siRNA. Membranes were probed with an anti-GFP (Roche, Cat #11814460001) and alpha tubulin antibody (DSHB, Cat #12G10). (C-E). RT-qPCR analysis of WT worms grown on indicated siRNA for *hsp-4* transcript relative to *cdc42*. Red box indicates what is shown in main text. *Ordinary one-way ANOVA followed by Tukey’s multiple comparisons test. p < 0.05 is considered statistically significant*.

**Fig. S11. Loss of *hsp-4* increases autophagy independent of IRE-1 signaling but requires ER-Phagy receptor, C18E9.2.** (A-B). Western blots (A) and quantification (B)) from three biological replicas of DA2123 (“WT”) and MTX339 (DA2123; AGD925 (“xbp-1s OE”)) grown on indicated siRNA. (C). Three biological replicas of DA2123 grown on indicated siRNA. Membranes were probed with an anti-GFP (Roche, Cat #11814460001) and alpha tubulin antibody (DSHB, Cat #12G10). (D). RT-qPCR of *C18E9.2* transcript of WT worms on indicated siRNA relative to *cdc42*. (E). Quantification of wrmScarlet intensity from PHX4415 grown on indicated siRNA. (F). RT-qPCR of *hsp-4* transcript of WT worms on indicated siRNA relative to *cdc42*. Red box indicates what is shown in main text. *(C-E). Ordinary one-way ANOVA followed by Tukey’s multiple comparisons test. p < 0.05 is considered statistically significant*.

**Fig. S12. *Other conserved ER-Phagy receptors or DNJ-29 are not required for upregulation of autophagy triggered by* hsp-4 *knockdown.*** (A-L). Western blots (A, D, G, J) and quantification (B-C, E-F, H-I, K-L) from three biological replicas of DA2123 grown on indicated siRNA or genetic background probed with indicated antibodies. For G-I, “WT” indicates DA2123 and *“atln-1* KO” indicates MTX341 (RB1127; DA2123). Membranes were probed with an anti-GFP (Roche, Cat #11814460001) and alpha tubulin antibody (DSHB, Cat #12G10). (M-O). RT-qPCR analysis of indicated gene grown on indicated siRNA relative to *cdc42*. *(B-C, E-F, H-I, K-L). Two-way ANOVA followed by Tukey’s multiple comparisons test. (M-O). One-way ANOVA followed by Tukey’s multiple comparisons test. p < 0.05 is considered statistically significant*.

**Fig. S13. Loss of C18E9.2 does not rescue lifespan reduction caused by loss of HSP-4 but is required to delay paralysis in GMC101 lacking HSP-4.** (A) Three biological replicas of lifespan assays of WT worms grown on indicated siRNA (B). Three biological replicas of paralysis assays of GMC101 on indicated siRNA. The number of animals and median lifespan/paralysis data per experiment is in Tables S4/S5. *Log-rank Mantel-Cox test. p < 0.05 is considered statistically significant*.

**Fig. S14. *ER stress increases IRE-1/Sec-62 interaction.*** Three biological replicas of co-immunnoprecipitation followed western blot assays of Tm-treated A549 cells pulling on and probed for IRE-1 (Cell Signaling, Cat #3294S; Rabbit IgG, Cell Signaling, Cat #7074S), Sec-62 (Abcam, Cat #EPR9212; Light Chain-Specific Rabbit IgG (Cell Signaling, Cat #D4W3E), and BiP (Proteintech, Cat #66574; Mouse IgG, Cell Signaling Cat #7076S). “Recovery” indicates cells treated with Tm for 24 hours, then had the media replaced with Tm-free DMEM media supplemented with 10% FBS and 100 μg/mL penicillin and streptomycin mixture, and finally collected after given timepoint. Red box indicates what is shown in main text.

**Table S1.** *C. elegans* strains, laboratories of origin (“**Source**”), and plasmid injection concentrations for generation of transgenic animals used in this study.

**Table S2.** Target genes of siRNA and qPCR primers used in this study.

**Table S3.** Antibodies, blocking buffers, and dilutions used in this study.

**Table S4.** Lifespan data showing the number of animals and median lifespan per experiment. Bolded number indicates the biological replica in the order as displayed in the respective supplemental figure. *Shown in main text

**Table S5.** Paralysis data showing the number of animals and median day of paralysis per experiment. Bolded number indicates the biological replica in the order as displayed in the respective supplemental figure. *Shown in main text

**Table S6.** Genes, log fold changes (LogFC), log of counts per million reads (LogCPM), F-statistics (F), and p-values upregulated and downregulated by *hsp-3* siRNA relative to *pos-1* siRNA.

**Table S7.** Genes, log fold changes (LogFC), log of counts per million reads (LogCPM), F-statistics (F), and p-values upregulated and downregulated by *hsp-4* siRNA relative to *pos-1* siRNA.

**Table S8.** Significantly upregulated or downregulated genes on *hsp-3* and/or *hsp-4* siRNA.

