## Supplemental Figures for "Functionally diversified BiP orthologs control body growth, reproduction, stress resistance, aging, and ER-Phagy in *Caenorhabditis elegans*"

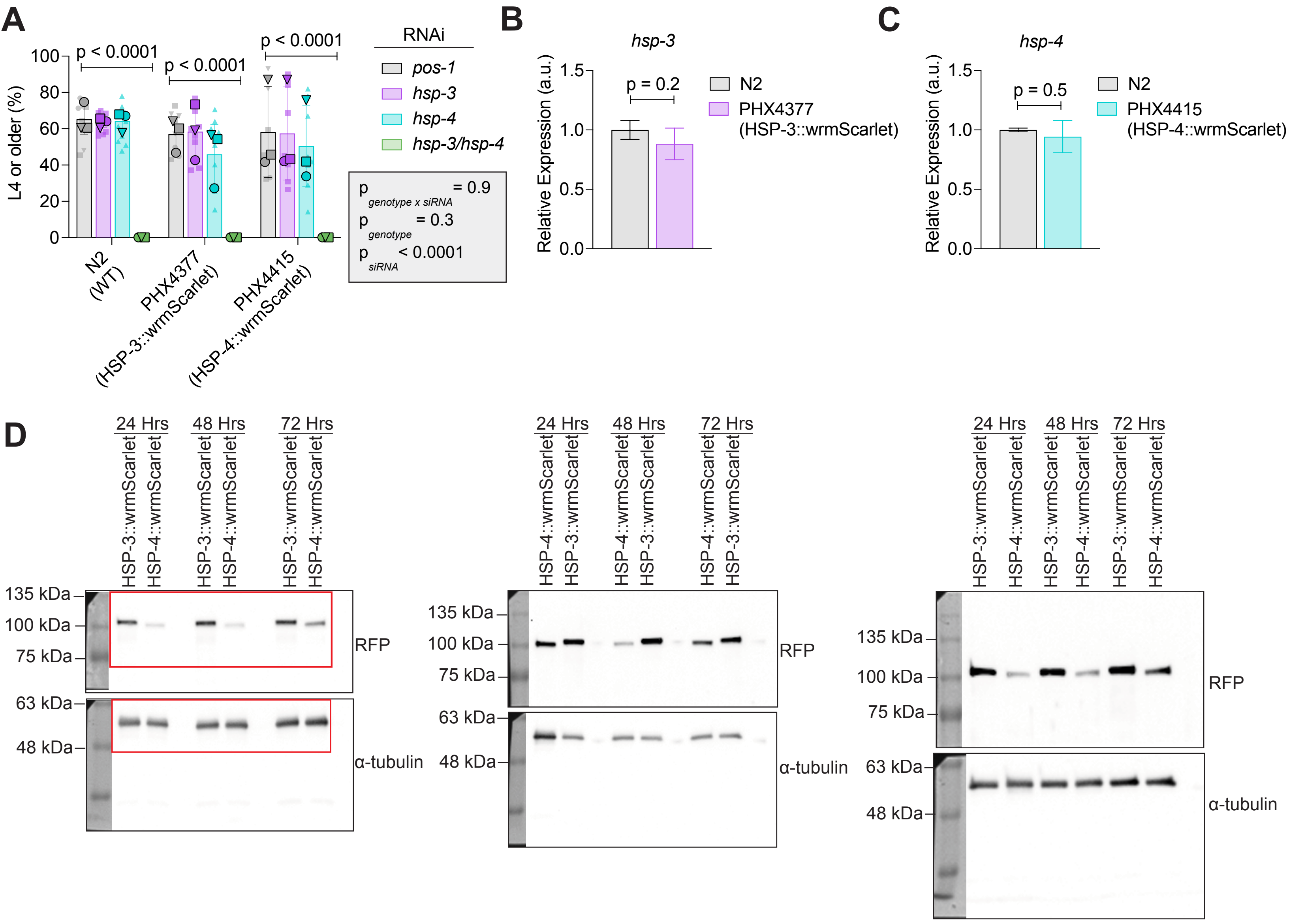
A
L4 or older (%)
p < 0.0001
p < 0.0001
p < 0.0001
N2 (WT)
PHX4377 (HSP-3::wrmScarlet)
PHX4415 (HSP-4::wrmScarlet)
RNAi
pos-1
hsp-3
hsp-4
hsp-3/hsp-4
p genotype x siRNA = 0.9
p genotype = 0.3
p siRNA < 0.0001
B
hsp-3
Relative Expression (a.u.)
p = 0.2
N2
PHX4377 (HSP-3::wrmScarlet)
C
hsp-4
p = 0.5
N2
PHX4415 (HSP-4::wrmScarlet)
D
24 Hrs
48 Hrs
72 Hrs
HSP-3::wrmScarlet
HSP-4::wrmScarlet
135 kDa
100 kDa
75 kDa
63 kDa
48 kDa
RFP
α-tubulin

### Figure S1

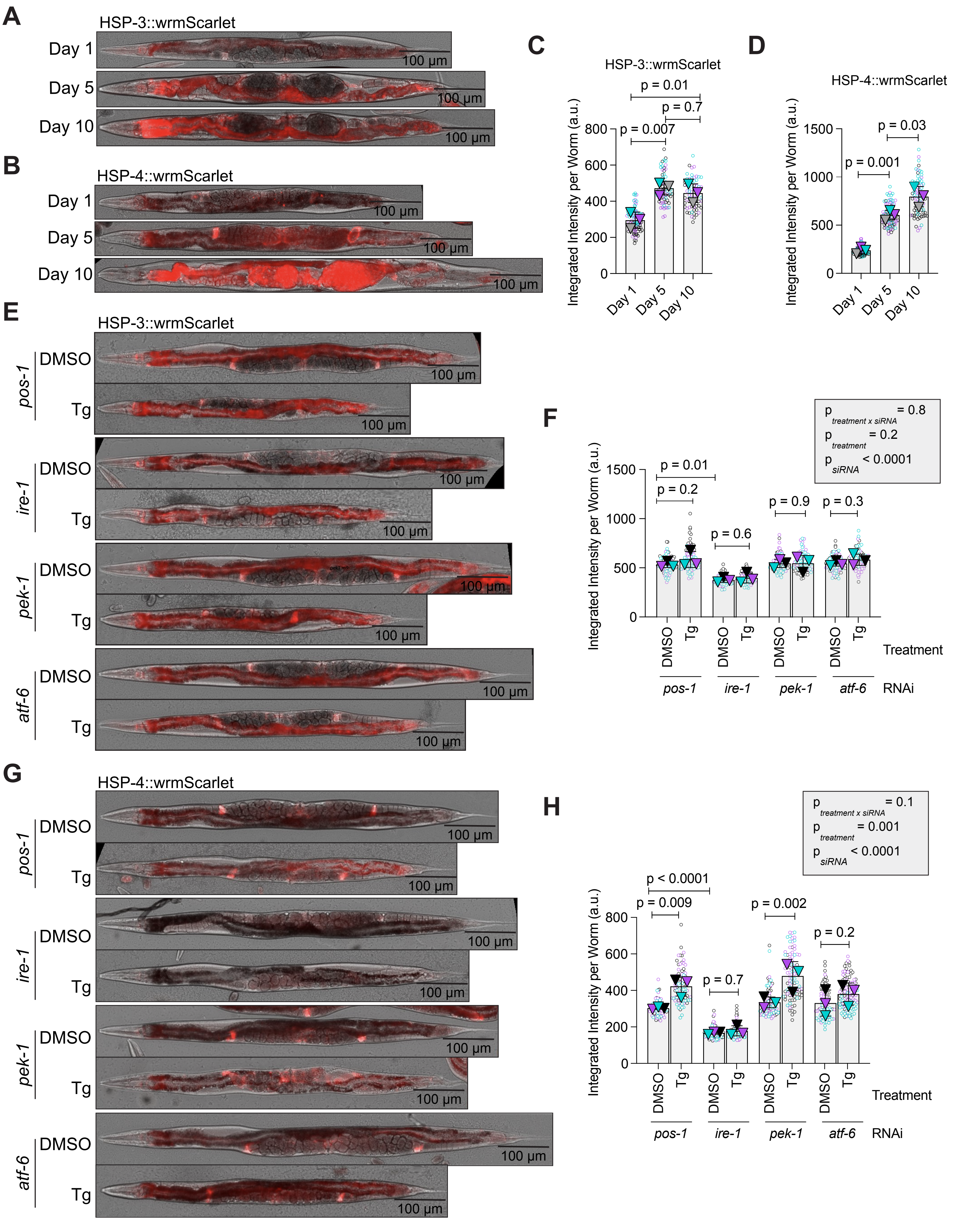
A
HSP-3::wrmScarlet
Day 1
Day 5
Day 10
100 µm
B
HSP-4::wrmScarlet
Day 1
Day 5
Day 10
100 µm
C
HSP-3::wrmScarlet
p = 0.01
p = 0.7
p = 0.007
Integrated Intensity per Worm (a.u.)
Day 1
Day 5
Day 10
D
HSP-4::wrmScarlet
p = 0.03
p = 0.001
Integrated Intensity per Worm (a.u.)
E
HSP-3::wrmScarlet
pos-1
ire-1
pek-1
atf-6
DMSO
Tg
F
p treatment x siRNA = 0.8
p treatment = 0.2
p siRNA < 0.0001
p = 0.01
p = 0.2
p = 0.6
p = 0.9
p = 0.3
Treatment
RNAi
G
HSP-4::wrmScarlet
H
p treatment x siRNA = 0.1
p treatment = 0.001
p siRNA < 0.0001
p < 0.0001
p = 0.009
p = 0.7
p = 0.002
p = 0.2

### Figure S2

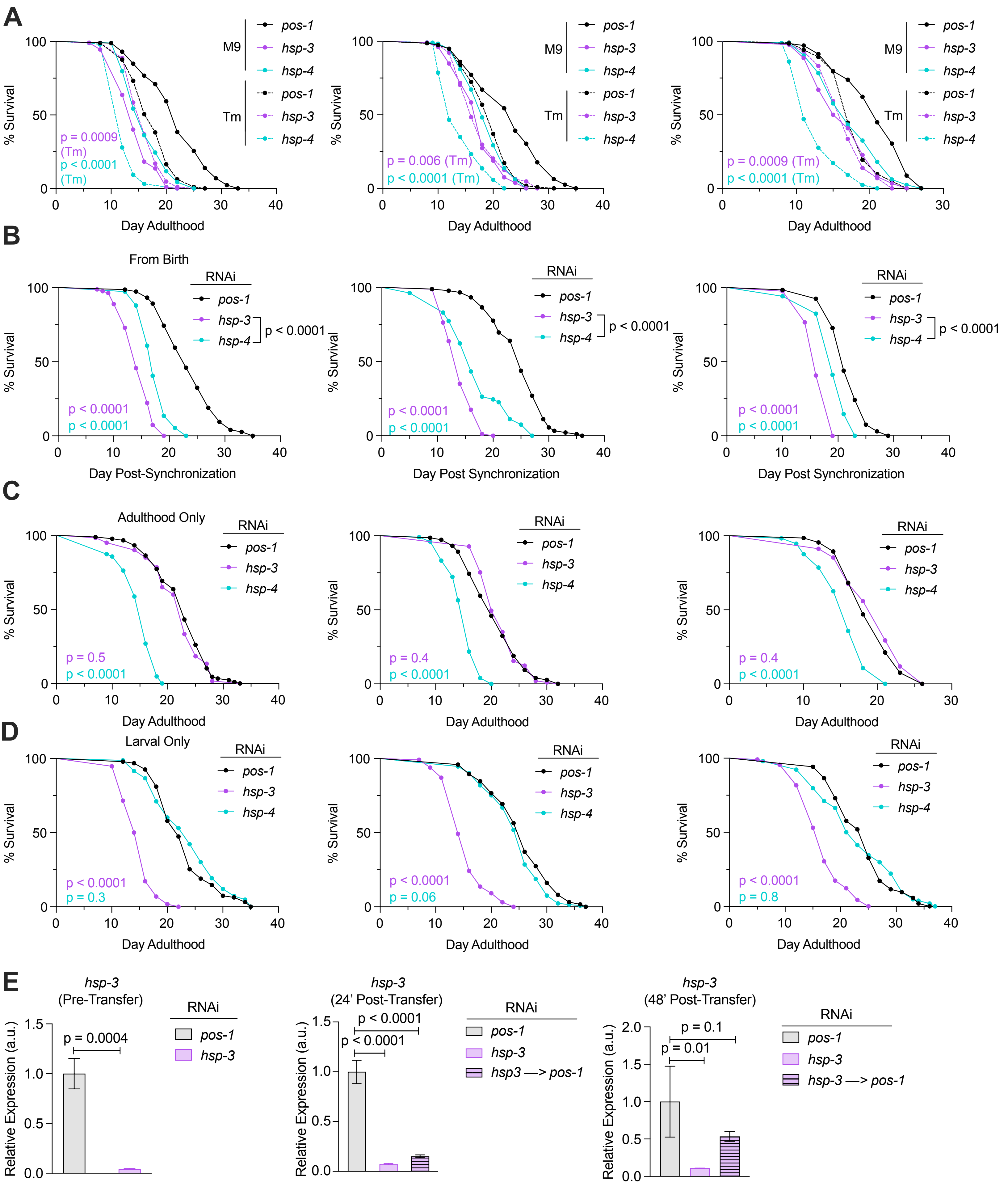

A
B
From Birth
C
Adulthood Only
D
Larval Only
E
% Survival
Day Adulthood
Day Post-Synchronization
Day Post Synchronization
M9
Tm
RNAi
pos-1
hsp-3
hsp-4
p = 0.0009 (Tm)
p < 0.0001 (Tm)
p = 0.006 (Tm)
p < 0.0001
p = 0.5
p = 0.4
p = 0.3
p = 0.06
p = 0.8
hsp-3 (Pre-Transfer)
hsp-3 (24' Post-Transfer)
hsp-3 (48' Post-Transfer)
Relative Expression (a.u.)
p = 0.0004
p = 0.1
p = 0.01
hsp3 —> pos-1
hsp-3 —> pos-1


### Figure S3

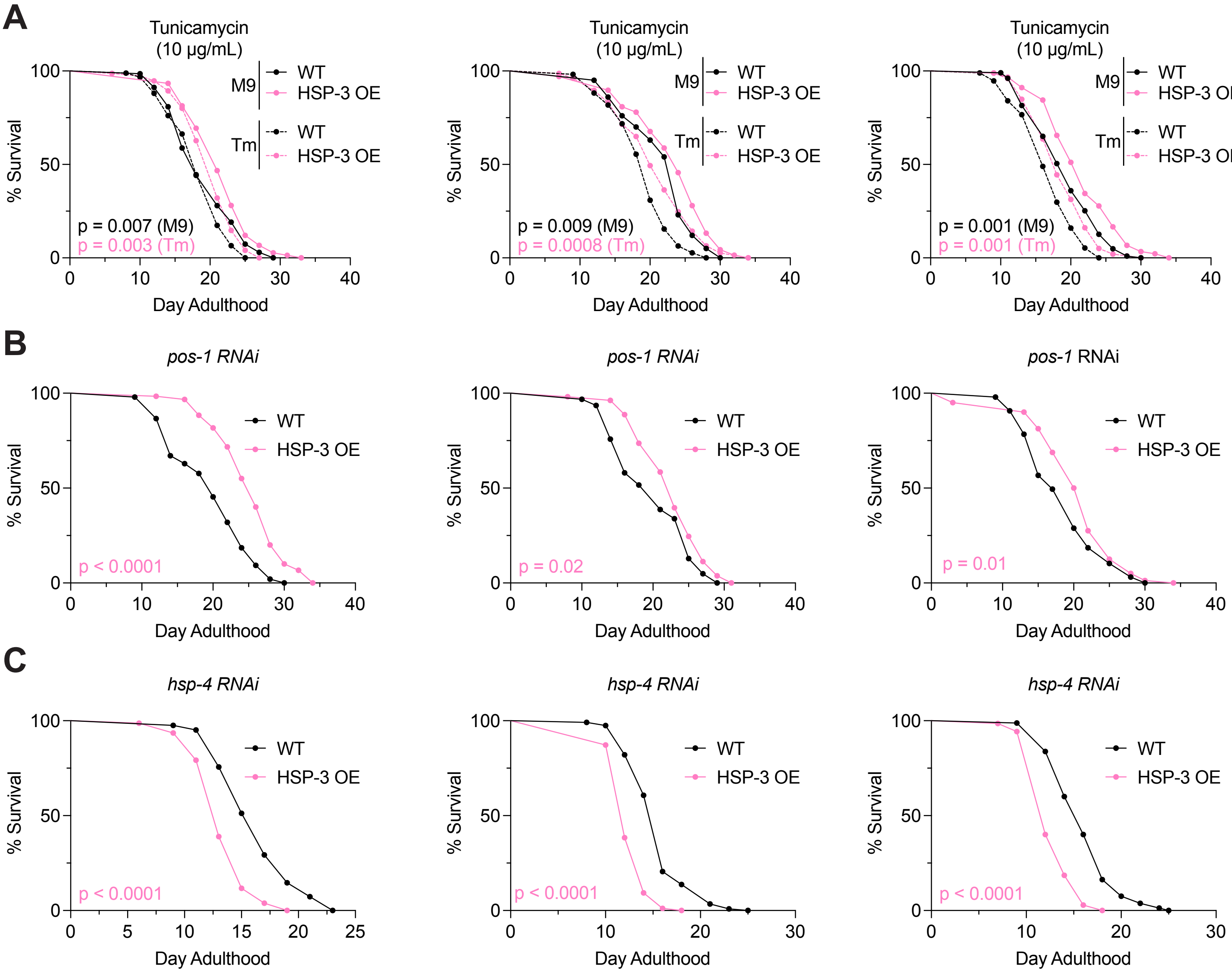
A
Tunicamycin
(10 μg/mL)
M9
WT
HSP-3 OE
Tm
WT
HSP-3 OE
p = 0.007 (M9)
p = 0.003 (Tm)
% Survival
Day Adulthood
Tunicamycin
(10 μg/mL)
p = 0.009 (M9)
p = 0.0008 (Tm)
Tunicamycin
(10 μg/mL)
p = 0.001 (M9)
p = 0.001 (Tm)
B
pos-1 RNAi
WT
HSP-3 OE
p < 0.0001
p = 0.02
pos-1 RNAi
p = 0.01
C
hsp-4 RNAi
WT
HSP-3 OE
p < 0.0001
p < 0.0001
p < 0.0001

### Figure S4

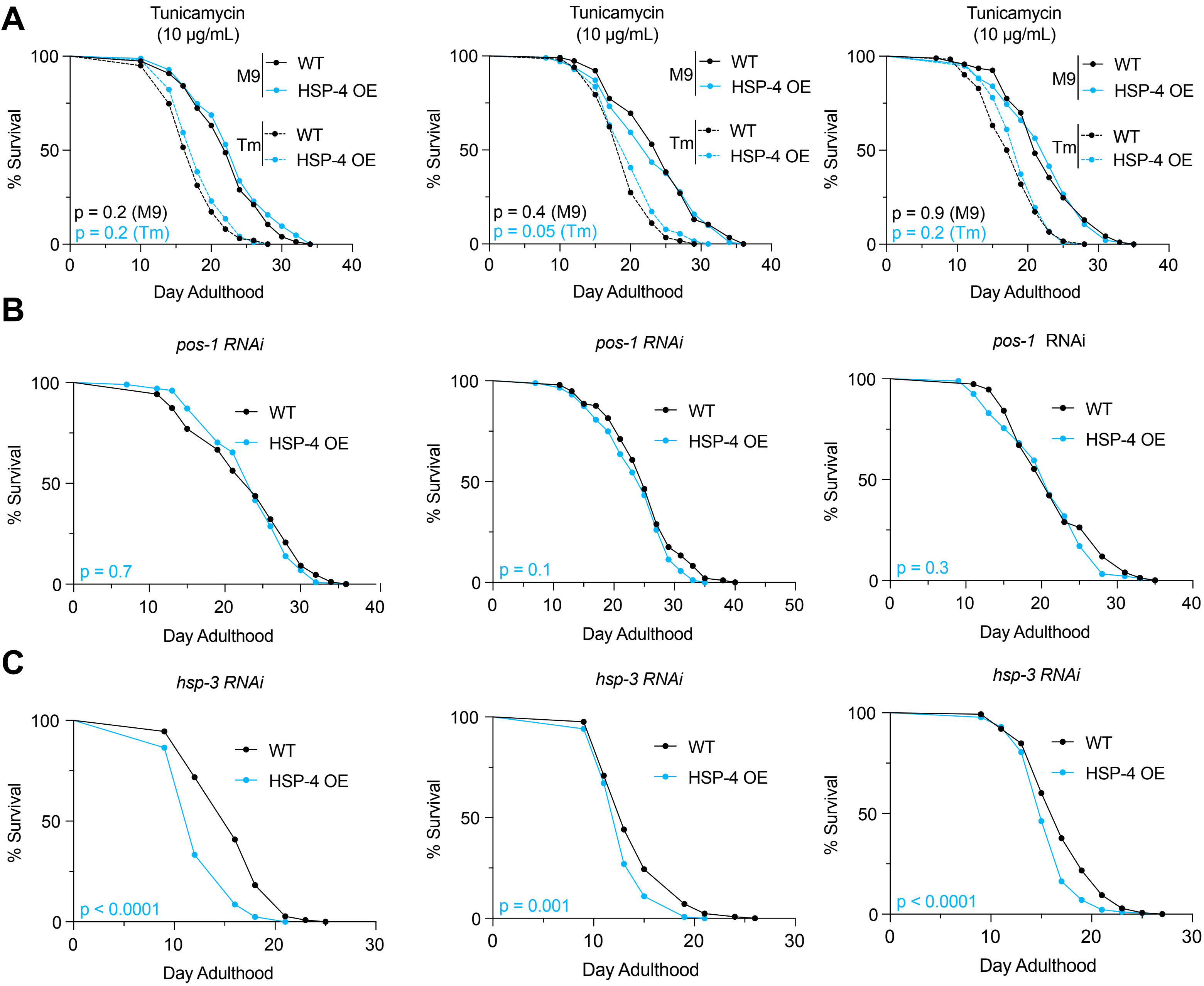
A
Tunicamycin (10 μg/mL)
M9 WT
M9 HSP-4 OE
Tm WT
Tm HSP-4 OE
p = 0.2 (M9)
p = 0.2 (Tm)
p = 0.4 (M9)
p = 0.05 (Tm)
p = 0.9 (M9)
p = 0.2 (Tm)
% Survival
Day Adulthood
B
pos-1 RNAi
WT
HSP-4 OE
p = 0.7
p = 0.1
p = 0.3
C
hsp-3 RNAi
p < 0.0001
p = 0.001
p < 0.0001

### Figure S5

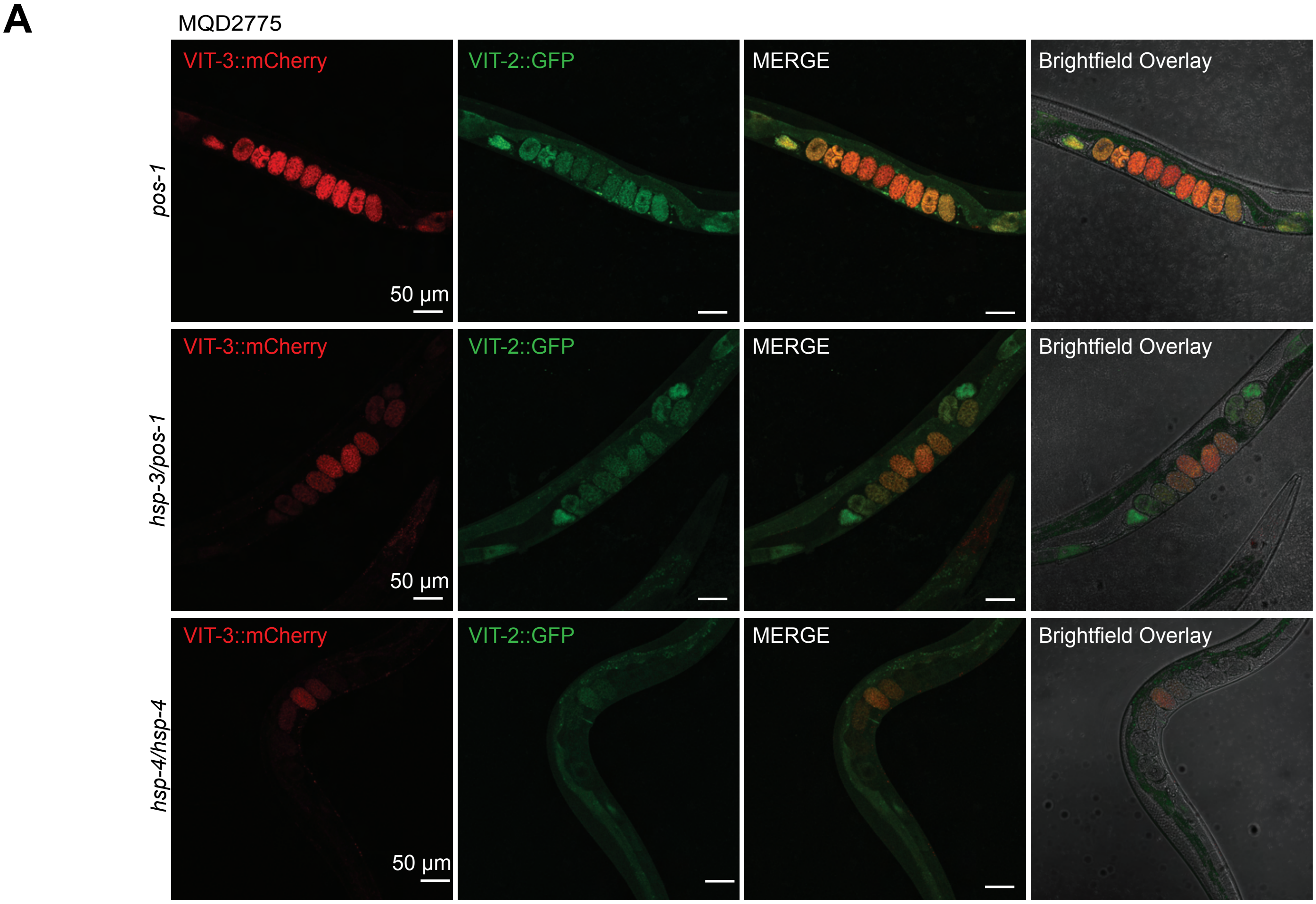
A
MQD2775
VIT-3::mCherry
VIT-2::GFP
MERGE
Brightfield Overlay
pos-1
hsp-3/pos-1
hsp-4/hsp-4
50 µm
50 µm
50 µm

### Figure S6

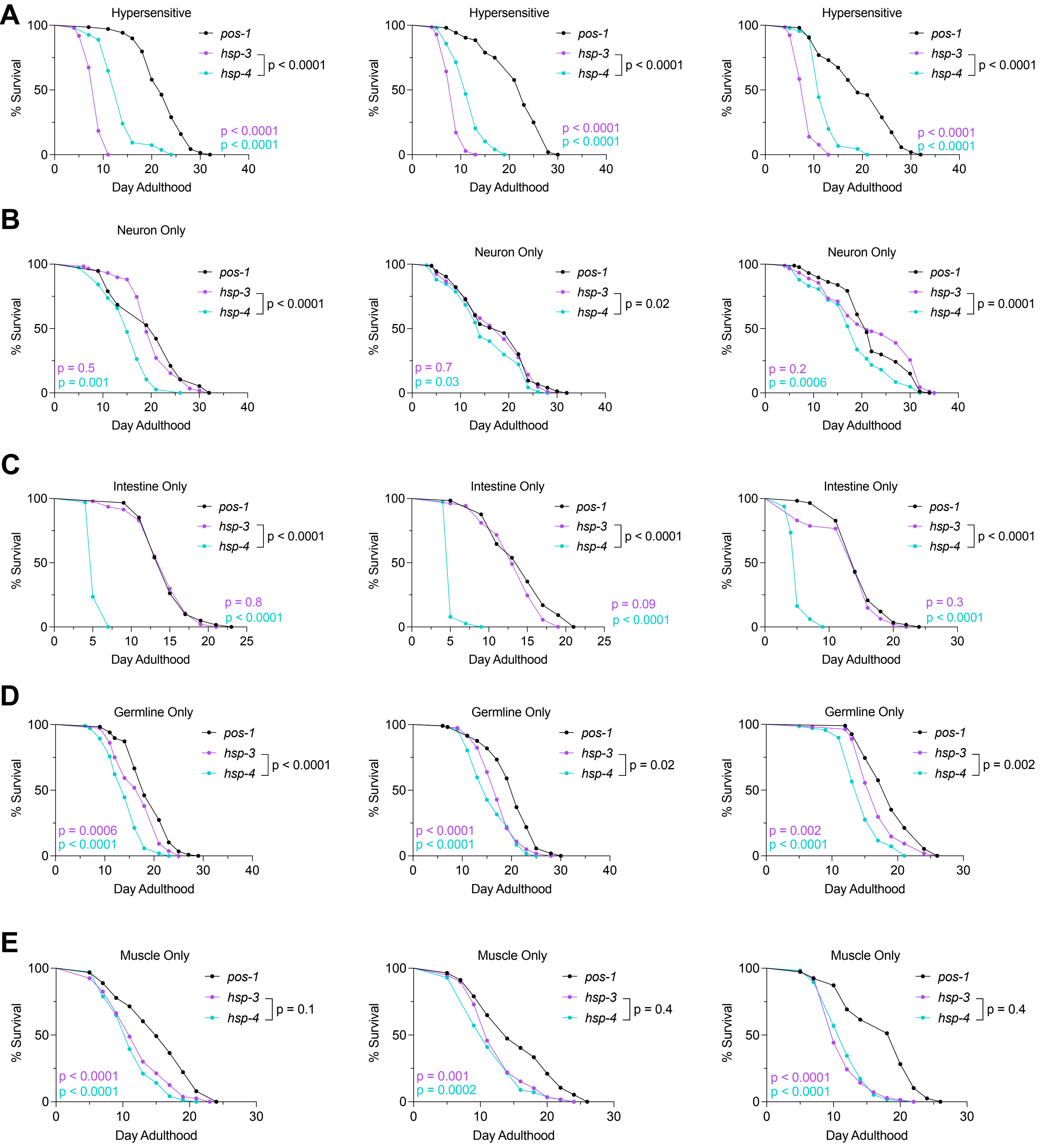
A
Hypersensitive
% Survival
Day Adulthood
pos-1
hsp-3
hsp-4
p < 0.0001
p < 0.0001
p < 0.0001
B
Neuron Only
p < 0.0001
p = 0.5
p = 0.001
p = 0.02
p = 0.7
p = 0.03
p = 0.0001
p = 0.2
p = 0.0006
C
Intestine Only
p < 0.0001
p = 0.8
p < 0.0001
p = 0.09
p < 0.0001
p = 0.3
p < 0.0001
D
Germline Only
p < 0.0001
p = 0.0006
p < 0.0001
p = 0.02
p < 0.0001
p < 0.0001
p = 0.002
p = 0.002
p < 0.0001
E
Muscle Only
p = 0.1
p < 0.0001
p < 0.0001
p = 0.4
p = 0.001
p = 0.0002
p = 0.4
p < 0.0001
p < 0.0001

### Figure S7

**A**

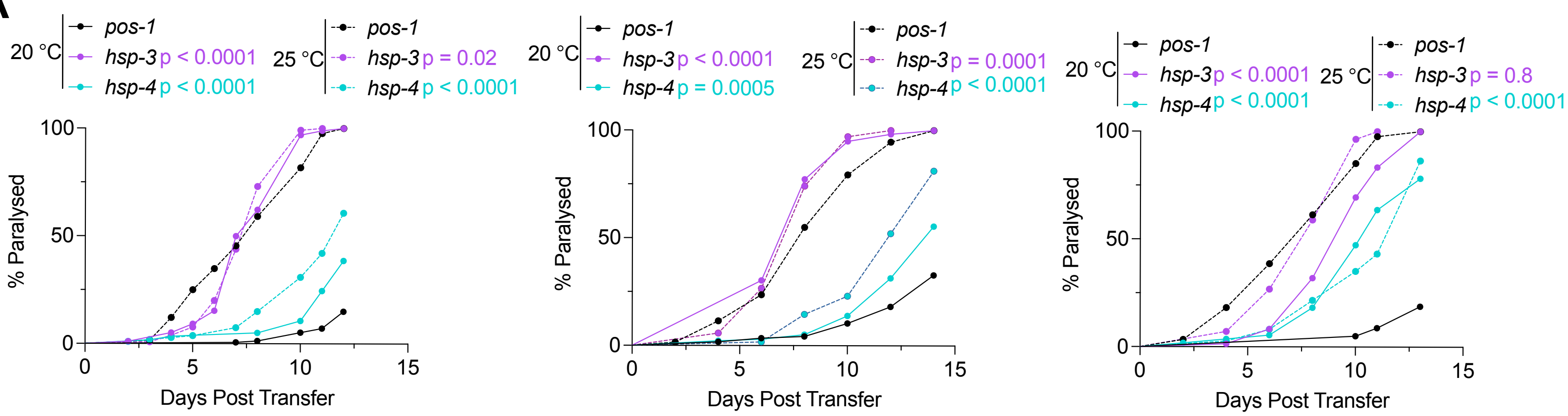
20 °C
pos-1
hsp-3 p < 0.0001
hsp-4 p < 0.0001
25 °C
pos-1
hsp-3 p = 0.02
hsp-4 p < 0.0001
% Paralysed
Days Post Transfer
20 °C
pos-1
hsp-3 p < 0.0001
hsp-4 p = 0.0005
25 °C
pos-1
hsp-3 p = 0.0001
hsp-4 p < 0.0001
% Paralysed
Days Post Transfer
20 °C
pos-1
hsp-3 p < 0.0001
hsp-4 p < 0.0001
25 °C
pos-1
hsp-3 p = 0.8
hsp-4 p < 0.0001
% Paralysed
Days Post Transfer

### Figure S8

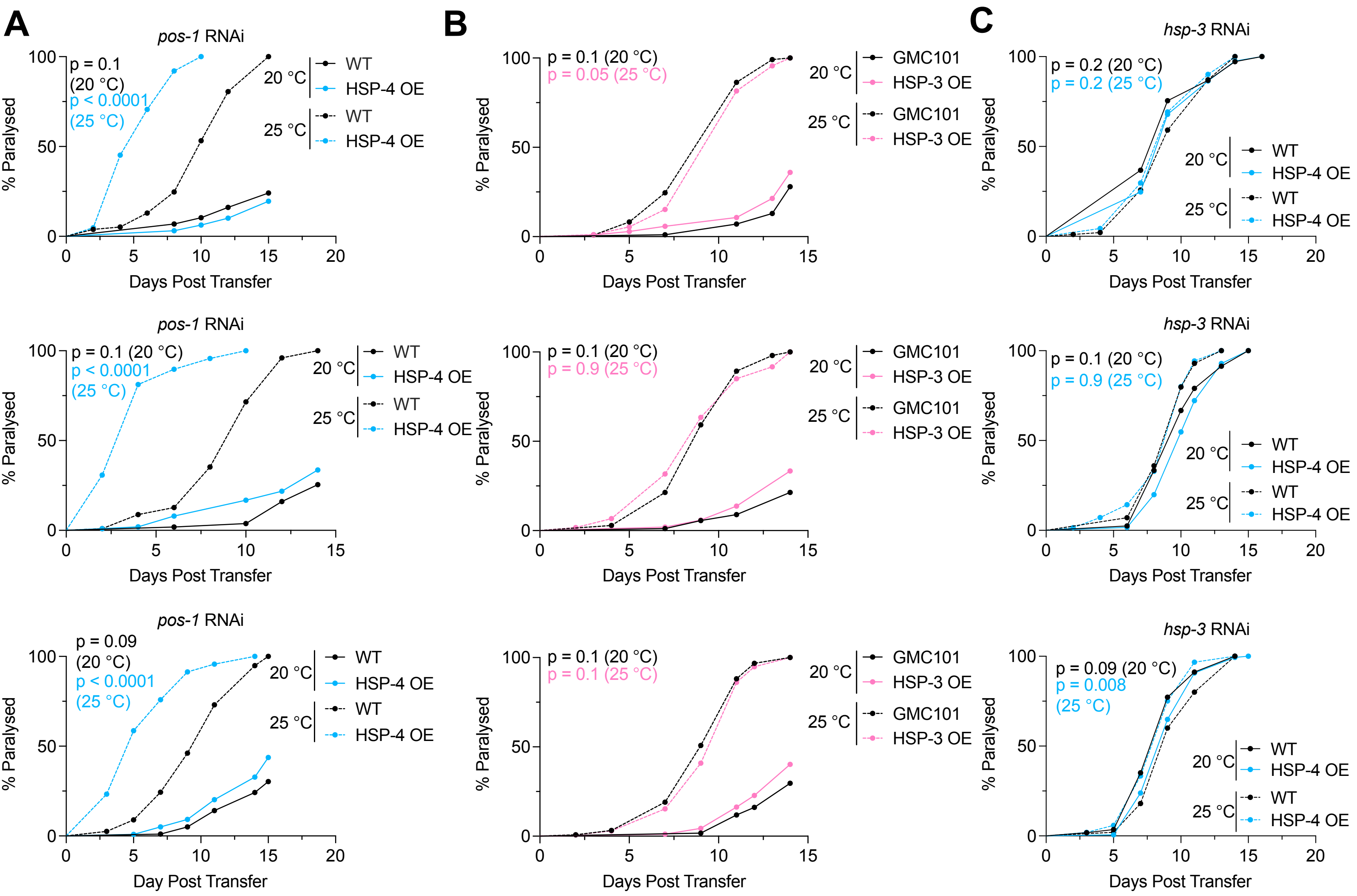
A
B
C
pos-1 RNAi
p = 0.1 (20 °C)
p < 0.0001 (25 °C)
20 °C
25 °C
WT
HSP-4 OE
% Paralysed
Days Post Transfer
p = 0.09 (20 °C)
Day Post Transfer
p = 0.1 (20 °C)
p = 0.05 (25 °C)
GMC101
HSP-3 OE
p = 0.9 (25 °C)
p = 0.1 (25 °C)
hsp-3 RNAi
p = 0.2 (20 °C)
p = 0.2 (25 °C)
p = 0.008 (25 °C)

### Figure S9

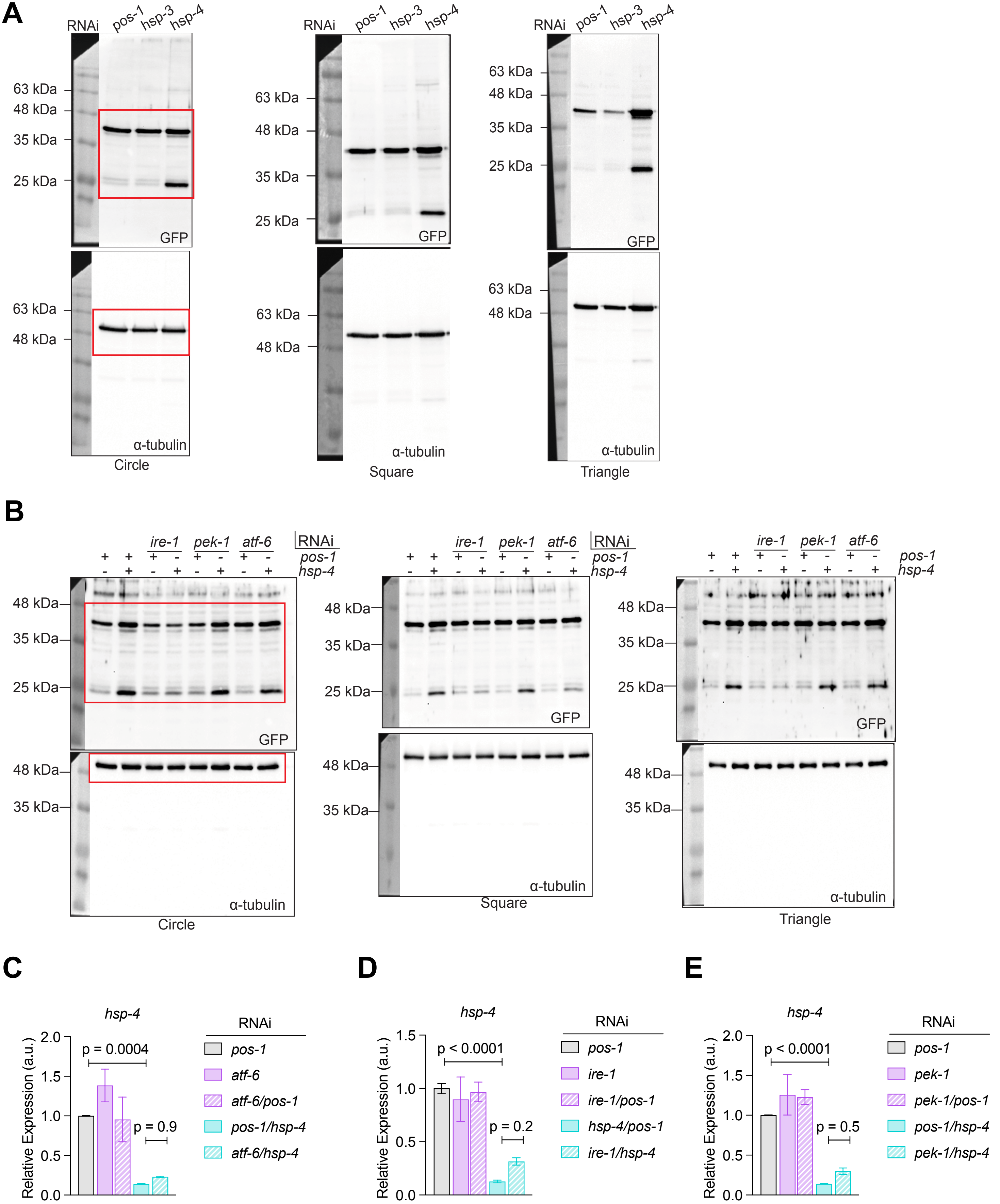
A
RNAi
pos-1
hsp-3
hsp-4
63 kDa
48 kDa
35 kDa
25 kDa
GFP
α-tubulin
Circle
Square
Triangle
B
ire-1
pek-1
atf-6
RNAi
pos-1
hsp-4
GFP
α-tubulin
C
D
E
hsp-4
Relative Expression (a.u.)
p = 0.0004
p = 0.9
p < 0.0001
p = 0.2
p = 0.5
pos-1
atf-6
atf-6/pos-1
pos-1/hsp-4
atf-6/hsp-4
ire-1
ire-1/pos-1
hsp-4/pos-1
ire-1/hsp-4
pek-1
pek-1/pos-1
pek-1/hsp-4

### Figure S10

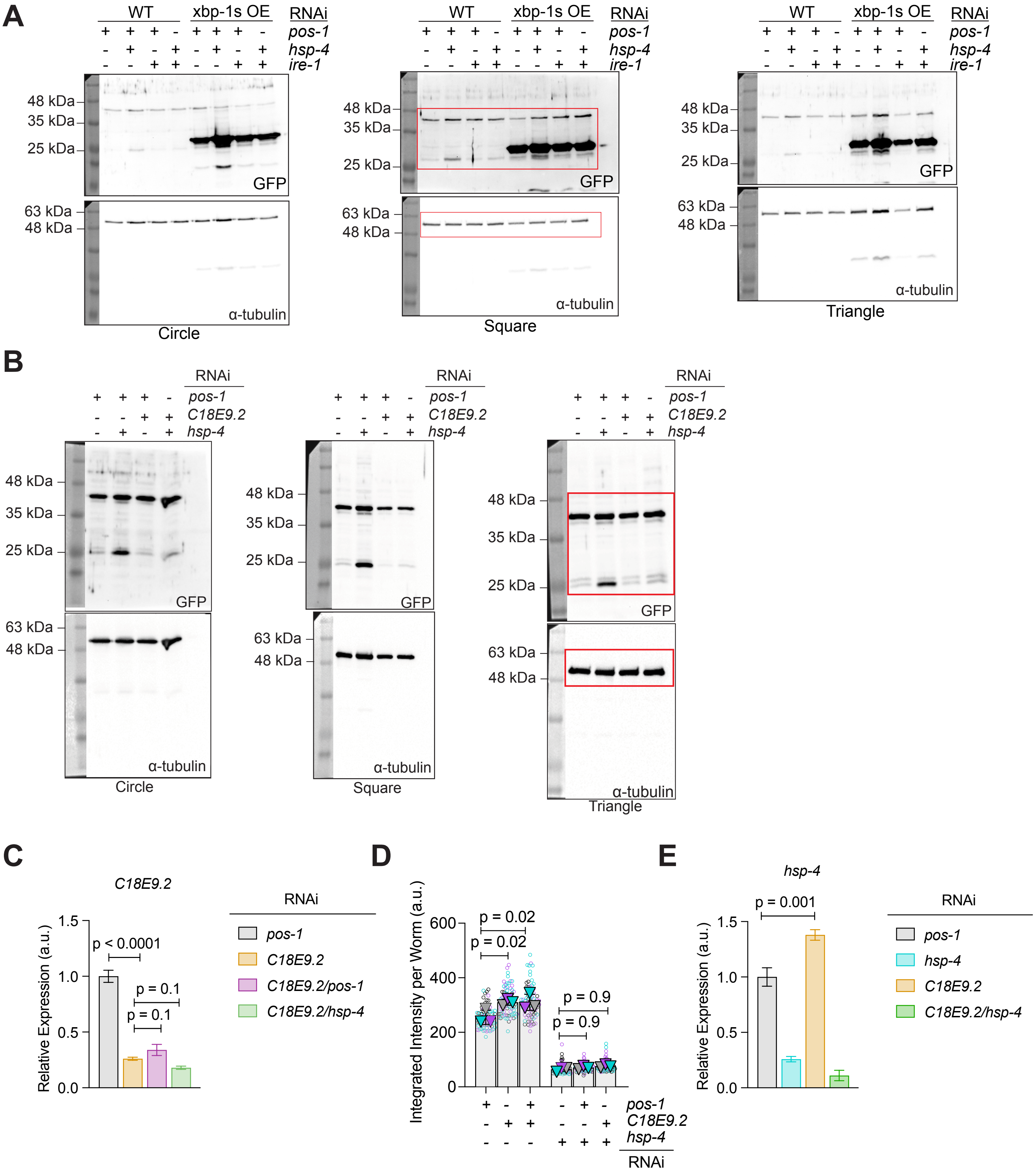
A
WT
xbp-1s OE
RNAi
pos-1
hsp-4
ire-1
48 kDa
35 kDa
25 kDa
GFP
63 kDa
48 kDa
α-tubulin
Circle
Square
Triangle
B
RNAi
pos-1
C18E9.2
hsp-4
GFP
α-tubulin
Circle
Square
Triangle
C
C18E9.2
Relative Expression (a.u.)
p < 0.0001
p = 0.1
p = 0.1
RNAi
pos-1
C18E9.2
C18E9.2/pos-1
C18E9.2/hsp-4
D
Integrated Intensity per Worm (a.u.)
p = 0.02
p = 0.02
p = 0.9
p = 0.9
pos-1
C18E9.2
hsp-4
RNAi
E
hsp-4
Relative Expression (a.u.)
p = 0.001
RNAi
pos-1
hsp-4
C18E9.2
C18E9.2/hsp-4

### Figure S11

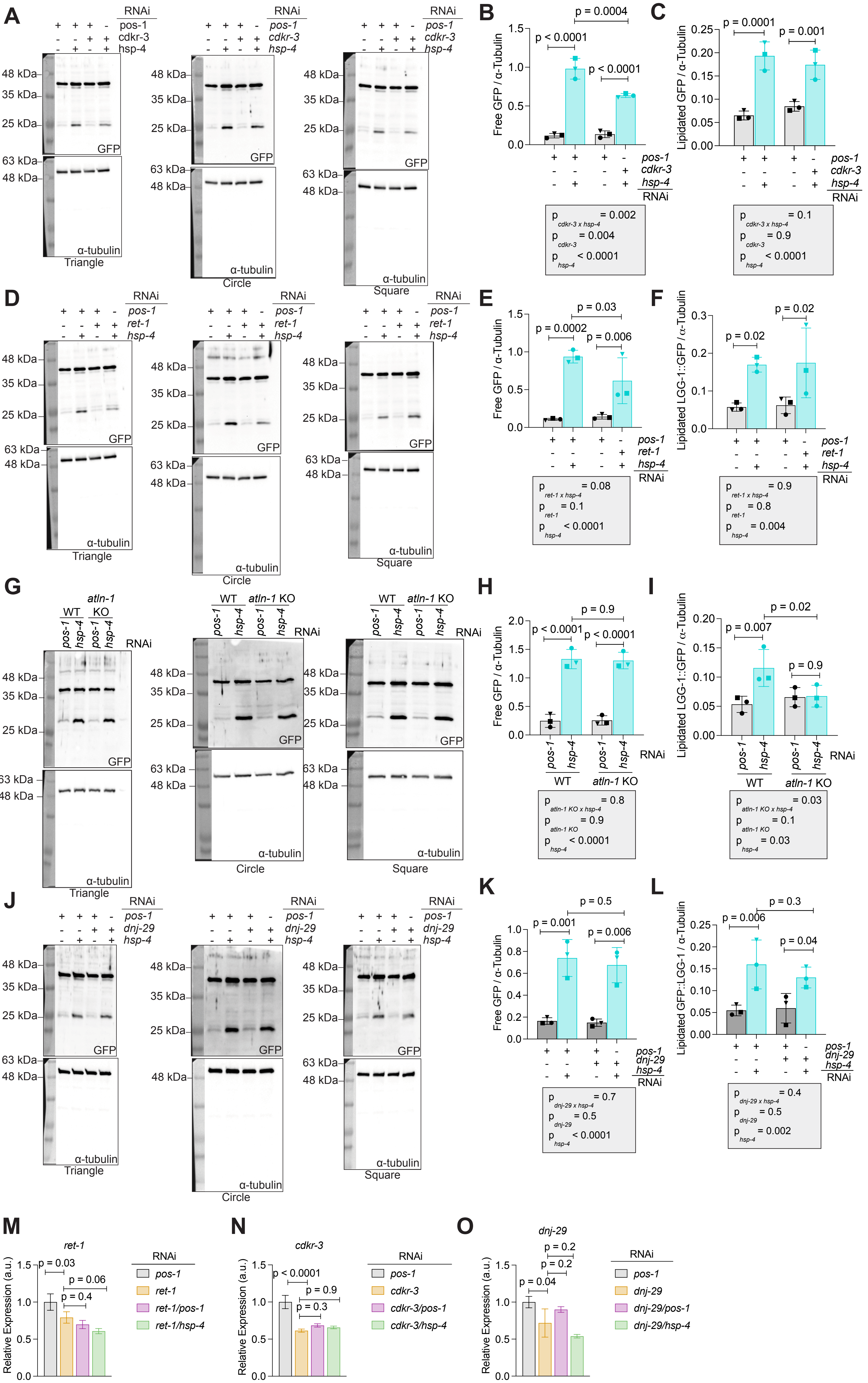
A
RNAi
pos-1
cdkr-3
hsp-4
48 kDa
35 kDa
25 kDa
63 kDa
GFP
α-tubulin
Triangle
Circle
Square
B
Free GFP / α-Tubulin
p = 0.0004
p < 0.0001
p < 0.0001
p cdkr-3 x hsp-4 = 0.002
p cdkr-3 = 0.004
p hsp-4 < 0.0001
C
Lipidated GFP / α-Tubulin
p = 0.0001
p = 0.001
p cdkr-3 x hsp-4 = 0.1
p cdkr-3 = 0.9
p hsp-4 < 0.0001
D
ret-1
E
p = 0.03
p = 0.0002
p = 0.006
p ret-1 x hsp-4 = 0.08
p ret-1 = 0.1
p hsp-4 < 0.0001
F
Lipidated LGG-1::GFP / α-Tubulin
p = 0.02
p = 0.02
p ret-1 x hsp-4 = 0.9
p ret-1 = 0.8
p hsp-4 = 0.004
G
WT
atln-1 KO
H
p = 0.9
p < 0.0001
p < 0.0001
p atln-1 KO x hsp-4 = 0.8
p atln-1 KO = 0.9
p hsp-4 < 0.0001
I
p = 0.02
p = 0.007
p = 0.9
p atln-1 KO x hsp-4 = 0.03
p atln-1 KO = 0.1
p hsp-4 = 0.03
J
dnj-29
K
p = 0.5
p = 0.001
p = 0.006
p dnj-29 x hsp-4 = 0.7
p dnj-29 = 0.5
p hsp-4 < 0.0001
L
Lipidated GFP::LGG-1 / α-Tubulin
p = 0.3
p = 0.006
p = 0.04
p dnj-29 x hsp-4 = 0.4
p dnj-29 = 0.5
p hsp-4 = 0.002
M
ret-1
Relative Expression (a.u.)
p = 0.03
p = 0.06
p = 0.4
ret-1/pos-1
ret-1/hsp-4
N
cdkr-3
p < 0.0001
p = 0.9
p = 0.3
cdkr-3/pos-1
cdkr-3/hsp-4
O
dnj-29
p = 0.2
p = 0.2
p = 0.04
dnj-29/pos-1
dnj-29/hsp-4

**Figure S12**

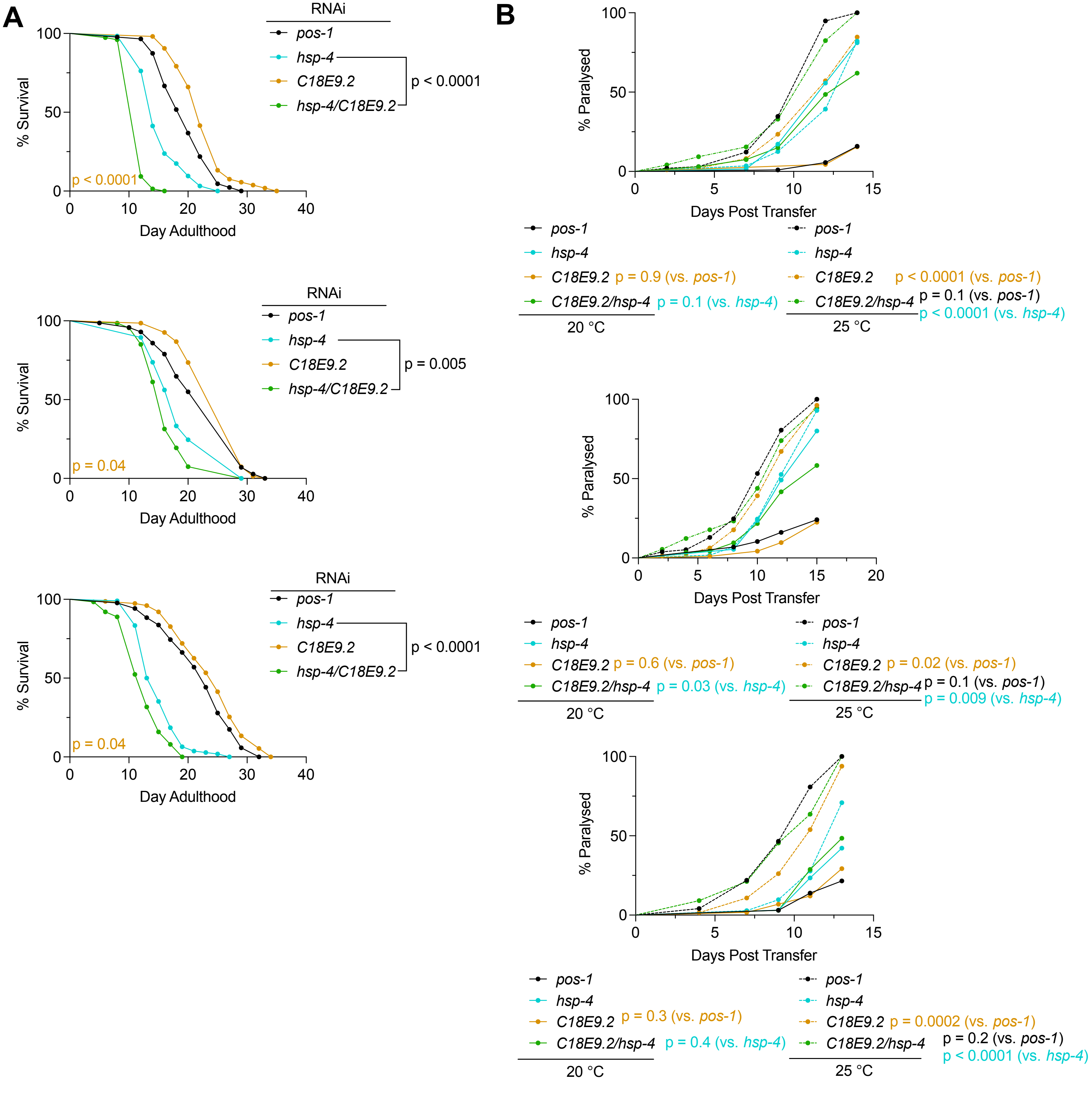

A
RNAi
pos-1
hsp-4
C18E9.2
hsp-4/C18E9.2
p < 0.0001
% Survival
Day Adulthood
p < 0.0001
p = 0.005
p = 0.04
p < 0.0001
p = 0.04
B
% Paralysed
Days Post Transfer
pos-1
hsp-4
C18E9.2 p = 0.9 (vs. pos-1)
C18E9.2/hsp-4 p = 0.1 (vs. hsp-4)
20 °C
C18E9.2 p < 0.0001 (vs. pos-1)
C18E9.2/hsp-4 p = 0.1 (vs. pos-1)
p < 0.0001 (vs. hsp-4)
25 °C
C18E9.2 p = 0.6 (vs. pos-1)
C18E9.2/hsp-4 p = 0.03 (vs. hsp-4)
C18E9.2 p = 0.02 (vs. pos-1)
C18E9.2/hsp-4 p = 0.1 (vs. pos-1)
p = 0.009 (vs. hsp-4)
C18E9.2 p = 0.3 (vs. pos-1)
C18E9.2/hsp-4 p = 0.4 (vs. hsp-4)
C18E9.2 p = 0.0002 (vs. pos-1)
C18E9.2/hsp-4 p = 0.2 (vs. pos-1)
p < 0.0001 (vs. hsp-4)


### Figure S13

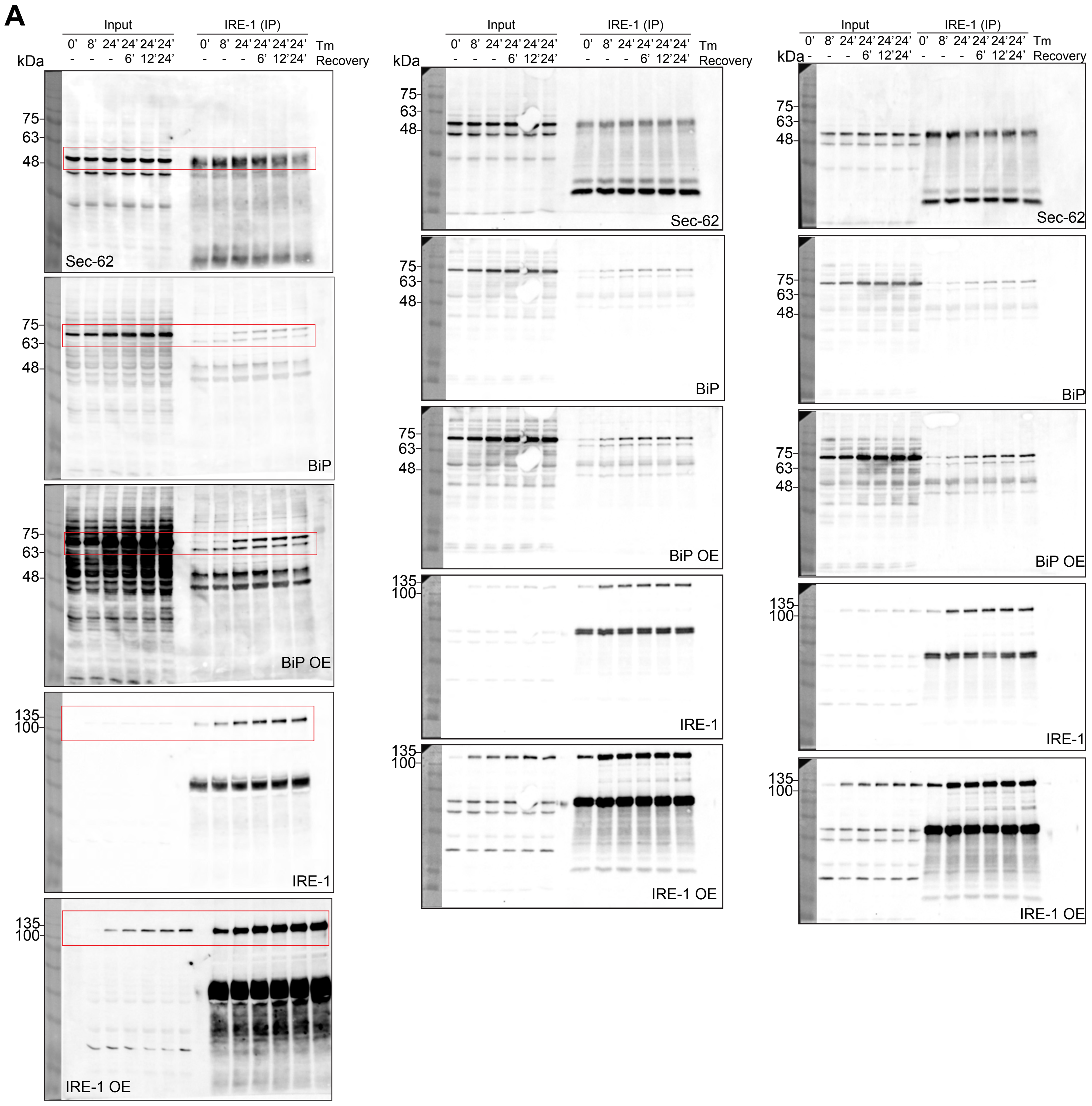
A
Input
IRE-1 (IP)
0' 8' 24' 24' 24' 24'
Tm
- - - 6' 12' 24'
Recovery
kDa
75
63
48
135
100
Sec-62
BiP
BiP OE
IRE-1
IRE-1 OE

### Figure S14
