## Supplemental Tables for "Functionally diversified BiP orthologs control body growth, reproduction, stress resistance, aging, and ER-Phagy in *Caenorhabditis elegans*"

| **Name** | **Description** | **Source** | **Plasmid Concentration** |
| --- | --- | --- | --- |
| N2 (Bristol) | *wild type* | Caenorhabditis Genetics Center (CGC) |  |
| DA2123 | *adIs2122 [lgg-1p::GFP::lgg-1 + rol-6(su1006)]* | Caenorhabditis Genetics Center (CGC) |  |
| GMC101 | *dvIs100 [unc-54p::A-beta-1-42::unc-54 3'-UTR + mtl-2p::GFP]* | Caenorhabditis Genetics Center (CGC) |  |
| BW1940 | *ctIs40 [dbl-1(+) + sur-5::GFP]* | Caenorhabditis Genetics Center (CGC) |  |
| PHX4377 | *hsp-3::wrmScarlet* | SUNY Biotech (CRISPR-Cas9) |  |
| PHX4415 | *hsp-4::wrmScarlet* | SUNY Biotech (CRISPR-Cas9) |  |
| MTX203 | *mtmEx74 [eef-1A.1p::HA::hsp-3; myo-2p::gfp]* | SUNY Biotech | 30 ng/uL (eef-1A.1p::hsp-3); 20 ng/uL (myo-2::gfp) |
| MTX205 | *mtmEx76 [eef-1A.1p::HA::hsp-4; myo-2p::gfp]* | SUNY Biotech | 30 ng/uL (eef-1A.1p::hsp-4); 20 ng/uL (myo-2::gfp) |
| MTX291 | *mtmIs19 [eef-1A.1p::HA::hsp-4; myo-2p::gfp]* | Truttmann Lab (UV-integration of MTX205) | 30 ng/uL (eef-1A.1p::hsp-4); 20 ng/uL (myo-2::gfp) |
| MTX300 | *dvIs100 [unc-54p::A-beta-1-42::unc-54 3'-UTR + mtl-2p::GFP]; mtmIs19 [eef-1A.1p::hsp-4 myo-2p::gfp]* | Truttmann Lab |  |
| MTX328 | *dvIs100 [unc-54p::A-beta-1-42::unc-54 3'-UTR + mtl-2p::GFP]; mtmEx74 [eef-1A.1p::hsp-3; myo-2p::gfp* | Truttmann Lab |  |
| MTX339 | *agd925 [sur-5p::xbp-1s; myo-2::rfp] x adI2122 [lgg-1::GFP::lgg-1] (AGD925 x DA2123)* | Truttmann Lab; AGD925 gift from Dillin lab |  |
| MTX341 | *Y54G2A.2(ok1144) x adI2122 [lgg-1::GFP::lgg-1] (RB1127 x DA2123)* | Truttmann Lab/Caenorhabditis Genetics Center (CGC) |  |

Table S1

| **Target** | **Library of Origin (siRNA)** | **Forward rt-qPCR Sequencing Primer  (5’-3’)** | **Reverse rt-qPCR Sequence Primer**  **(5’-3’)** |
| --- | --- | --- | --- |
| *pos-1* | Vidal |  |  |
| *hsp-3* | Vidal | ACCGTCACCATCCAGGTC | TCCGGTGAGGTCGAACTTT |
| *hsp-4* | Vidal | CAGATGAAAACTCAAATCGCC | GGTTGCTTCCGAGCCACTCAA |
| *ire-1* | Vidal | AAGTGCCGTTTTTGCCGTTT | TGAGGACAAGACCATTGGACAG |
| *pek-1* | Vidal | CTGAGAAGGCAACGCTCTCT | ATCACCGCTACTCTGGATGG |
| *atf-6* | Vidal | CCTACTTTACGGGACCGACG | AATCTCCTAAACTCCCGCCG |
| *C18E9.2* | Ahringer | GACAGCCACGACCATGTTTG | ATTCCCAAAGAGTATCGACAGC |
| *dnj-29* | Vidal | TCCAGGAGCCTCGAAACCAG | TCACAACCGTCGCATTGGCA |
| *ret-1* | Ahringer | AAGGATATTAGTGATGAAGATGTGAAGC | ATGTGGCGATGATGGATGAG |
| *cdkr-3* | Ahringer | GTCAGACGACTTGCCAATCGATAT | CGGCATGTCCAAAATGGCAT |
| *cdc42* | N/A | CTGCTGGACAGGAAGATTACG | CTCGGACATTCTCGAATGAAG |

Table S2

| **Target/Epitope** | **Manufacturer** | **Catalog #** | **Blocking Solution** | **Dilution Solution** | **Dilution/Concentration** |
| --- | --- | --- | --- | --- | --- |
| GFP | Roche | 11814460001 | 5% milk/0.1% TBS-T | 2.5% milk/0.1% TBS-T | 1:1,000 |
| RFP | Chromotek/Proteintech | 6g6 | 5% milk/0.1% TBS-T | 2.5% milk/0.1% TBS-T | 1:1,000 |
| ɑ-tubulin | Developmental Studies Hybridoma Bank | 12G10 | 5% milk/0.1% TBS-T | 2.5% milk/0.1% TBS-T | 0.4 ug/mL |
| Mouse IgG | Cell Signaling | 7076S | N/A | 0.1% TBS-T | 1:10,000 |
| Sec62 | Abcam | EPR9212 | 5% BSA/0.1% TBS-1 | 2.5% BSA/0.1% TBS-T | 1:1,000 |
| Rabbit IgG (Light-Chain Specific) | Cell Signaling | D4W3E | N/A | 2.5% BSA/0.1% TBS-T | 1:10,000 |
| Grp78/BiP | Proteintech | 66574 | 5% BSA/0.1% TBS-3 | 2.5% BSA/0.1% TBS-T | 1:1,000 |
| Mouse IgG | Cell Signaling | 7076S | N/A | 2.5% BSA/0.1% TBS-T | 1:10,000 |
| IRE1α | Cell Signaling | 3294S | 5% BSA/0.1% TBS-5 | 2.5% BSA/0.1% TBS-T | 1:1,000 |
| Rabbit IgG | Cell Signaling | 7074S | N/A | 2.5% BSA/0.1% TBS-T | 1:10,000 |

Table S3

| **Fig. 1H/Fig. S3A** |  |  |  |  |  |  |  |  |  |  |  |  |  |  |  |  |  |  |  |  |
| --- | --- | --- | --- | --- | --- | --- | --- | --- | --- | --- | --- | --- | --- | --- | --- | --- | --- | --- | --- | --- |
| **#1*** | M9 | M9 | M9 | Tm | Tm | Tm | **#2** | M9 | M9 | M9 | Tm | Tm | Tm | **#3** | M9 | M9 | M9 | Tm | Tm | Tm |
|  | *pos-1* | *hsp-3* | *hsp-4* | *pos-1* | *hsp-3* | *hsp-4* |  | *pos-1* | *hsp-3* | *hsp-4* | *pos-1* | *hsp-3* | *hsp-4* |  | *pos-1* | *hsp-3* | *hsp-4* | *pos-1* | *hsp-3* | *hsp-4* |
| # of Animals | 86 | 110 | 94 | 97 | 106 | 97 |  | 79 | 106 | 116 | 116 | 128 | 103 |  | 69 | 88 | 83 | 103 | 101 | 87 |
| Median Survival | 22 | 14 | 16 | 18 | 16 | 12 |  | 24 | 18 | 18 | 20 | 16 | 12 |  | 21 | 16 | 17 | 17 | 17 | 11 |
| **Fig. 1I/Fig. S4A** |  |  |  |  |  |  |  |  |  |  |  |  |  |  |  |  |  |  |  |  |
| **#1*** | M9 | M9 | Tm | Tm |  |  | **#2** | M9 | M9 | Tm | Tm |  |  | **#3** | M9 | M9 | Tm | Tm |  |  |
|  | WT | HSP-3 OE | WT | HSP-3 OE |  |  |  | WT | HSP-3 OE | WT | HSP-3 OE |  |  |  | WT | HSP-3 OE | WT | HSP-3 OE |  |  |
| # of Animals | 68 | 75 | 92 | 75 |  |  |  | 100 | 68 | 110 | 77 |  |  |  | 103 | 90 | 94 | 99 |  |  |
| Median Survival | 18 | 21 | 18 | 21 |  |  |  | 24 | 24 | 20 | 20 |  |  |  | 20 | 22 | 16 | 18 |  |  |
| **Fig. 1J/Fig. S5A** |  |  |  |  |  |  |  |  |  |  |  |  |  |  |  |  |  |  |  |  |
| **#1*** | M9 | M9 | Tm | Tm |  |  | **#2** | M9 | M9 | Tm | Tm |  |  | **#3** | M9 | M9 | Tm | Tm |  |  |
|  | WT | HSP-4 OE | WT | HSP-4 OE |  |  |  | WT | HSP-4 OE | WT | HSP-4 OE |  |  |  | WT | HSP-4 OE | WT | HSP-4 OE |  |  |
| # of Animals | 76 | 99 | 83 | 96 |  |  |  | 115 | 117 | 101 | 128 |  |  |  | 93 | 122 | 94 | 118 |  |  |
| Median Survival | 22 | 18 | 24 | 18 |  |  |  | 25 | 20 | 23 | 20 |  |  |  | 21 | 18 | 23 | 19 |  |  |
| **Fig. 4A/Fig. S4B** |  |  |  |  |  |  |  |  |  |  |  |  |  |  |  |  |  |  |  |  |
| **#1*** | WT | HSP-3 OE |  |  |  |  | **#2** | WT | HSP-3 OE |  |  |  |  | **#3** | WT | HSP-3 OE |  |  |  |  |
| # of Animals | 97 | 60 |  |  |  |  |  | 97 | 80 |  |  |  |  |  | 89 | 87 |  |  |  |  |
| Median Survival | 20 | 26 |  |  |  |  |  | 17 | 21 |  |  |  |  |  | 21 | 23 |  |  |  |  |
| **Fig. 4B/Fig. S5B** |  |  |  |  |  |  |  |  |  |  |  |  |  |  |  |  |  |  |  |  |
| **#1*** | WT | HSP-4 OE |  |  |  |  | **#2** | WT | HSP-4 OE |  |  |  |  | **#3** | WT | HSP-4 OE |  |  |  |  |
| # of Animals | 87 | 101 |  |  |  |  |  | 97 | 88 |  |  |  |  |  | 76 | 94 |  |  |  |  |
| Median Survival | 24 | 24 |  |  |  |  |  | 25 | 25 |  |  |  |  |  | 21 | 21 |  |  |  |  |
| **Fig. 4C/Fig. S3B** |  |  |  |  |  |  |  |  |  |  |  |  |  |  |  |  |  |  |  |  |
| **#1*** | *pos-1* | *hsp-3* | *hsp-4* |  |  |  | **#2** | *pos-1* | *hsp-3* | *hsp-4* |  |  |  | **#3** | *pos-1* | *hsp-3* | *hsp-4* |  |  |  |
| # of Animals | 74 | 81 | 74 |  |  |  |  | 89 | 80 | 53 |  |  |  |  | 66 | 81 | 34 |  |  |  |
| Median Survival | 23 | 14 | 17 |  |  |  |  | 25 | 14 | 16 |  |  |  |  | 21 | 16 | 19 |  |  |  |
| **Fig. 4D/Fig. S4C** |  |  |  |  |  |  |  |  |  |  |  |  |  |  |  |  |  |  |  |  |
| **#1*** | WT | HSP-3 OE |  |  |  |  | **#2** | WT | HSP-3 OE |  |  |  |  | **#3** | WT | HSP-3 OE |  |  |  |  |
| # of Animals | 41 | 77 |  |  |  |  |  | 117 | 86 |  |  |  |  |  | 80 | 70 |  |  |  |  |
| Median Survival | 17 | 13 |  |  |  |  |  | 16 | 12 |  |  |  |  |  | 16 | 12 |  |  |  |  |
| **Fig. 4E/Fig. S5C** |  |  |  |  |  |  |  |  |  |  |  |  |  |  |  |  |  |  |  |  |
| **#1*** | WT | HSP-4 OE |  |  |  |  | **#2** | WT | HSP-4 OE |  |  |  |  | **#3** | WT | HSP-4 OE |  |  |  |  |
| # of Animals | 110 | 81 |  |  |  |  |  | 127 | 137 |  |  |  |  |  | 138 | 184 |  |  |  |  |
| Median Survival | 16 | 12 |  |  |  |  |  | 13 | 13 |  |  |  |  |  | 17 | 15 |  |  |  |  |
| **Fig. 4F/Fig. S3C** |  |  |  |  |  |  |  |  |  |  |  |  |  |  |  |  |  |  |  |  |
| **#1*** | *pos-1* | *hsp-3* | *hsp-4* |  |  |  | **#2** | *pos-1* | *hsp-3* | *hsp-4* |  |  |  | **#3** | *pos-1* | *hsp-3* | *hsp-4* |  |  |  |
| # of Animals | 88 | 60 | 63 |  |  |  |  | 74 | 97 | 101 |  |  |  |  | 66 | 34 | 56 |  |  |  |
| Median Survival | 23 | 23 | 16 |  |  |  |  | 20 | 20 | 16 |  |  |  |  | 18 | 21 | 16 |  |  |  |
| **Fig. 4G/Fig. S3D** |  |  |  |  |  |  |  |  |  |  |  |  |  |  |  |  |  |  |  |  |
| **#1*** | *pos-1* | *hsp-3* | *hsp-4* |  |  |  | **#2** | *pos-1* | *hsp-3* | *hsp-4* |  |  |  | **#3** | *pos-1* | *hsp-3* | *hsp-4* |  |  |  |
| # of Animals | 66 | 81 | 34 |  |  |  |  | 124 | 132 | 133 |  |  |  |  | 52 | 115 | 104 |  |  |  |
| Median Survival | 21 | 16 | 19 |  |  |  |  | 26 | 14 | 26 |  |  |  |  | 25 | 17 | 22 |  |  |  |
| **Fig. 4H/Fig. S7C** |  |  |  |  |  |  |  |  |  |  |  |  |  |  |  |  |  |  |  |  |
| **#1*** | *pos-1* | *hsp-3* | *hsp-4* |  |  |  | **#2** | *pos-1* | *hsp-3* | *hsp-4* |  |  |  | **#3** | *pos-1* | *hsp-3* | *hsp-4* |  |  |  |
| # of Animals | 61 | 47 | 34 |  |  |  |  | 65 | 53 | 38 |  |  |  |  | 58 | 47 | 49 |  |  |  |
| Median Survival | 15 | 15 | 5 |  |  |  |  | 15 | 13 | 5 |  |  |  |  | 14 | 14 | 5 |  |  |  |
| **Fig. 4I/Fig. S7D** |  |  |  |  |  |  |  |  |  |  |  |  |  |  |  |  |  |  |  |  |
| **#1*** | *pos-1* | *hsp-3* | *hsp-4* |  |  |  | **#2** | *pos-1* | *hsp-3* | *hsp-4* |  |  |  | **#3** | *pos-1* | *hsp-3* | *hsp-4* |  |  |  |
| # of Animals | 117 | 108 | 103 |  |  |  |  | 105 | 119 | 117 |  |  |  |  | 94 | 54 | 69 |  |  |  |
| Median Survival | 18 | 18 | 14 |  |  |  |  | 21 | 17 | 15 |  |  |  |  | 19 | 17 | 15 |  |  |  |
| **Fig. 4J/Fig. S7E** |  |  |  |  |  |  |  |  |  |  |  |  |  |  |  |  |  |  |  |  |
| **#1*** | *pos-1* | *hsp-3* | *hsp-4* |  |  |  | **#2** | *pos-1* | *hsp-3* | *hsp-4* |  |  |  | **#3** | *pos-1* | *hsp-3* | *hsp-4* |  |  |  |
| # of Animals | 39 | 70 | 58 |  |  |  |  | 57 | 59 | 56 |  |  |  |  | 63 | 80 | 71 |  |  |  |
| Median Survival | 20 | 10 | 12 |  |  |  |  | 14 | 11 | 11 |  |  |  |  | 15 | 11 | 11 |  |  |  |
| **Fig. 4K/Fig. S7B** |  |  |  |  |  |  |  |  |  |  |  |  |  |  |  |  |  |  |  |  |
| **#1*** | *pos-1* | *hsp-3* | *hsp-4* |  |  |  | **#2** | *pos-1* | *hsp-3* | *hsp-4* |  |  |  | **#3** | *pos-1* | *hsp-3* | *hsp-4* |  |  |  |
| # of Animals | 19 | 59 | 38 |  |  |  |  | 87 | 90 | 83 |  |  |  |  | 73 | 105 | 117 |  |  |  |
| Median Survival | 21 | 19 | 15 |  |  |  |  | 21 | 21 | 19 |  |  |  |  | 19 | 19 | 14 |  |  |  |
| **Fig. 4L/Fig. S7A** |  |  |  |  |  |  |  |  |  |  |  |  |  |  |  |  |  |  |  |  |
| **#1*** | *pos-1* | *hsp-3* | *hsp-4* |  |  |  | **#2** | *pos-1* | *hsp-3* | *hsp-4* |  |  |  | **#3** | *pos-1* | *hsp-3* | *hsp-4* |  |  |  |
| # of Animals | 69 | 49 | 54 |  |  |  |  | 52 | 70 | 49 |  |  |  |  | 52 | 65 | 45 |  |  |  |
| Median Survival | 22 | 9 | 14 |  |  |  |  | 23 | 9 | 11 |  |  |  |  | 19 | 9 | 11 |  |  |  |
| **Fig. 6F/Fig. S13A** |  |  |  |  |  |  |  |  |  |  |  |  |  |  |  |  |  |  |  |  |
| **#1*** | *pos-1* | *hsp-4* | *C18E9.2* | *hsp-4/C18E9.2* |  |  | **#2** | *pos-1* | *hsp-4* | *C18E9.2* | *hsp-4/C18E9.2* |  |  | **#2** | *pos-1* | *hsp-4* | *C18E9.2* | *hsp-4/C18E9.2* |  |  |
| # of Animals | 87 | 63 | 53 | 75 |  |  |  | 71 | 57 | 68 | 67 |  |  |  | 86 | 108 | 75 | 63 |  |  |
| Median Survival | 20 | 14 | 22 | 12 |  |  |  | 29 | 18 | 29 | 16 |  |  |  | 23 | 14 | 25 | 13 |  |  |

Table S4

| **Fig. 5A+B/S8** | **#1*** | 20 °C | 20 °C | 20 °C | 25 °C | 25 °C | 25 °C |  |  |
| --- | --- | --- | --- | --- | --- | --- | --- | --- | --- |
|  |  | *pos-1* | *hsp-3* | *hsp-4* | *pos-1* | *hsp-3* | *hsp-4* |  |  |
|  | # Paralyzed | 23 | 98 | 69 | 132 | 130 | 65 |  |  |
|  | # Not Paralyzed | 131 | 0 | 110 | 0 | 0 | 42 |  |  |
|  | Median Paralysis | Undef. | 7.5 | Undef. | 8 | 8 | 12 |  |  |
|  | **#2** | 20 °C | 20 °C | 20 °C | 25 °C | 25 °C | 25 °C |  |  |
|  |  | *pos-1* | *hsp-3* | *hsp-4* | *pos-1* | *hsp-3* | *hsp-4* |  |  |
|  | # Paralyzed | 38 | 119 | 76 | 131 | 139 | 95 |  |  |
|  | # Not Paralyzed | 78 | 0 | 61 | 0 | 0 | 22 |  |  |
|  | Median Paralysis | Undef. | 8 | 14 | 8 | 8 | 12 |  |  |
|  | **#3** | 20 °C | 20 °C | 20 °C | 25 °C | 25 °C | 25 °C |  |  |
|  |  | *pos-1* | *hsp-3* | *hsp-4* | *pos-1* | *hsp-3* | *hsp-4* |  |  |
|  | # Paralyzed | 15 | 72 | 43 | 88 | 56 | 32 |  |  |
|  | # Not Paralyzed | 65 | 0 | 12 | 0 | 0 | 5 |  |  |
|  | Median Paralysis | Undef. | 10 | 11 | 8 | 8 | 13 |  |  |
| **Fig. 5C/S9A** | **#1*** | 20 °C | 20 °C | 25 °C | 25 °C |  |  |  |  |
|  |  | WT | HSP-4 OE | WT | HSP-4 OE |  |  |  |  |
|  | # Paralyzed | 21 | 25 | 77 | 126 |  |  |  |  |
|  | # Not Paralyzed | 66 | 103 | 0 | 0 |  |  |  |  |
|  | Median Paralysis | Undef. | Undef. | 10 | 6 |  |  |  |  |
|  | **#2** | 20 °C | 20 °C | 25 °C | 25 °C |  |  |  |  |
|  |  | WT | HSP-4 OE | WT | HSP-4 OE |  |  |  |  |
|  | # Paralyzed | 27 | 34 | 102 | 117 |  |  |  |  |
|  | # Not Paralyzed | 79 | 67 | 0 | 0 |  |  |  |  |
|  | Median Paralysis | Undef. | Undef. | 10 | 4 |  |  |  |  |
|  | **#3** | 20 °C | 20 °C | 25 °C | 25 °C |  |  |  |  |
|  |  | WT | HSP-4 OE | WT | HSP-4 OE |  |  |  |  |
|  | # Paralyzed | 69 | 67 | 0 | 0 |  |  |  |  |
|  | # Not Paralyzed | 79 | 67 | 0 | 0 |  |  |  |  |
|  | Median Paralysis | Undef. | Undef. | 11 | 5 |  |  |  |  |
| **Fig. 5C/S9A** | **#1*** | 20 °C | 20 °C | 25 °C | 25 °C |  |  |  |  |
|  |  | WT | HSP-4 OE | WT | HSP-4 OE |  |  |  |  |
|  | # Paralyzed | 21 | 25 | 77 | 126 |  |  |  |  |
|  | # Not Paralyzed | 66 | 103 | 0 | 0 |  |  |  |  |
|  | Median Paralysis | Undef. | Undef. | 10 | 6 |  |  |  |  |
|  | **#2** | 20 °C | 20 °C | 25 °C | 25 °C |  |  |  |  |
|  |  | WT | HSP-4 OE | WT | HSP-4 OE |  |  |  |  |
|  | # Paralyzed | 27 | 34 | 102 | 117 |  |  |  |  |
|  | # Not Paralyzed | 79 | 67 | 0 | 0 |  |  |  |  |
|  | Median Paralysis | Undef. | Undef. | 10 | 4 |  |  |  |  |
|  | **#3** | 20 °C | 20 °C | 25 °C | 25 °C |  |  |  |  |
|  |  | WT | HSP-4 OE | WT | HSP-4 OE |  |  |  |  |
|  | # Paralyzed | 69 | 67 | 0 | 0 |  |  |  |  |
|  | # Not Paralyzed | 79 | 67 | 0 | 0 |  |  |  |  |
|  | Median Paralysis | Undef. | Undef. | 11 | 5 |  |  |  |  |
| **Fig. 5D/S9B** | **#1*** | 20 °C | 20 °C | 25 °C | 25 °C |  |  |  |  |
|  |  | WT | HSP-3 OE | WT | HSP-3 OE |  |  |  |  |
|  | # Paralyzed | 28 | 37 | 110 | 92 |  |  |  |  |
|  | # Not Paralyzed | 72 | 66 | 0 | 0 |  |  |  |  |
|  | Median Paralysis | Undef. | Undef. | 11 | 11 |  |  |  |  |
|  | **#2** | 20 °C | 20 °C | 25 °C | 25 °C |  |  |  |  |
|  |  | WT | HSP-3 OE | WT | HSP-3 OE |  |  |  |  |
|  | # Paralyzed | 19 | 17 | 103 | 60 |  |  |  |  |
|  | # Not Paralyzed | 70 | 34 | 0 | 0 |  |  |  |  |
|  | Median Paralysis | Undef. | Undef. | 9 | 9 |  |  |  |  |
|  | **#3** | 20 °C | 20 °C | 25 °C | 25 °C |  |  |  |  |
|  |  | WT | HSP-3 OE | WT | HSP-3 OE |  |  |  |  |
|  | # Paralyzed | 35 | 37 | 126 | 137 |  |  |  |  |
|  | # Not Paralyzed | 83 | 55 | 0 | 0 |  |  |  |  |
|  | Median Paralysis | Undef. | Undef. | 9 | 11 |  |  |  |  |
| **Fig. 5D/S9B** | **#1*** | 20 °C | 20 °C | 25 °C | 25 °C |  |  |  |  |
|  |  | WT | HSP-3 OE | WT | HSP-3 OE |  |  |  |  |
|  | # Paralyzed | 28 | 37 | 110 | 92 |  |  |  |  |
|  | # Not Paralyzed | 72 | 66 | 0 | 0 |  |  |  |  |
|  | Median Paralysis | Undef. | Undef. | 11 | 11 |  |  |  |  |
|  | **#2** | 20 °C | 20 °C | 25 °C | 25 °C |  |  |  |  |
|  |  | WT | HSP-3 OE | WT | HSP-3 OE |  |  |  |  |
|  | # Paralyzed | 19 | 17 | 103 | 60 |  |  |  |  |
|  | # Not Paralyzed | 70 | 34 | 0 | 0 |  |  |  |  |
|  | Median Paralysis | Undef. | Undef. | 9 | 9 |  |  |  |  |
|  | **#3** | 20 °C | 20 °C | 25 °C | 25 °C |  |  |  |  |
|  |  | WT | HSP-3 OE | WT | HSP-3 OE |  |  |  |  |
|  | # Paralyzed | 35 | 37 | 126 | 137 |  |  |  |  |
|  | # Not Paralyzed | 83 | 55 | 0 | 0 |  |  |  |  |
|  | Median Paralysis | Undef. | Undef. | 9 | 11 |  |  |  |  |
| **Fig. 5D/S9B** | **#1*** | 20 °C | 20 °C | 25 °C | 25 °C |  |  |  |  |
|  |  | WT | HSP-3 OE | WT | HSP-3 OE |  |  |  |  |
|  | # Paralyzed | 28 | 37 | 110 | 92 |  |  |  |  |
|  | # Not Paralyzed | 72 | 66 | 0 | 0 |  |  |  |  |
|  | Median Paralysis | Undef. | Undef. | 11 | 11 |  |  |  |  |
|  | **#2** | 20 °C | 20 °C | 25 °C | 25 °C |  |  |  |  |
|  |  | WT | HSP-3 OE | WT | HSP-3 OE |  |  |  |  |
|  | # Paralyzed | 19 | 17 | 103 | 60 |  |  |  |  |
|  | # Not Paralyzed | 70 | 34 | 0 | 0 |  |  |  |  |
|  | Median Paralysis | Undef. | Undef. | 9 | 9 |  |  |  |  |
|  | **#3** | 20 °C | 20 °C | 25 °C | 25 °C |  |  |  |  |
|  |  | WT | HSP-3 OE | WT | HSP-3 OE |  |  |  |  |
|  | # Paralyzed | 35 | 37 | 126 | 137 |  |  |  |  |
|  | # Not Paralyzed | 83 | 55 | 0 | 0 |  |  |  |  |
|  | Median Paralysis | Undef. | Undef. | 9 | 11 |  |  |  |  |
| **Fig. 5E/S9C** | **#1*** | 20 °C | 20 °C | 25 °C | 25 °C |  |  |  |  |
|  |  | WT | HSP-4 OE | WT | HSP-4 OE |  |  |  |  |
|  | # Paralyzed | 106 | 81 | 93 | 91 |  |  |  |  |
|  | # Not Paralyzed | 0 | 0 | 0 | 0 |  |  |  |  |
|  | Median Paralysis | 9 | 9 | 9 | 9 |  |  |  |  |
|  | **#2** | 20 °C | 20 °C | 25 °C | 25 °C |  |  |  |  |
|  | # Paralyzed | WT | HSP-4 OE | WT | HSP-4 OE |  |  |  |  |
|  | # Not Paralyzed | 81 | 126 | 114 | 70 |  |  |  |  |
|  | Median Paralysis | 0 | 0 | 0 | 0 |  |  |  |  |
|  |  | 10 | 10 | 10 | 10 |  |  |  |  |
|  | **#3** | 20 °C | 20 °C | 25 °C | 25 °C |  |  |  |  |
|  |  | WT | HSP-4 OE | WT | HSP-4 OE |  |  |  |  |
|  | # Paralyzed | 57 | 151 | 50 | 153 |  |  |  |  |
|  | # Not Paralyzed | 0 | 0 | 0 | 0 |  |  |  |  |
|  | Median Paralysis | 9 | 9 | 9 | 9 |  |  |  |  |
| **Fig. 5E/S9C** | **#1*** | 20 °C | 20 °C | 25 °C | 25 °C |  |  |  |  |
|  |  | WT | HSP-4 OE | WT | HSP-4 OE |  |  |  |  |
|  | # Paralyzed | 106 | 81 | 93 | 91 |  |  |  |  |
|  | # Not Paralyzed | 0 | 0 | 0 | 0 |  |  |  |  |
|  | Median Paralysis | 9 | 9 | 9 | 9 |  |  |  |  |
|  | **#2** | 20 °C | 20 °C | 25 °C | 25 °C |  |  |  |  |
|  |  | WT | HSP-4 OE | WT | HSP-4 OE |  |  |  |  |
|  | # Paralyzed | 81 | 126 | 114 | 70 |  |  |  |  |
|  | # Not Paralyzed | 0 | 0 | 0 | 0 |  |  |  |  |
|  | Median Paralysis | 10 | 10 | 10 | 10 |  |  |  |  |
|  | **#3** | 20 °C | 20 °C | 25 °C | 25 °C |  |  |  |  |
|  |  | WT | HSP-4 OE | WT | HSP-4 OE |  |  |  |  |
|  | # Paralyzed | 57 | 151 | 50 | 153 |  |  |  |  |
|  | # Not Paralyzed | 0 | 0 | 0 | 0 |  |  |  |  |
|  | Median Paralysis | 9 | 9 | 9 | 9 |  |  |  |  |
| **Fig. 5E/S9C** | **#1*** | 20 °C | 20 °C | 25 °C | 25 °C |  |  |  |  |
|  |  | WT | HSP-4 OE | WT | HSP-4 OE |  |  |  |  |
|  | # Paralyzed | 106 | 81 | 93 | 91 |  |  |  |  |
|  | # Not Paralyzed | 0 | 0 | 0 | 0 |  |  |  |  |
|  | Median Paralysis | 9 | 9 | 9 | 9 |  |  |  |  |
|  | **#2** | 20 °C | 20 °C | 25 °C | 25 °C |  |  |  |  |
|  |  | WT | HSP-4 OE | WT | HSP-4 OE |  |  |  |  |
|  | # Paralyzed | 81 | 126 | 114 | 70 |  |  |  |  |
|  | # Not Paralyzed | 0 | 0 | 0 | 0 |  |  |  |  |
|  | Median Paralysis | 10 | 10 | 10 | 10 |  |  |  |  |
|  | **#3** | 20 °C | 20 °C | 25 °C | 25 °C |  |  |  |  |
|  |  | WT | HSP-4 OE | WT | HSP-4 OE |  |  |  |  |
|  | # Paralyzed | 57 | 151 | 50 | 153 |  |  |  |  |
|  | # Not Paralyzed | 0 | 0 | 0 | 0 |  |  |  |  |
|  | Median Paralysis | 9 | 9 | 9 | 9 |  |  |  |  |
| **Fig. 6G+H/S13B** | **#1*** | 20 °C | 20 °C | 20 °C | 20 °C | 25 °C | 25 °C | 25 °C | 25 °C |
|  |  | *pos-1* | *hsp-4* | *C18E9.2* | *hsp-4/C18E9.2* | *pos-1* | *hsp-4* | *C18E9.2* | *hsp-4/C18E9.2* |
|  | # Paralyzed | 17 | 99 | 14 | 83 | 98 | 92 | 83 | 97 |
|  | # Not Paralyzed | 90 | 23 | 76 | 51 | 0 | 20 | 15 | 0 |
|  | Median Paralysis | Undef. | 12 | Undef. | 14 | 12 | 14 | 12 | 12 |
|  | **#2** | 20 °C | 20 °C | 20 °C | 20 °C | 25 °C | 25 °C | 25 °C | 25 °C |
|  |  | *pos-1* | *hsp-4* | *C18E9.2* | *hsp-4/C18E9.2* | *pos-1* | *hsp-4* | *C18E9.2* | *hsp-4/C18E9.2* |
|  | # Paralyzed | 21 | 44 | 21 | 67 | 77 | 53 | 76 | 69 |
|  | # Not Paralyzed | 66 | 11 | 72 | 48 | 0 | 4 | 3 | 4 |
|  | Median Paralysis | Undef. | 15 | Undef. | 15 | 10 | 12 | 12 | 12 |
|  | **#3** | 20 °C | 20 °C | 20 °C | 20 °C | 25 °C | 25 °C | 25 °C | 25 °C |
|  |  | *pos-1* | *hsp-4* | *C18E9.2* | *hsp-4/C18E9.2* | *pos-1* | *hsp-4* | *C18E9.2* | *hsp-4/C18E9.2* |
|  | # Paralyzed | 14 | 27 | 17 | 32 | 73 | 51 | 61 | 33 |
|  | # Not Paralyzed | 51 | 37 | 41 | 34 | 0 | 21 | 4 | 0 |
|  | Median Paralysis | Undef. | Undef. | Undef. | Undef. | 11 | 13 | 11 | 11 |

Table S5

| *hsp-3* siRNA - Upregulated | | | | | *hsp-3* siRNA - Downegulated | | | | |
| --- | --- | --- | --- | --- | --- | --- | --- | --- | --- |
| **Gene** | **logFC** | **logCPM** | **F** | **PValue** | **Gene** | **logFC** | **logCPM** | **F** | **PValue** |
| *abf-4* | 4.868975411 | -2.318114971 | 7.6902571 | 0.01754739 | *abu-9* | -6.9898895 | -1.9800123 | 8.79635062 | 0.01704855 |
| *aip-1* | 2.01064538 | 6.281732911 | 49.31841056 | 7.67E-06 | *col-123* | -1.9246734 | -0.3679438 | 12.0610284 | 0.00394328 |
| *warf-1* | 2.000746882 | 4.486062768 | 22.95288231 | 0.000324519 | *cpr-2* | -2.4434223 | -2.1258835 | 8.55079086 | 0.01150558 |
| *WBGene00000191* | 1.627491251 | 1.973772606 | 11.78741515 | 0.004259997 | *dhs-15* | -1.6918951 | 0.48625788 | 11.0098874 | 0.00533377 |
| *cyp-14A5* | 1.665544089 | 4.674006483 | 61.19594505 | 2.32E-06 | *ech-9* | -1.8794485 | 1.28790355 | 8.8468199 | 0.01046144 |
| *ckb-2* | 2.323798837 | 7.055811516 | 95.73376093 | 1.75E-07 | *grl-27* | -1.8670156 | 1.58253465 | 26.2765852 | 0.00017552 |
| *dhs-9* | 1.634444197 | 6.12913709 | 111.2599831 | 7.11E-08 | *hsp-3* | -3.7358584 | 9.62286441 | 66.2969663 | 1.51E-06 |
| *WBGene00001091* | 1.545467861 | 8.516343946 | 52.53229927 | 5.37E-06 | *lbp-8* | -2.0139146 | 2.06158288 | 13.675123 | 0.00254538 |
| *far-3* | 3.374904915 | 7.589868735 | 90.31592961 | 2.53E-07 | *pqn-91* | -1.5104238 | 0.39106285 | 10.8535251 | 0.00558587 |
| *gem-4* | 1.719828431 | 4.369642044 | 17.37466618 | 0.001036726 | *smf-3* | -1.7520778 | 2.42400407 | 24.9227202 | 0.00022325 |
| *gst-38* | 1.534020649 | 5.036865288 | 25.70408966 | 0.000194881 | *spp-4* | -1.6225231 | 3.5539164 | 22.1852662 | 0.00037354 |
| *his-25* | 1.622827387 | -0.335308732 | 6.03170466 | 0.028360822 | *srh-1* | -1.7883227 | -1.8268082 | 4.76900701 | 0.04726179 |
| *hsp-16.1* | 3.596652032 | -0.450353387 | 10.53933977 | 0.006155126 | *sri-40* | -1.6436446 | 1.50901105 | 7.22624332 | 0.0182149 |
| *hsp-16.2* | 4.200291301 | 6.462412632 | 10.72392372 | 0.005824446 | *srv-13* | -2.5280136 | -1.9955499 | 4.97063345 | 0.04342175 |
| *hsp-16.11* | 3.09310012 | 9.169827711 | 6.767888807 | 0.021516022 | *str-41* | -3.0078509 | -1.8155339 | 11.4888059 | 0.00463976 |
| *hsp-16.41* | 4.362029546 | 7.552686199 | 9.112702052 | 0.009595774 | *tbh-1* | -2.3945397 | 3.62846081 | 5.49120468 | 0.03514052 |
| *hsp-16.48* | 3.118354224 | 6.646645191 | 6.970592059 | 0.019976636 | *vit-1* | -2.2584925 | 6.61798559 | 5.5035216 | 0.03496741 |
| *hsp-17* | 2.794190137 | 7.027714569 | 91.39363769 | 2.33E-07 | *vit-3* | -4.9217435 | 6.62861605 | 5.66370356 | 0.03280497 |
| *hsp-70* | 2.816831474 | 7.791404392 | 6.475310107 | 0.023989563 | *vit-4* | -5.2444412 | 6.66488231 | 6.06264043 | 0.02806797 |
| *nas-3* | 1.915000948 | 1.046415962 | 10.35058796 | 0.006502285 | *C10C5.4* | -1.5090605 | 5.12971534 | 10.6510309 | 0.00595251 |
| *pqn-97* | 1.51961583 | -0.593412792 | 4.867080096 | 0.045345878 | *C26G2.2* | -1.5457494 | 3.63279838 | 5.06927019 | 0.04172962 |
| *pqn-98* | 2.134881996 | -0.625483152 | 5.248994763 | 0.0387318 | *cyp-25A1* | -1.5615732 | 0.95724698 | 5.92957123 | 0.0295528 |
| *rab-11.2* | 4.259513096 | 4.152709529 | 20.82600397 | 0.000493408 | *sdz-6* | -6.6177507 | -0.5863521 | 9.58981251 | 0.00893286 |
| *rrf-2* | 2.434289518 | 6.667065273 | 56.64138293 | 3.62E-06 | *pals-38* | -1.6356982 | 0.56301976 | 6.47867059 | 0.02391814 |
| *skr-5* | 2.312006217 | 3.32999632 | 33.11226355 | 5.86E-05 | *bgnt-1.3* | -4.4133551 | -1.4677885 | 4.91566401 | 0.0477293 |
| *sri-36* | 3.74557197 | -0.05722095 | 25.24284613 | 0.000210738 | *WBGene00009230* | -2.8489993 | -0.3269326 | 17.7364008 | 0.00094959 |
| *sri-39* | 3.209225039 | -0.316330612 | 10.90827449 | 0.006036954 | *F29D10.2* | -1.6691157 | -1.0661672 | 7.20068392 | 0.0183463 |
| *sri-74* | 2.550965052 | -2.113683159 | 4.950669647 | 0.045289116 | *F36D1.8* | -2.8452616 | -0.2134833 | 30.4581518 | 8.78E-05 |
| *fbxa-199* | 1.701675373 | 0.623562506 | 10.90180658 | 0.005506581 | *oac-20* | -4.179017 | 3.30992647 | 46.6658424 | 1.03E-05 |
| *srx-111* | 2.762759189 | -2.251873176 | 6.026176049 | 0.028421549 | *clec-24* | -3.5278136 | -0.678648 | 20.0438916 | 0.00057573 |
| *str-144* | 2.726780902 | -0.951794978 | 12.67778366 | 0.003324151 | *clec-28* | -2.4906268 | 1.18792963 | 35.6224071 | 4.08E-05 |
| *arrd-3* | 3.394383643 | 1.675101447 | 35.07869424 | 4.43E-05 | *clec-33* | -3.1528125 | -0.3404786 | 15.7539607 | 0.00151086 |
| *tba-7* | 1.798129986 | 6.018070125 | 137.7294222 | 1.93E-08 | *F49C12.7* | -1.8150414 | 3.18620294 | 54.1846501 | 4.54E-06 |
| *tbb-6* | 5.040562933 | 7.113539526 | 112.4318247 | 6.85E-08 | *F55G11.2* | -1.829499 | 2.71043474 | 29.8537373 | 9.67E-05 |
| *tsp-1* | 1.817806414 | 2.155083938 | 9.511804619 | 0.00844766 | *H25K10.1* | -1.8653058 | 1.97420587 | 27.2279335 | 0.00014898 |
| *tsp-2* | 1.513387786 | 2.589777862 | 8.683292738 | 0.011038895 | *K01D12.8* | -2.1059781 | 3.16421759 | 43.9868648 | 1.39E-05 |
| *pals-26* | 2.249434333 | 1.554116124 | 13.05769622 | 0.003011368 | *cyp-14A2* | -2.2815941 | 2.12495995 | 14.6517181 | 0.00198961 |
| *WBGene00007181* | 1.674007913 | 0.378709677 | 14.47028472 | 0.002073197 | *WBGene00010753* | -2.261379 | -1.4464233 | 5.11178234 | 0.04096908 |
| *C04F12.1* | 2.164545643 | 6.131884396 | 89.86109762 | 2.54E-07 | *M04C9.4* | -1.7481981 | 2.1271637 | 29.8482666 | 9.67E-05 |
| *rnh-1.3* | 2.234747791 | 2.850679486 | 37.84802319 | 3.01E-05 | *cyp-14A4* | -1.9512256 | 2.62737807 | 30.8700734 | 8.24E-05 |
| *C08E8.4* | 2.791108794 | 6.604062957 | 32.93889711 | 6.08E-05 | *chil-23* | -1.6907918 | 1.71657982 | 9.02641052 | 0.00986727 |
| *fbxa-98* | 1.57491208 | 4.499792884 | 9.443116174 | 0.008633383 | *ugt-30* | -1.7561046 | 2.51719429 | 14.9866247 | 0.00182371 |
| *fbxc-58* | 2.197246775 | 4.872333019 | 29.42866201 | 0.000104507 | *T05E12.3* | -2.189148 | -1.2235268 | 7.64136428 | 0.01569206 |
| *C31H5.7* | 2.952892841 | -1.374846893 | 12.49820924 | 0.003492006 | *T13F3.6* | -1.5543197 | 4.57640506 | 8.03935314 | 0.01370404 |
| *C32H11.9* | 2.691997271 | 4.115677062 | 12.99560882 | 0.003062067 | *chil-25* | -1.886331 | -0.7127229 | 6.16678573 | 0.0269233 |
| *fipr-23* | 2.582037377 | -0.908851281 | 5.61866517 | 0.033349247 | *sysm-1* | -3.794604 | 0.97215434 | 18.9782901 | 0.00086709 |
| *C45B11.2* | 3.040742398 | 2.231312382 | 27.930549 | 0.00013324 | *clec-144* | -3.4965479 | -1.2494549 | 22.6928961 | 0.00033849 |
| *fbxa-141* | 1.928921004 | 2.174120761 | 12.40530136 | 0.003592367 | *comt-2* | -1.5405255 | -1.5819794 | 6.56626918 | 0.02314596 |
| *oac-14* | 1.944172202 | 7.553951143 | 20.29570187 | 0.000550098 | *clec-4* | -1.5830284 | 2.34391312 | 7.09622345 | 0.01908716 |
| *ifas-2* | 2.603532477 | -0.639370044 | 5.083240111 | 0.041489293 | *nhr-234* | -1.8273258 | 2.18956103 | 8.0336696 | 0.0137307 |
| *F11D11.3* | 4.018540745 | 3.1378401 | 40.73757438 | 2.09E-05 | *Y40H7A.10* | -1.7938901 | 3.8328264 | 13.6504313 | 0.0025732 |
| *F14F8.8* | 3.961909848 | -0.601727534 | 7.561620927 | 0.017162809 | *clec-247* | -1.5758366 | -1.3387954 | 5.29738306 | 0.03795237 |
| *F15B9.6* | 4.630056842 | 4.934525845 | 68.65711588 | 1.23E-06 | *pals-30* | -1.7538281 | -1.1787808 | 5.08579869 | 0.04139403 |
| *F19B2.5* | 1.577255748 | 8.904539383 | 7.034272562 | 0.019519861 | *cest-2.3* | -1.7342173 | -0.9116671 | 6.30338306 | 0.02555571 |
| *F20G2.1* | 2.052795174 | 3.173109905 | 30.40176307 | 8.86E-05 | *ZK218.4* | -3.7743898 | -1.9325726 | 14.2925786 | 0.00216926 |
| *F47B8.4* | 2.09531466 | 2.619286297 | 8.487172757 | 0.011780988 | *clec-61* | -2.5776003 | 2.06984462 | 15.3575033 | 0.00166962 |
| *F53B2.8* | 1.715490579 | 7.598588128 | 18.41890725 | 0.000820707 | *ZK1025.2* | -1.5624887 | 3.98670409 | 5.487592 | 0.03519149 |
| *fipr-26* | 4.086320249 | -0.484429839 | 4.952493871 | 0.045255383 | *WBGene00015077* | -2.2615777 | 1.67002252 | 10.6363372 | 0.00595928 |
| *gmd-2* | 2.867101119 | 1.6765979 | 14.86703679 | 0.001885027 | *clec-10* | -1.5304405 | 4.95423889 | 24.5408802 | 0.00024033 |
| *mpst-2* | 4.399256148 | -1.812227427 | 6.40936786 | 0.028915211 | *C04G6.2* | -1.592456 | 4.11344647 | 6.53783999 | 0.02343409 |
| *cdr-4* | 3.703921268 | 9.34060361 | 66.95217157 | 1.42E-06 | *C07G1.7* | -2.1827072 | 0.78641203 | 10.2816136 | 0.00664005 |
| *K10G4.3* | 2.647219884 | 2.192493771 | 76.8742759 | 6.33E-07 | *C09B8.4* | -1.7234828 | 2.71703162 | 7.45080777 | 0.01681552 |
| *M01G12.7* | 4.807510504 | 2.216293872 | 34.00508289 | 5.20E-05 | *C18A11.4* | -2.6626553 | -1.1632283 | 23.303885 | 0.00030121 |
| *M01G12.9* | 1.85440437 | 6.962501169 | 57.90467919 | 3.15E-06 | *C18H7.1* | -4.0509306 | -0.8885721 | 12.3143982 | 0.00367405 |
| *M163.5* | 1.77254369 | -1.684751921 | 4.65332905 | 0.049646181 | *C30G12.2* | -1.6739556 | 3.51167019 | 19.0635188 | 0.00070882 |
| *arrd-11* | 4.440165923 | 2.311400743 | 91.7308634 | 2.25E-07 | *ilys-2* | -2.7986465 | 1.75869956 | 13.1375183 | 0.00329991 |
| *zip-6* | 2.644757284 | 0.832955315 | 17.12820639 | 0.001093501 | *C53B7.2* | -1.7182467 | 2.45819373 | 6.2246971 | 0.02637627 |
| *R12H7.4* | 1.626859827 | -0.803684472 | 7.643311226 | 0.015681368 | *D1014.6* | -5.9704136 | 3.15621297 | 20.0065427 | 0.00058416 |
| *cyp-13A5* | 1.651241581 | 5.393597859 | 4.771065747 | 0.047274382 | *D1014.7* | -4.647236 | 3.13797815 | 17.952097 | 0.00091013 |
| *T23F11.6* | 1.72976552 | 0.741643322 | 9.32614415 | 0.008935953 | *oac-12* | -1.852063 | -0.8742543 | 4.92518348 | 0.04425416 |
| *zip-10* | 2.434777207 | 5.048322593 | 26.20210892 | 0.000179396 | *F07E5.7* | -2.0728004 | 0.40752713 | 10.9257393 | 0.00547387 |
| *arrd-8* | 2.207922388 | -0.38056419 | 11.23578538 | 0.004992449 | *F14H12.7* | -1.8022816 | -1.3060189 | 4.95419523 | 0.04372065 |
| *Y26D4A.3* | 2.64227567 | -0.065649012 | 8.715312984 | 0.010904467 | *pud-2.1* | -4.8403522 | 5.16787247 | 13.421226 | 0.00273327 |
| *pals-13* | 3.849684874 | -2.13821717 | 5.971996332 | 0.029024801 | *pud-4* | -5.714297 | 2.93629892 | 57.8481962 | 3.22E-06 |
| *Y37H2A.11* | 4.417262907 | -2.153068509 | 7.036004255 | 0.023414761 | *pud-1.1* | -5.2780525 | 0.7040734 | 48.7580745 | 1.90E-05 |
| *Y38E10A.22* | 2.596049464 | 7.135180633 | 210.703282 | 1.35E-09 | *pud-2.2* | -5.3346383 | 6.75731475 | 71.1871797 | 1.01E-06 |
| *Y38H6C.9* | 1.618275466 | 1.259949361 | 15.1048567 | 0.001768056 | *pud-3* | -6.9694638 | 3.55227436 | 78.984085 | 9.92E-07 |
| *Y54G11A.4* | 1.531723335 | 3.823433942 | 46.13924898 | 1.08E-05 | *F23F12.3* | -2.0153957 | 2.53131975 | 31.5052468 | 7.47E-05 |
| *Y75B12B.3* | 1.843508281 | -1.105476792 | 8.005316435 | 0.013833115 | *folt-2* | -1.6531494 | 8.54636977 | 10.0934602 | 0.00704776 |
| *fbxa-30* | 2.959693627 | 2.617459957 | 25.51112338 | 0.000202608 | *F42A10.7* | -1.7575645 | 3.47167112 | 21.9823261 | 0.00038871 |
| *fbxa-50* | 1.538817677 | -0.486808094 | 7.021823025 | 0.019570697 | *WBGene00018448* | -4.8508703 | 0.11326008 | 67.0645882 | 1.38E-06 |
| *clec-60* | 2.64537488 | 3.483639466 | 10.35081633 | 0.006515535 | *WBGene00018449* | -3.2511027 | -1.404927 | 17.5301261 | 0.00099493 |
| *ZK896.1* | 2.466011157 | 3.521908323 | 38.20775368 | 2.87E-05 | *WBGene00018450* | -3.2511027 | -1.404927 | 17.5301261 | 0.00099493 |
| *ZK970.7* | 1.709529666 | 5.123797259 | 14.6098265 | 0.002010732 | *WBGene00018451* | -3.2511027 | -1.404927 | 17.5301261 | 0.00099493 |
| *B0348.2* | 3.173544727 | 1.923395746 | 51.95764445 | 5.70E-06 | *F45D11.15* | -3.8317203 | 1.24413377 | 6.07693999 | 0.02918875 |
| *dod-20* | 1.910714926 | 2.949000539 | 23.7349127 | 0.000277753 | *F45D11.16* | -3.8317203 | 1.24413377 | 6.07693999 | 0.02918875 |
| *cyp-35A1* | 1.632617645 | 2.217943957 | 23.57035763 | 0.000286453 | *F47B7.4* | -3.4905042 | -2.0655156 | 6.43860423 | 0.02547529 |
| *C06E4.6* | 2.460693813 | 1.212791514 | 18.84263872 | 0.000743506 | *F48G7.5* | -2.5951338 | 3.10246002 | 10.4109801 | 0.00639792 |
| *irg-1* | 1.965067536 | 4.441307806 | 22.15309634 | 0.000378718 | *F49D11.3* | -3.0200139 | -0.9804452 | 9.32685735 | 0.0089421 |
| *WBGene00015596* | 3.217937857 | -0.389958848 | 10.27279052 | 0.006651038 | *oac-32* | -2.6248797 | 0.10250739 | 32.6568196 | 6.27E-05 |
| *WBGene00015597* | 2.909220877 | 3.319353763 | 84.08821356 | 3.76E-07 | *F54D10.8* | -2.2888718 | 0.43297117 | 20.5579288 | 0.00062684 |
| *fbxa-163* | 4.373011025 | 3.109920872 | 36.36214805 | 3.73E-05 | *F56A4.2* | -1.709734 | 7.38047537 | 18.6913133 | 0.00077322 |
| *fbxa-165* | 4.693871971 | -0.602489783 | 10.9730645 | 0.005926587 | *F56A4.3* | -5.6210279 | 5.24017392 | 4.80374597 | 0.04999579 |
| *fbxa-158* | 2.409155459 | 2.873323268 | 18.16403565 | 0.000868207 | *H01M10.2* | -1.8611443 | -1.8280983 | 5.87351548 | 0.03016023 |
| *C14C6.3* | 3.199913918 | -0.40189837 | 8.706261942 | 0.013958184 | *WBGene00019208* | -1.9824151 | 0.03328993 | 7.65878104 | 0.01563022 |
| *nhr-155* | 3.623367027 | 0.895037292 | 26.63791278 | 0.000166332 | *K01A2.4* | -1.8292328 | 0.52094923 | 9.11930062 | 0.00954969 |
| *C14C6.6* | 3.2545837 | 0.95718496 | 8.081275191 | 0.014413872 | *cyp-35A5* | -1.541129 | 4.11588672 | 18.2396824 | 0.00085032 |
| *WBGene00015761* | 3.646933383 | 0.41060129 | 6.866390813 | 0.023207687 | *clec-43* | -1.837131 | 0.62026841 | 5.97113076 | 0.02907938 |
| *C14C6.8* | 3.209334424 | 0.610892659 | 6.911075219 | 0.020414872 | *WBGene00019934* | -1.5576067 | -0.6306365 | 8.15554303 | 0.01314145 |
| *pals-32* | 2.359235166 | 2.800266569 | 32.76706919 | 6.17E-05 | *hacd-1* | -1.5513023 | 4.75149274 | 22.62205 | 0.00034526 |
| *fbxa-12* | 3.356720124 | 1.302787209 | 40.63413756 | 2.09E-05 | *ugt-53* | -2.4708936 | 3.03128335 | 48.8970573 | 7.91E-06 |
| *WBGene00016526* | 2.931348901 | -0.14350986 | 21.20674071 | 0.000453594 | *clec-53* | -1.5672735 | 2.654718 | 7.14619709 | 0.01874625 |
| *fbxc-5* | 1.651299058 | 4.146964706 | 32.66391664 | 6.26E-05 | *math-38* | -2.1933215 | 0.13695564 | 14.435543 | 0.00209158 |
| *fbxc-4* | 1.587832139 | 4.135734743 | 29.0614671 | 0.000109818 | *dach-1* | -3.73706 | 2.09359134 | 46.0060479 | 1.10E-05 |
| *fbxc-2* | 1.587832139 | 4.135734743 | 29.0614671 | 0.000109818 | *T10B5.8* | -2.2187964 | 3.00005834 | 32.7636405 | 6.17E-05 |
| *fbxc-1* | 1.651299058 | 4.146964706 | 32.66391664 | 6.26E-05 | *math-42* | -1.5308347 | 1.43376449 | 19.786535 | 0.00060765 |
| *C49G7.7* | 3.50330426 | 4.988340921 | 30.66620443 | 8.59E-05 | *clec-218* | -1.7426039 | 4.5659774 | 8.03333993 | 0.01373224 |
| *C49G7.10* | 1.927549007 | 6.017976376 | 17.4674084 | 0.001015087 | *clec-118* | -1.8776605 | -1.2661411 | 14.7623235 | 0.00192579 |
| *D1022.5* | 2.147366096 | 0.071774604 | 5.913754953 | 0.029690104 | *clec-209* | -1.7097301 | 7.2401654 | 15.0283158 | 0.00181084 |
| *EEED8.12* | 2.212006668 | -0.610808052 | 5.407168555 | 0.036299941 | *pud-1.2* | -5.5113138 | 8.19530346 | 56.3900298 | 3.71E-06 |
| *numr-1* | 2.837336844 | 2.330328988 | 7.955752034 | 0.014102298 | *Y27F2A.9* | -1.9045263 | -0.8312195 | 9.61414819 | 0.00815602 |
| *F13A2.2* | 2.540021612 | -1.162017666 | 8.999833106 | 0.009926427 | *comt-4* | -2.1852748 | 0.84371715 | 12.050973 | 0.00395443 |
| *F14F9.3* | 2.152614976 | 5.030558965 | 14.46683552 | 0.002084791 | *clec-70* | -2.8062262 | -1.6212221 | 11.328006 | 0.00536222 |
| *F22E5.6* | 4.906560106 | 5.396821986 | 120.5749914 | 4.49E-08 | *clec-71* | -1.5357965 | 0.09730042 | 6.47408823 | 0.02395936 |
| *F25A2.1* | 1.585925851 | 1.906815327 | 17.44514645 | 0.001014332 | *plep-1* | -1.9816568 | 0.62049236 | 20.2059512 | 0.00055663 |
| *math-28* | 2.498269775 | -1.361349619 | 5.472175021 | 0.035361446 | *WBGene00022209* | -2.5622259 | -1.8589405 | 5.36204858 | 0.03838546 |
| *fbxb-53* | 4.747246336 | -1.070842703 | 22.40183217 | 0.0003581 | *srt-13* | -2.4344046 | -2.2427047 | 5.01471392 | 0.04263206 |
| *F41B4.3* | 1.768050368 | 3.461860737 | 32.35991875 | 6.56E-05 | *ZK105.6* | -2.1221319 | -0.7675644 | 9.49352271 | 0.00847266 |
| *WBGene00018325* | 1.762702071 | 1.028909879 | 4.879105026 | 0.04517032 | *F11D5.7* | -2.0697841 | 0.43108884 | 7.70381032 | 0.01751779 |
| *fbxa-182* | 3.216032925 | 4.566287169 | 111.3057298 | 7.09E-08 | *H39E23.3* | -3.7011831 | -2.1197003 | 5.34969158 | 0.04475912 |
| *F48G7.13* | 3.150981179 | 1.278091469 | 20.00477933 | 0.000584371 | *WBGene00044503* | -1.8724915 | -1.2840197 | 8.21424341 | 0.01288218 |
| *F57B9.3* | 4.075695327 | 4.480082359 | 64.38011785 | 1.78E-06 | *F23G4.1* | -1.8000607 | -0.5617615 | 13.8434779 | 0.00243583 |
| *fbxc-3* | 1.587832139 | 4.135734743 | 29.0614671 | 0.000109818 | *folt-3* | -2.5857592 | -1.60453 | 14.6757588 | 0.00196818 |
| *K06H6.1* | 3.450389802 | 1.238645463 | 7.632489531 | 0.017932034 | *T19H5.6* | -1.9856628 | 0.5023092 | 14.7448818 | 0.00193424 |
| *K06H6.2* | 3.230570712 | 2.072808723 | 5.178253166 | 0.041362276 | *C04G6.13* | -2.0866561 | -2.3255916 | 5.95913544 | 0.0291702 |
| *K06H6.4* | 2.782273762 | -1.646214457 | 5.221356887 | 0.039199114 | *T27A10.8* | -4.3000344 | -0.5450452 | 5.18886901 | 0.0397241 |
| *WBGene00019454* | 3.61503429 | -2.186617751 | 9.844115793 | 0.01131314 | *T14G12.12* | -2.0257727 | -0.4527414 | 20.7787367 | 0.00049471 |
| *K09D9.1* | 2.842242071 | 5.27899087 | 32.88637048 | 6.13E-05 | *BE0003N10.6* | -5.1912027 | 4.82047596 | 16.0026516 | 0.00142812 |
| *pgph-1* | 5.848838635 | -0.811223703 | 6.674334077 | 0.023428104 | *F54D8.10* | -2.6757308 | -1.2960764 | 8.54021359 | 0.01154616 |
| *fil-2* | 1.571639286 | -1.059188579 | 5.845213778 | 0.031811137 | *WBGene00304827* | -2.562085 | -1.7034195 | 4.91945851 | 0.04436031 |
| *M60.7* | 1.72041712 | 3.054061603 | 37.97669301 | 2.96E-05 |  |  |  |  |  |
| *R03H10.6* | 2.426650973 | 3.147587696 | 39.4245725 | 2.45E-05 |  |  |  |  |  |
| *T03F1.6* | 1.623593555 | 2.180652338 | 10.9925836 | 0.005367038 |  |  |  |  |  |
| *T05A8.7* | 2.407265637 | -2.074489516 | 5.868074683 | 0.030224465 |  |  |  |  |  |
| *lgc-1* | 1.794275987 | -0.341220311 | 15.80010166 | 0.001491542 |  |  |  |  |  |
| *phat-5* | 1.561467642 | 0.926638698 | 4.756355476 | 0.047515863 |  |  |  |  |  |
| *T08E11.1* | 2.671672163 | 0.831044484 | 10.7441706 | 0.005789443 |  |  |  |  |  |
| *WBGene00020364* | 2.330078111 | 0.864892333 | 38.11173733 | 2.91E-05 |  |  |  |  |  |
| *cnp-3* | 1.744656303 | 5.659045584 | 21.13013843 | 0.000463944 |  |  |  |  |  |
| *T24E12.5* | 1.508825455 | 5.567578031 | 15.48095008 | 0.001619971 |  |  |  |  |  |
| *bath-25* | 1.926652623 | 0.769432953 | 5.68737782 | 0.032498883 |  |  |  |  |  |
| *WBGene00020981* | 1.696156673 | 4.77010592 | 9.908086576 | 0.007462455 |  |  |  |  |  |
| *W09G12.7* | 2.418812288 | 5.016078359 | 8.647432517 | 0.01117043 |  |  |  |  |  |
| *clec-121* | 4.120053988 | -0.537082972 | 5.463360621 | 0.036882392 |  |  |  |  |  |
| *Y34F4.4* | 2.054683168 | 4.023362518 | 12.87362295 | 0.003164579 |  |  |  |  |  |
| *comt-3* | 2.045129483 | 6.598764589 | 26.86466427 | 0.00015997 |  |  |  |  |  |
| *Y41D4B.15* | 1.579068862 | 0.522856269 | 7.629569282 | 0.015757008 |  |  |  |  |  |
| *fbxa-48* | 3.445071562 | -0.495975908 | 34.32476774 | 4.91E-05 |  |  |  |  |  |
| *fbxa-66* | 3.544810782 | 2.050605095 | 86.31640862 | 3.22E-07 |  |  |  |  |  |
| *Y58A7A.3* | 1.655236911 | 7.246741991 | 34.6455419 | 4.71E-05 |  |  |  |  |  |
| *Y58A7A.4* | 2.55196236 | 2.864428693 | 22.2979435 | 0.000368172 |  |  |  |  |  |
| *Y58A7A.5* | 1.885875161 | 6.307958002 | 31.41181706 | 7.65E-05 |  |  |  |  |  |
| *Y71G12B.2* | 1.739449633 | 3.008501621 | 40.45981624 | 2.14E-05 |  |  |  |  |  |
| *fbxa-25* | 2.752434 | 0.925369207 | 26.6839643 | 0.000163541 |  |  |  |  |  |
| *fbxa-138* | 2.041733499 | 0.102670652 | 19.73801215 | 0.000613896 |  |  |  |  |  |
| *Y94H6A.10* | 2.715268913 | 9.596041326 | 48.90785682 | 7.94E-06 |  |  |  |  |  |
| *Y102A11A.9* | 2.152603239 | 4.113963048 | 38.4359266 | 2.78E-05 |  |  |  |  |  |
| *fbxa-35* | 1.70535635 | -0.98905673 | 8.201648899 | 0.012937303 |  |  |  |  |  |
| *fbxa-36* | 2.248475207 | -0.661440618 | 6.514021092 | 0.023616513 |  |  |  |  |  |
| *WBGene00022545* | 2.321013459 | 1.765737362 | 12.36346749 | 0.003638599 |  |  |  |  |  |
| *sdz-35* | 4.858593477 | 5.194006589 | 78.43531514 | 5.76E-07 |  |  |  |  |  |
| *ZC239.14* | 3.166118284 | 3.131825123 | 78.4879089 | 5.62E-07 |  |  |  |  |  |
| *ZK177.9* | 4.90179488 | -1.829198385 | 10.66229239 | 0.007174369 |  |  |  |  |  |
| *ZK488.5* | 3.011200201 | 0.214291107 | 8.799533338 | 0.010624733 |  |  |  |  |  |
| *ZK488.6* | 3.68064976 | -0.154791407 | 6.82984913 | 0.022156265 |  |  |  |  |  |
| *ZK1240.1* | 4.548041601 | -0.387620784 | 29.40614585 | 0.000103858 |  |  |  |  |  |
| *ZK1240.8* | 3.020818343 | -0.858132951 | 17.30890237 | 0.001046357 |  |  |  |  |  |
| *Y75B8A.39* | 3.296401743 | 2.820637535 | 35.15087178 | 4.39E-05 |  |  |  |  |  |
| *F40F12.9* | 1.87970979 | 2.739588874 | 18.38134823 | 0.000822449 |  |  |  |  |  |
| *Y6G8.5* | 2.83862627 | 1.041448553 | 17.55969907 | 0.000994069 |  |  |  |  |  |
| *Y71G12B.32* | 1.893412406 | 1.910420671 | 18.51172 | 0.000799199 |  |  |  |  |  |
| *K10G6.5* | 3.710443383 | -1.035647338 | 9.573126421 | 0.008279416 |  |  |  |  |  |
| *Y82E9BL.18* | 1.681420974 | 3.346493782 | 16.72739229 | 0.001197272 |  |  |  |  |  |
| *T08A9.13* | 1.666659633 | 0.271583677 | 10.56805808 | 0.006082595 |  |  |  |  |  |
| *ZC21.10* | 3.854474968 | 0.562718453 | 41.28188152 | 1.93E-05 |  |  |  |  |  |
| *F41E6.15* | 2.537806853 | -1.063545507 | 8.411338061 | 0.012054147 |  |  |  |  |  |
| *B0205.13* | 1.68188246 | 4.831128425 | 19.64890379 | 0.000629695 |  |  |  |  |  |
| *B0205.14* | 2.915950399 | 2.540165804 | 35.50054643 | 4.15E-05 |  |  |  |  |  |
| *C01G10.17* | 1.994904985 | 0.742214776 | 6.468885269 | 0.024047505 |  |  |  |  |  |
| *B0303.16* | 2.289710437 | -1.383281467 | 12.57623133 | 0.0034179 |  |  |  |  |  |
| *WBGene00045188* | 2.519155216 | -1.495016953 | 4.683962142 | 0.049001162 |  |  |  |  |  |
| *fbxc-12* | 2.130926781 | -1.470445296 | 4.926165063 | 0.04574516 |  |  |  |  |  |
| *M01B2.13* | 1.652594037 | 2.184054334 | 13.1554276 | 0.002921196 |  |  |  |  |  |
| *eol-1* | 1.801272822 | 1.532676901 | 12.80417452 | 0.003211594 |  |  |  |  |  |
| *C25F9.11* | 1.956856754 | 3.955044162 | 12.63642592 | 0.003375485 |  |  |  |  |  |
| *C25F9.12* | 1.533037172 | 1.70190023 | 5.022880714 | 0.042539524 |  |  |  |  |  |
| *Y43F8B.15* | 1.808198683 | 2.348388004 | 15.62906756 | 0.001554632 |  |  |  |  |  |
| *Y37H2A.14* | 2.453090263 | 6.432705491 | 31.13766935 | 7.98E-05 |  |  |  |  |  |
| *F33H12.7* | 3.216175312 | 6.516671115 | 54.62433049 | 4.41E-06 |  |  |  |  |  |
| *WBGene00077629* | 2.356445942 | 1.623683703 | 35.78398616 | 3.99E-05 |  |  |  |  |  |
| *Y26D4A.21* | 2.635258976 | -0.364623067 | 26.98042622 | 0.000155412 |  |  |  |  |  |
| *C39B5.14* | 1.714511398 | 0.520391745 | 9.982109726 | 0.007271638 |  |  |  |  |  |
| *C33D9.13* | 1.836079027 | 4.335014185 | 49.32219518 | 7.55E-06 |  |  |  |  |  |
| *B0462.5* | 1.982320138 | 2.923324004 | 27.03405614 | 0.000153992 |  |  |  |  |  |
| *F19B10.13* | 2.921190005 | 0.348981551 | 16.02459396 | 0.001413164 |  |  |  |  |  |
| *ZC239.22* | 1.593767487 | 1.138255216 | 8.839841723 | 0.010458321 |  |  |  |  |  |
| *pals-20* | 2.372283398 | -0.662102557 | 14.37867295 | 0.002122093 |  |  |  |  |  |
| *K01A6.8* | 2.061313448 | -0.639088112 | 7.787726703 | 0.01491151 |  |  |  |  |  |
| *Y43F8B.25* | 1.974554745 | 1.068544684 | 17.1164072 | 0.001093612 |  |  |  |  |  |
| *anr-16* | 2.432172397 | -0.873371461 | 14.98522112 | 0.001821389 |  |  |  |  |  |
| *anr-32* | 1.71875802 | 0.442717881 | 12.93427537 | 0.003100306 |  |  |  |  |  |
| *Y14H12A.4* | 1.564760585 | 1.976994097 | 33.43855284 | 5.59E-05 |  |  |  |  |  |
| *C54F6.18* | 2.177981252 | 0.185136721 | 5.766136984 | 0.031504494 |  |  |  |  |  |
| *WBGene00269338* | 3.108122404 | 1.914694175 | 21.84412669 | 0.000402397 |  |  |  |  |  |
| *Y41C4A.32* | 3.869668774 | 9.918314264 | 208.9030404 | 1.43E-09 |  |  |  |  |  |
| *H05L14.3* | 2.162316109 | -1.08571704 | 11.57528389 | 0.004958712 |  |  |  |  |  |
| *F19B10.14* | 2.202947543 | -0.852155223 | 5.33339451 | 0.037400764 |  |  |  |  |  |
| *WBGene00271786* | 3.150828678 | -0.632067247 | 8.078328002 | 0.013522857 |  |  |  |  |  |
| *H04D03.7* | 2.22788195 | 1.694437914 | 15.97639272 | 0.001429584 |  |  |  |  |  |

Table S6

| *hsp-4* siRNA - Upregulated | | | | | *hsp-4* siRNA - Downregulated | | | | |
| --- | --- | --- | --- | --- | --- | --- | --- | --- | --- |
| **Gene** | **logFC** | **logCPM** | **F** | **PValue** | **Gene** | **logFC** | **logCPM** | **F** | **PValue** |
| *acr-18* | 1.624232525 | 1.928524235 | 15.88313583 | 0.001461982 | *abu-9* | -6.9898895 | -1.9800123 | 9.34486433 | 0.01478665 |
| *warf-1* | 3.694171745 | 4.486062768 | 70.15874439 | 1.09E-06 | *amt-4* | -1.8631544 | 5.62447874 | 44.4037565 | 1.33E-05 |
| *WBGene00000191* | 3.564013981 | 1.973772606 | 53.96990634 | 4.64E-06 | *aqp-1* | -2.3270465 | 4.22616353 | 35.2332014 | 4.36E-05 |
| *bre-1* | 2.01519665 | 7.529133704 | 118.4116512 | 4.87E-08 | *asm-2* | -3.5726133 | 3.79539294 | 41.2985177 | 1.95E-05 |
| *bro-1* | 1.704971166 | 1.566247158 | 19.59507948 | 0.000632722 | *asm-3* | -2.5525693 | 1.50491294 | 11.3216411 | 0.00488655 |
| *ckb-2* | 4.336628809 | 7.055811516 | 282.3855136 | 2.08E-10 | *asp-1* | -2.236157 | 11.5624655 | 23.9834048 | 0.00026519 |
| *cnd-1* | 1.518103901 | -1.06707699 | 8.488331815 | 0.01174761 | *asp-2* | -2.1161302 | 8.90166347 | 32.034565 | 6.95E-05 |
| *crn-4* | 1.699838023 | 2.441527316 | 35.03067981 | 4.44E-05 | *asp-6* | -1.8019337 | 10.0610335 | 43.8259328 | 1.41E-05 |
| *cul-6* | 1.838675724 | 4.529542037 | 34.21619179 | 4.99E-05 | *cpr-2* | -1.6651043 | -2.1258835 | 5.01194044 | 0.04268124 |
| *daf-14* | 1.516450989 | 5.281853432 | 66.66423 | 1.43E-06 | *cpr-4* | -2.1878162 | 8.73338025 | 20.8526574 | 0.00049074 |
| *dnj-28* | 1.635293712 | 5.528326103 | 48.68787532 | 8.10E-06 | *cpr-5* | -1.7641238 | 9.36013599 | 8.46501452 | 0.01186842 |
| *dsc-4* | 1.512836519 | 8.45248183 | 39.83396996 | 2.32E-05 | *dhs-2* | -1.531004 | 5.00284523 | 37.0808479 | 3.34E-05 |
| *fis-2* | 2.404150372 | 5.686299006 | 97.80009942 | 1.54E-07 | *dhs-14* | -1.9405357 | 5.04998262 | 20.9886385 | 0.00047739 |
| *fkh-3* | 5.624340117 | -1.585258758 | 6.127567322 | 0.028640005 | *ech-9* | -2.2177965 | 1.28790355 | 11.9953282 | 0.00403194 |
| *gcy-13* | 2.540199543 | -1.759240173 | 12.54130506 | 0.003450846 | *gfi-1* | -1.7341967 | 9.77196609 | 28.5366401 | 0.00011967 |
| *gem-4* | 2.510332376 | 4.369642044 | 35.50711967 | 4.20E-05 | *gst-10* | -2.0175862 | 7.43963067 | 27.0693087 | 0.00015447 |
| *gpa-6* | 1.801407156 | 3.457900182 | 19.40729784 | 0.000658476 | *hsp-4* | -2.0809817 | 9.04243988 | 74.0507476 | 7.86E-07 |
| *gst-9* | 1.894674094 | 1.889063255 | 8.025305992 | 0.013770033 | *ins-35* | -2.8830709 | -0.3965523 | 23.2443029 | 0.00030463 |
| *gst-38* | 1.857379943 | 5.036865288 | 37.11102385 | 3.34E-05 | *lbp-8* | -4.4283102 | 2.06158288 | 41.5723614 | 1.86E-05 |
| *his-17* | 2.422678655 | -0.871930928 | 6.171634843 | 0.026916671 | *lys-1* | -2.082528 | 8.06952107 | 51.562867 | 5.94E-06 |
| *ins-34* | 4.432479728 | -1.498614389 | 50.50399896 | 6.65E-06 | *lys-4* | -3.0935427 | 7.44815519 | 68.5480078 | 1.25E-06 |
| *kgb-2* | 2.074068345 | 3.901135995 | 53.8658791 | 4.69E-06 | *lys-7* | -1.8903533 | 7.05825099 | 13.8575651 | 0.0024378 |
| *lec-8* | 1.540300631 | 8.299732228 | 32.4464741 | 6.47E-05 | *lys-8* | -1.8234993 | 7.06526533 | 18.2562083 | 0.00085067 |
| *mig-21* | 1.582837042 | 0.997245207 | 9.315695189 | 0.008965836 | *lys-10* | -4.8645897 | 1.56791284 | 7.35809908 | 0.01840473 |
| *nhr-3* | 1.653909482 | 5.851413269 | 44.90138028 | 1.24E-05 | *nuc-1* | -2.3556654 | 6.26551036 | 82.1389241 | 4.34E-07 |
| *nhr-6* | 1.646862187 | 3.705041094 | 9.390847473 | 0.008777888 | *pcp-1* | -1.8079019 | 6.80418269 | 34.4543244 | 4.87E-05 |
| *nhr-117* | 1.516031375 | 3.718871441 | 30.65843511 | 8.51E-05 | *pcp-3* | -1.5573362 | 7.35076112 | 69.2979094 | 1.15E-06 |
| *nsf-1* | 1.941907385 | 7.659426718 | 144.463894 | 1.44E-08 | *pmp-5* | -1.725267 | 7.43611684 | 50.7642827 | 6.47E-06 |
| *rab-11.2* | 7.644277048 | 4.152709529 | 68.57917611 | 1.24E-06 | *trpl-5* | -1.7178899 | 0.47807461 | 9.046339 | 0.00977776 |
| *rrf-2* | 4.061099616 | 6.667065273 | 138.3358002 | 1.94E-08 | *smf-3* | -2.4891975 | 2.42400407 | 47.1666669 | 9.59E-06 |
| *sid-2* | 2.510809444 | 6.050443419 | 111.1552152 | 7.15E-08 | *spp-1* | -1.5733898 | 4.13710437 | 18.8189147 | 0.00074891 |
| *sip-1* | 1.818810202 | 7.510914222 | 11.22325019 | 0.005028172 | *spp-4* | -2.6957191 | 3.5539164 | 56.2040455 | 3.71E-06 |
| *sir-2.3* | 4.083094674 | 2.198671653 | 65.98465667 | 1.52E-06 | *spp-11* | -1.9641421 | -1.6180664 | 7.21971521 | 0.01822136 |
| *skr-3* | 1.647663173 | 7.169750453 | 39.67958333 | 2.37E-05 | *spp-16* | -1.5212584 | 3.77384893 | 22.4359239 | 0.00035574 |
| *skr-5* | 1.639954789 | 3.32999632 | 17.19015883 | 0.001075222 | *srx-68* | -1.8789388 | -1.3398504 | 12.0170528 | 0.0039923 |
| *srh-116* | 3.211340328 | -2.558422345 | 6.576166264 | 0.024227361 | *str-7* | -3.1867244 | 2.26588807 | 37.5363618 | 3.14E-05 |
| *sri-36* | 3.960476554 | -0.05722095 | 28.9330593 | 0.000112139 | *str-168* | -2.6706761 | 0.24438117 | 34.0228381 | 5.13E-05 |
| *sri-39* | 3.118681901 | -0.316330612 | 10.33722904 | 0.007121177 | *tag-10* | -1.8303551 | 6.66136563 | 12.3617601 | 0.00364032 |
| *sri-67* | 4.72629458 | -0.922421927 | 8.729445421 | 0.015320711 | *vit-1* | -3.0520997 | 6.61798559 | 9.40096519 | 0.0087497 |
| *sri-70* | 4.068395177 | 0.216168155 | 45.47390874 | 1.16E-05 | *vit-3* | -5.6850406 | 6.62861605 | 6.99822483 | 0.01977691 |
| *sri-74* | 2.779426382 | -2.113683159 | 6.252423982 | 0.02728745 | *vit-4* | -4.9363686 | 6.66488231 | 5.57779591 | 0.0339445 |
| *srp-2* | 1.607633404 | 3.151797493 | 21.01304156 | 0.000471692 | *wrt-7* | -2.4669862 | -1.2590036 | 6.33407514 | 0.02525972 |
| *srp-7* | 4.654096095 | 9.756704862 | 357.6292886 | 4.53E-11 | *C08B6.2* | -2.2052525 | -1.5305755 | 4.99571009 | 0.04297041 |
| *srp-8* | 1.743425678 | 3.028120234 | 34.95137261 | 4.49E-05 | *ugt-22* | -1.9713986 | 7.97072187 | 49.2031443 | 7.65E-06 |
| *srw-86* | 2.658434403 | 1.688335438 | 29.07013962 | 0.000109664 | *cbl-1* | -2.3227805 | 4.97186779 | 39.9445429 | 2.32E-05 |
| *fbxa-199* | 2.196941976 | 0.623562506 | 18.43944395 | 0.000811994 | *hrg-7* | -3.4363835 | 7.13399911 | 46.8784467 | 1.01E-05 |
| *str-82* | 4.433081649 | -1.385076882 | 25.80574833 | 0.000418668 | *ugt-23* | -2.0264939 | 5.88942242 | 84.622779 | 3.62E-07 |
| *str-124* | 5.492083974 | -1.618932497 | 12.92931056 | 0.003482619 | *cest-9.1* | -1.5748926 | 4.57156903 | 41.5941961 | 1.86E-05 |
| *str-245* | 3.705383331 | -1.659077296 | 10.76979108 | 0.00572653 | *cest-7* | -2.2152225 | 1.6971791 | 26.5074141 | 0.00016861 |
| *syx-2* | 1.555782056 | 4.944138457 | 33.44272806 | 5.58E-05 | *C31H5.6* | -1.690242 | 4.3379687 | 51.2321926 | 6.15E-06 |
| *arrd-3* | 3.727045091 | 1.675101447 | 41.82454817 | 1.81E-05 | *cyp-25A1* | -2.7192214 | 0.95724698 | 16.040841 | 0.00141514 |
| *tba-7* | 2.833240964 | 6.018070125 | 319.1791485 | 9.47E-11 | *C55A1.6* | -2.8276904 | -0.6584651 | 7.06860984 | 0.01924144 |
| *tbb-6* | 4.886166412 | 7.113539526 | 107.1197429 | 9.17E-08 | *ttr-44* | -2.5224317 | 5.88648122 | 125.674751 | 3.39E-08 |
| *tsp-1* | 2.225000588 | 2.155083938 | 14.16909079 | 0.002249345 | *cest-3* | -2.3986393 | 2.95680342 | 42.7366352 | 1.61E-05 |
| *tsp-2* | 2.858603935 | 2.589777862 | 29.49796714 | 0.00010335 | *clec-17* | -2.8905152 | 3.10647861 | 13.7742559 | 0.00249124 |
| *ubc-23* | 2.327388291 | 4.170878924 | 173.4203187 | 4.62E-09 | *F01D5.1* | -3.6361839 | 2.48991383 | 37.1999925 | 3.33E-05 |
| *zmp-1* | 1.756988704 | 2.844024053 | 74.63108466 | 7.51E-07 | *F01D5.2* | -4.7451835 | 0.84426881 | 72.1718397 | 9.11E-07 |
| *plin-1* | 2.194249364 | 8.978094654 | 170.0495338 | 5.22E-09 | *F01D5.3* | -4.5966904 | 2.8941251 | 39.255225 | 2.53E-05 |
| *B0457.6* | 1.868241175 | 4.212900073 | 48.61864874 | 8.16E-06 | *F01D5.5* | -1.8896576 | 4.72173379 | 8.95447359 | 0.01010044 |
| *C06B3.7* | 2.042773607 | 3.877768773 | 25.75458004 | 0.000193148 | *acox-1.4* | -2.2660137 | 5.58945475 | 90.6294365 | 2.42E-07 |
| *C06C3.7* | 1.877782252 | -0.953116363 | 9.645120993 | 0.008076944 | *clec-227* | -4.0377681 | 1.4576759 | 29.6825523 | 0.00010034 |
| *C08E8.4* | 3.82099309 | 6.604062957 | 56.72625001 | 3.59E-06 | *clec-57* | -1.8787441 | 5.39217642 | 32.915606 | 6.10E-05 |
| *fbxa-98* | 2.65961167 | 4.499792884 | 25.45383198 | 0.000204684 | *clec-54* | -2.1994634 | 2.87087135 | 43.0862206 | 1.55E-05 |
| *fbxc-58* | 3.325403955 | 4.872333019 | 62.69427819 | 2.06E-06 | *F09B12.3* | -1.9933273 | 7.75817747 | 14.0231291 | 0.00233546 |
| *C11E4.8* | 2.961661115 | 2.395694863 | 73.90484213 | 7.95E-07 | *F09C8.1* | -1.5345385 | 5.61446288 | 31.5757328 | 7.41E-05 |
| *pals-6* | 2.270671056 | 3.867779724 | 50.43106234 | 6.70E-06 | *cpt-5* | -2.1839405 | 3.95611369 | 33.1923423 | 5.81E-05 |
| *C23H4.6* | 3.134973011 | 4.64488968 | 138.227687 | 1.89E-08 | *stdh-2* | -3.5025745 | 1.74760288 | 41.5110294 | 1.88E-05 |
| *fbxa-82* | 1.889982696 | 3.104009749 | 24.07083298 | 0.000260921 | *ctsa-1.2* | -1.8000709 | 8.10562028 | 6.50304421 | 0.02374131 |
| *C25F9.5* | 1.740687287 | 5.180862439 | 47.66249282 | 9.07E-06 | *F14D7.6* | -1.630942 | 4.27994001 | 25.1274955 | 0.00021515 |
| *C31H5.7* | 2.619920036 | -1.374846893 | 9.541037897 | 0.008346258 | *cyp-13A12* | -1.9380177 | 2.93221224 | 10.9590708 | 0.00543255 |
| *C32H11.9* | 2.544002168 | 4.115677062 | 11.69717966 | 0.004386846 | *cyp-35D1* | -3.0260376 | 2.85832124 | 56.7334989 | 3.53E-06 |
| *dod-21* | 1.950941506 | 4.930421698 | 4.708556741 | 0.048544707 | *clec-42* | -1.6611524 | 3.95638724 | 6.49005515 | 0.0238572 |
| *C34C12.4* | 1.971922664 | 5.624770775 | 132.756179 | 2.42E-08 | *F17B5.1* | -1.7533753 | 4.27812915 | 8.48637964 | 0.0117841 |
| *zip-5* | 2.230140431 | 4.265397385 | 24.38773249 | 0.000248144 | *cth-1* | -2.9972503 | 7.51191728 | 34.9175333 | 4.56E-05 |
| *C44H9.4* | 3.430306769 | 5.208263157 | 126.3978674 | 3.30E-08 | *asah-2* | -3.1762725 | 5.18432033 | 91.4000203 | 2.36E-07 |
| *C45B11.2* | 4.355674562 | 2.231312382 | 54.36260669 | 4.51E-06 | *thn-1* | -3.3447998 | -1.3959957 | 22.1923367 | 0.00037302 |
| *gale-1* | 3.220517627 | 8.89472988 | 111.7181828 | 6.93E-08 | *thn-2* | -4.6314872 | 2.63715209 | 136.977419 | 2.00E-08 |
| *fbxa-141* | 3.53690756 | 2.174120761 | 39.6030587 | 2.41E-05 | *cyp-37B1* | -3.6316285 | 3.49082015 | 41.7102669 | 1.85E-05 |
| *mfb-1* | 1.772541207 | 4.148613677 | 60.79639881 | 2.40E-06 | *WBGene00009230* | -5.0513566 | -0.3269326 | 36.4883357 | 3.62E-05 |
| *E04D5.4* | 1.576558593 | 2.256288878 | 8.737832228 | 0.010842279 | *clec-64* | -2.6054799 | 2.05269587 | 44.1243149 | 1.36E-05 |
| *ugt-44* | 1.761128656 | 6.99949451 | 11.28500505 | 0.004938739 | *F35E12.6* | -1.6235204 | 6.92884845 | 16.7173715 | 0.00120629 |
| *F01D5.7* | 2.108717184 | 1.601692172 | 24.26555063 | 0.000251703 | *oac-20* | -3.1181429 | 3.30992647 | 29.4579018 | 0.00010402 |
| *gfat-1* | 2.675499309 | 8.402494321 | 156.315181 | 8.83E-09 | *clec-166* | -1.9599839 | 4.39493815 | 64.2510433 | 1.76E-06 |
| *F08G2.5* | 2.563614958 | 3.765066167 | 32.2848442 | 6.70E-05 | *clec-165* | -1.6923914 | 1.87095399 | 30.7953682 | 8.33E-05 |
| *F11D11.3* | 4.368102573 | 3.1378401 | 47.50989158 | 9.37E-06 | *clec-169* | -2.2101727 | 3.74274615 | 42.7401034 | 1.61E-05 |
| *F13A7.11* | 2.040467276 | 1.991445989 | 19.53313199 | 0.000641088 | *argk-1* | -3.4707176 | 8.47997587 | 18.8005937 | 0.00075506 |
| *F15B9.6* | 3.736773309 | 4.934525845 | 47.59341522 | 9.28E-06 | *lipl-2* | -1.6896533 | 5.08480902 | 12.1938862 | 0.003814 |
| *F16B12.4* | 1.919166134 | 3.924709895 | 52.30347727 | 5.50E-06 | *WBGene00009838* | -2.2696321 | 0.05926314 | 18.0567094 | 0.00088381 |
| *F17C11.11* | 1.833343245 | 6.472401018 | 34.14794709 | 5.09E-05 | *clec-31* | -2.6364826 | 0.25163658 | 25.4815626 | 0.00020193 |
| *plp-2* | 2.013746372 | -1.338818072 | 10.24374697 | 0.006710206 | *clec-24* | -5.4312417 | -0.678648 | 32.2752232 | 6.64E-05 |
| *pals-14* | 2.51589719 | 2.608230816 | 20.54469515 | 0.000520648 | *clec-28* | -4.0331605 | 1.18792963 | 75.9854918 | 6.77E-07 |
| *F23B2.10* | 1.548251527 | 0.182989036 | 4.94179851 | 0.043947668 | *clec-33* | -4.5155569 | -0.3404786 | 25.6925357 | 0.00019506 |
| *F35E8.2* | 2.508868213 | 0.5187223 | 34.94642807 | 4.49E-05 | *clec-32* | -3.791174 | -2.0955333 | 13.8709745 | 0.00273965 |
| *WBGene00009428* | 2.47645099 | 2.326277007 | 27.68948269 | 0.000137788 | *F49C12.4* | -2.4289129 | 4.04804294 | 6.13106343 | 0.02733863 |
| *cbp-3* | 1.605374511 | 6.749081853 | 22.22901641 | 0.000373147 | *F49C12.7* | -3.2775039 | 3.18620294 | 154.136016 | 9.64E-09 |
| *fbxa-144* | 2.215591803 | -0.568107113 | 24.62097494 | 0.000235826 | *nspg-11* | -1.5708718 | 2.59085873 | 14.1593737 | 0.00224465 |
| *ipla-2* | 3.585786307 | 7.233620255 | 194.369778 | 2.25E-09 | *F49E12.10* | -1.7688243 | 1.77059354 | 9.16630792 | 0.00943149 |
| *F47B8.3* | 1.84382676 | 3.988048762 | 28.55416423 | 0.000119322 | *WBGene00009991* | -2.4344019 | 0.19512983 | 26.9212607 | 0.000157 |
| *F47B8.4* | 2.772681923 | 2.619286297 | 14.69424123 | 0.001968439 | *lipl-1* | -1.9756966 | 2.25546171 | 34.0924968 | 5.08E-05 |
| *gadr-6* | 1.523194085 | 3.763011576 | 45.0602816 | 1.22E-05 | *nep-17* | -1.8092283 | 9.14305943 | 22.4477066 | 0.0003572 |
| *fbxa-189* | 1.639004162 | 3.657698657 | 40.25652504 | 2.20E-05 | *cest-13* | -2.163551 | 4.70891703 | 85.8090942 | 3.34E-07 |
| *F49H6.5* | 1.813996946 | 0.712543609 | 8.130349195 | 0.013254588 | *F55G11.2* | -1.575774 | 2.71043474 | 22.6897366 | 0.0003387 |
| *F52B11.2* | 1.758011165 | 6.772777242 | 211.5628172 | 1.32E-09 | *F58B4.5* | -3.1502058 | 6.68457491 | 57.6603199 | 3.28E-06 |
| *F52B11.5* | 1.939375596 | 2.407004775 | 22.52915423 | 0.000349366 | *WBGene00010543* | -1.5042864 | -0.0060562 | 8.28518125 | 0.01257676 |
| *F53B2.8* | 3.149248946 | 7.598588128 | 55.84151949 | 3.91E-06 | *K06A4.7* | -2.8107573 | 2.17071203 | 21.59066 | 0.0004231 |
| *F54B8.4* | 1.577331829 | 1.411713931 | 28.21906037 | 0.000126118 | *K08F9.1* | -2.6454103 | 0.68337555 | 43.4260479 | 1.48E-05 |
| *F54C8.7* | 1.985314271 | 7.12278356 | 116.7071857 | 5.32E-08 | *cyp-14A2* | -2.4599082 | 2.12495995 | 16.8717607 | 0.00116375 |
| *hrg-3* | 2.44458495 | 0.673493473 | 17.75653811 | 0.000945288 | *WBGene00010753* | -3.0780382 | -1.4464233 | 8.30062531 | 0.01252126 |
| *clec-229* | 1.815560403 | 1.244291744 | 5.387873245 | 0.036633287 | *K10G4.5* | -2.5953248 | -0.4540175 | 17.6185265 | 0.0009752 |
| *mpst-2* | 6.672503759 | -1.812227427 | 22.45303611 | 0.000706483 | *asah-1* | -3.5998287 | 5.48557734 | 20.8413444 | 0.00049187 |
| *H40L08.1* | 1.708263952 | 6.290325801 | 137.3469231 | 1.97E-08 | *M04C9.4* | -2.5244047 | 2.1271637 | 58.1383264 | 3.08E-06 |
| *cdr-4* | 4.298578626 | 9.34060361 | 85.31473658 | 3.53E-07 | *mfsd-11* | -2.4557282 | 3.28031539 | 49.0648477 | 7.77E-06 |
| *cbp-2* | 1.706761626 | 5.326028971 | 63.03744757 | 1.96E-06 | *clec-258* | -2.9589139 | 3.88802902 | 66.9039091 | 1.40E-06 |
| *K08D8.4* | 1.929292304 | 5.148818597 | 21.00589861 | 0.000475728 | *cyp-14A4* | -1.5496819 | 2.62737807 | 20.5517515 | 0.00051824 |
| *phf-33* | 3.141493977 | -0.766867883 | 25.34672236 | 0.000206853 | *R07E3.1* | -1.5616356 | 8.10340811 | 55.1250469 | 4.13E-06 |
| *K10D3.6* | 1.552860031 | 0.425737073 | 12.34332166 | 0.003644692 | *chil-19* | -2.2395385 | 1.86319096 | 26.6317105 | 0.00016502 |
| *lgc-49* | 1.53939563 | 0.801035777 | 21.83115651 | 0.000400469 | *chil-21* | -1.5631995 | 0.51856329 | 11.6567406 | 0.00442158 |
| *K10G4.3* | 2.991346318 | 2.192493771 | 97.00872369 | 1.62E-07 | *chil-23* | -3.0769192 | 1.71657982 | 25.7844554 | 0.00019304 |
| *sodh-1* | 2.075846709 | 8.173944803 | 23.40403734 | 0.00029791 | *R09H10.5* | -1.5821194 | 7.55227934 | 69.8642806 | 1.10E-06 |
| *M01G12.7* | 7.010586139 | 2.216293872 | 72.19386044 | 9.28E-07 | *R11D1.7* | -1.9033661 | -2.2196032 | 5.82820845 | 0.03070006 |
| *M01G12.9* | 2.755482654 | 6.962501169 | 119.9698458 | 4.50E-08 | *pho-8* | -2.0017649 | 2.75260759 | 10.1143322 | 0.00700277 |
| *M163.5* | 1.888580822 | -1.684751921 | 5.478958329 | 0.035265171 | *T01D3.6* | -2.4441909 | 6.85860434 | 28.218831 | 0.00012732 |
| *arrd-11* | 5.847912956 | 2.311400743 | 161.5139067 | 7.20E-09 | *ugt-30* | -2.9958417 | 2.51719429 | 38.9042584 | 2.63E-05 |
| *glo-1* | 1.518457158 | 3.728663584 | 28.00578342 | 0.000130675 | *cest-1.2* | -2.9035198 | 4.8916316 | 138.534119 | 1.87E-08 |
| *zip-6* | 2.628018758 | 0.832955315 | 16.96396866 | 0.001135718 | *T03E6.8* | -1.8667837 | 2.84960634 | 38.032407 | 2.94E-05 |
| *R09H10.2* | 1.67227136 | 2.625711775 | 22.65315863 | 0.000341094 | *T05E12.3* | -3.7147021 | -1.2235268 | 15.8840721 | 0.00146165 |
| *swt-6* | 1.828755328 | 4.143602748 | 20.3567924 | 0.000543199 | *T10G3.4* | -1.8907432 | -1.8503278 | 6.12859299 | 0.0273209 |
| *WBGene00011210* | 2.384911982 | -1.442982079 | 5.597125169 | 0.035003598 | *T16G1.4* | -2.0055958 | 4.88805062 | 16.4674209 | 0.00127901 |
| *R10E8.8* | 1.854111489 | 3.8204652 | 32.31239319 | 6.60E-05 | *T16G1.5* | -1.6053521 | 2.89212506 | 7.62183772 | 0.01583347 |
| *R12H7.4* | 1.588382836 | -0.803684472 | 7.36921856 | 0.017273661 | *T16G1.6* | -1.9966975 | 6.63424032 | 23.9238949 | 0.00027031 |
| *allo-1* | 1.871887292 | 7.065269973 | 230.033739 | 7.75E-10 | *T16G1.7* | -2.2103163 | 5.01818675 | 46.0575874 | 1.09E-05 |
| *T03F6.3* | 2.260230513 | 6.733758099 | 26.98155638 | 0.0001568 | *T16G12.1* | -3.4683886 | 6.49044659 | 60.0951813 | 2.61E-06 |
| *T04F8.7* | 1.611185639 | 4.618072586 | 43.44361874 | 1.48E-05 | *clec-26* | -3.0766263 | 2.02167676 | 76.0306636 | 6.75E-07 |
| *T05F1.7* | 2.704305603 | 1.037032361 | 39.01107923 | 2.58E-05 | *sysm-1* | -3.1315951 | 0.97215434 | 15.4261951 | 0.00189072 |
| *T05F1.9* | 2.339084031 | 2.589370131 | 32.81378074 | 6.13E-05 | *clec-34* | -2.1362147 | -1.2834262 | 7.70628029 | 0.01534013 |
| *cgt-1* | 1.695030461 | 4.238875595 | 21.79208943 | 0.000404941 | *clec-38* | -2.449687 | -0.7587572 | 14.1841646 | 0.00223039 |
| *nhr-213* | 1.613159778 | 2.883106051 | 30.95345884 | 8.13E-05 | *W02B12.1* | -2.2858957 | 4.68007913 | 80.5742992 | 4.82E-07 |
| *T06D8.9* | 2.489820384 | 7.443891894 | 222.7941774 | 9.50E-10 | *clec-50* | -1.5295646 | 8.83181223 | 27.7335308 | 0.00013677 |
| *WBGene00011607* | 1.592713237 | 2.534383915 | 46.44343672 | 1.04E-05 | *oac-53* | -1.6110649 | -0.3138029 | 12.6151865 | 0.00338158 |
| *T08D10.3* | 1.684920364 | 1.327926902 | 8.978916651 | 0.010006257 | *oac-54* | -1.5430845 | 1.32152522 | 6.59398909 | 0.02290771 |
| *T11B7.1* | 1.906787439 | 0.348642797 | 26.46377688 | 0.000169895 | *Y6G8.2* | -1.7575838 | 3.495071 | 30.7101663 | 8.44E-05 |
| *sqst-1* | 1.945744645 | 8.561570424 | 68.19940963 | 1.26E-06 | *comt-2* | -2.2894182 | -1.5819794 | 12.9594621 | 0.00307928 |
| *T14G8.3* | 2.030208889 | 9.257088328 | 111.0711743 | 7.18E-08 | *Y32F6A.5* | -2.2876587 | 6.94232817 | 83.1740037 | 4.01E-07 |
| *T23F6.2* | 1.810684924 | 1.253139584 | 16.83607055 | 0.001166902 | *clec-8* | -1.6784741 | 2.41260734 | 21.0949768 | 0.00046394 |
| *T23F11.6* | 2.938285186 | 0.741643322 | 27.80088294 | 0.000135233 | *vmo-1* | -1.5922719 | -0.61455 | 4.83540348 | 0.04597214 |
| *zip-10* | 3.497093108 | 5.048322593 | 50.17240972 | 7.00E-06 | *Y39G8B.7* | -3.8233106 | -2.0900167 | 10.1070362 | 0.00699704 |
| *nmat-1* | 1.960511413 | 3.638851672 | 22.09059057 | 0.000381836 | *Y40H7A.10* | -5.032898 | 3.8328264 | 76.5307909 | 6.64E-07 |
| *Y17D7B.2* | 2.696774404 | 0.870165325 | 24.17630237 | 0.000255881 | *Y47H9C.1* | -2.3593987 | 2.444663 | 11.4979144 | 0.00464422 |
| *arrd-7* | 1.542845155 | 5.00962374 | 41.91042031 | 1.78E-05 | *Y47H10A.3* | -2.1217162 | 1.75123288 | 12.5263751 | 0.00347883 |
| *arrd-8* | 2.829736418 | -0.38056419 | 19.04625554 | 0.000711463 | *Y49E10.16* | -3.571239 | 5.48544582 | 107.094433 | 9.18E-08 |
| *Y26D4A.3* | 3.613812309 | -0.065649012 | 17.43010038 | 0.00101977 | *Y51H4A.5* | -3.7449403 | 1.97689401 | 22.0403797 | 0.00038717 |
| *Y37H2A.11* | 6.356484974 | -2.153068509 | 20.96563621 | 0.000907393 | *Y69H2.9* | -1.6249742 | 2.70654301 | 40.8456203 | 2.04E-05 |
| *Y38F1A.2* | 1.79808007 | 3.12503934 | 85.6165367 | 3.38E-07 | *clec-27* | -2.0512402 | -0.4083445 | 9.49022804 | 0.0084815 |
| *Y38H6C.9* | 3.249586503 | 1.259949361 | 61.46167432 | 2.26E-06 | *cest-2.1* | -1.9732996 | 4.53397616 | 52.5123518 | 5.38E-06 |
| *Y41E3.8* | 2.42026536 | 5.439716416 | 109.2963312 | 7.91E-08 | *ugt-6* | -1.6348852 | 5.18053822 | 23.2825758 | 0.00030482 |
| *Y43F8B.12* | 2.537587497 | 0.926380923 | 43.03435642 | 1.55E-05 | *ugt-4* | -1.7945317 | 2.88315055 | 19.5037203 | 0.0006451 |
| *Y47H10A.5* | 1.979347117 | 6.290215252 | 10.96443666 | 0.005423968 | *ZK550.2* | -1.9931875 | 3.06152455 | 40.3404317 | 2.17E-05 |
| *fbxc-23* | 2.539015871 | 2.408779383 | 26.04569372 | 0.000183291 | *clec-61* | -5.9669692 | 2.06984462 | 47.5630397 | 9.31E-06 |
| *Y54G11A.4* | 1.947950255 | 3.823433942 | 73.38443529 | 8.28E-07 | *ZK673.1* | -2.5087262 | 2.6688468 | 17.0048131 | 0.00112848 |
| *Y56A3A.33* | 2.734627096 | 2.175359492 | 56.12777808 | 3.74E-06 | *clec-51* | -1.7584748 | 4.77023504 | 35.2159027 | 4.33E-05 |
| *Y60A3A.8* | 2.026630294 | 5.139563859 | 79.06068601 | 5.38E-07 | *clec-52* | -2.6997834 | 2.99440148 | 25.4515985 | 0.00020476 |
| *fbxa-89* | 2.320497701 | 2.273725057 | 28.07014723 | 0.00012928 | *WBGene00015077* | -1.6049209 | 1.67002252 | 6.55993765 | 0.02320079 |
| *fbxa-90* | 1.852225105 | 3.679571223 | 74.35551178 | 7.67E-07 | *C03A7.2* | -1.6761635 | -0.7610771 | 13.6635924 | 0.00255309 |
| *fbxa-115* | 1.523514551 | 3.70462879 | 49.90545412 | 7.09E-06 | *C03A7.13* | -1.7213615 | 1.11464009 | 13.8241024 | 0.00244816 |
| *WBGene00013763* | 1.809952197 | 5.72093843 | 42.83118271 | 1.60E-05 | *clec-10* | -2.2108218 | 4.95423889 | 48.8295348 | 8.03E-06 |
| *Y116F11B.9* | 1.51274844 | -0.639381258 | 5.166585565 | 0.04003817 | *clec-89* | -1.6076618 | 1.40809636 | 10.9777273 | 0.00538452 |
| *fbxa-30* | 4.29021012 | 2.617459957 | 51.0757797 | 6.36E-06 | *math-10* | -1.5805519 | 1.21056048 | 15.997428 | 0.00142239 |
| *fbxa-31* | 1.84923513 | 3.743300765 | 20.129216 | 0.000567568 | *C17F4.7* | -1.7328506 | 10.4267979 | 25.0949307 | 0.00021641 |
| *WBGene00013841* | 2.024397169 | 2.098845648 | 16.57137287 | 0.001243604 | *prmt-6* | -2.0636719 | 2.87223779 | 36.0378698 | 3.85E-05 |
| *fbxa-37* | 1.661509787 | 3.219895682 | 24.79405 | 0.000228514 | *C23H5.8* | -1.503661 | 6.755068 | 10.7752695 | 0.00573615 |
| *ZK836.3* | 2.57708636 | 1.995134967 | 111.9731662 | 6.84E-08 | *dod-3* | -2.9155183 | 1.19173019 | 17.9961427 | 0.00089762 |
| *ZK896.1* | 3.603203146 | 3.521908323 | 75.72090749 | 6.91E-07 | *C27D9.2* | -4.0542825 | 2.53400627 | 29.8093908 | 9.83E-05 |
| *C37A5.3* | 1.811224015 | -0.422171814 | 9.757975048 | 0.007796252 | *mfsd-13.2* | -2.0588657 | 4.64330758 | 77.4716839 | 6.06E-07 |
| *WBGene00014979* | 6.646458095 | -1.761425315 | 17.09082164 | 0.001531636 | *clec-5* | -1.6591852 | 4.89545508 | 12.2090198 | 0.00379796 |
| *faah-2* | 1.751501022 | 7.478180544 | 30.4727596 | 8.85E-05 | *cest-10* | -1.9345914 | 4.26761718 | 24.1230204 | 0.00026053 |
| *B0244.4* | 2.06973872 | -0.452806557 | 12.39032434 | 0.003597556 | *abhd-3.2* | -1.7713844 | 5.01739021 | 37.018291 | 3.38E-05 |
| *ggtb-1* | 1.620996861 | 6.284307877 | 67.58972948 | 1.32E-06 | *ilys-2* | -2.9394597 | 1.75869956 | 14.7926262 | 0.00218565 |
| *B0348.2* | 3.375525606 | 1.923395746 | 58.33673755 | 3.02E-06 | *C46F2.1* | -1.8438784 | 4.30764017 | 29.5725673 | 0.00010145 |
| *B0403.3* | 2.672506332 | 4.455480576 | 124.2671376 | 3.63E-08 | *cyp-35A4* | -2.2322244 | 1.53980583 | 25.2520518 | 0.00021039 |
| *pdi-6* | 1.613518682 | 9.972065571 | 76.82195369 | 6.36E-07 | *cest-35.2* | -2.5468214 | 4.47984982 | 22.6320964 | 0.00034511 |
| *B0507.6* | 2.271734925 | 3.168199684 | 12.71369039 | 0.003305049 | *clec-86* | -3.5170311 | 3.16565565 | 58.0412262 | 3.13E-06 |
| *dod-20* | 2.304640659 | 2.949000539 | 34.01954482 | 5.13E-05 | *klo-2* | -1.6415147 | 3.36846418 | 28.1551903 | 0.00012746 |
| *C06E1.1* | 2.461775207 | 2.506982284 | 89.84476075 | 2.55E-07 | *E02H9.9* | -1.9879201 | -0.9392492 | 5.12859636 | 0.04069261 |
| *irg-1* | 2.91120628 | 4.441307806 | 45.96619714 | 1.12E-05 | *clec-7* | -3.4472245 | 2.13737206 | 64.0571103 | 1.79E-06 |
| *WBGene00015596* | 4.416625617 | -0.389958848 | 22.40221324 | 0.000358073 | *F11D5.5* | -2.4336441 | -0.9626593 | 22.7834764 | 0.00041142 |
| *WBGene00015597* | 4.262151448 | 3.319353763 | 168.4156163 | 5.55E-09 | *pud-2.1* | -4.8846645 | 5.16787247 | 13.817562 | 0.00246329 |
| *fbxa-163* | 5.966033992 | 3.109920872 | 66.66605019 | 1.46E-06 | *pud-4* | -8.3063924 | 2.93629892 | 81.1490077 | 4.73E-07 |
| *fbxa-165* | 6.236813986 | -0.602489783 | 26.77768732 | 0.00020636 | *pud-1.1* | -9.4258325 | 0.7040734 | 70.9733071 | 3.07E-06 |
| *fbxa-158* | 4.013934885 | 2.873323268 | 46.80492412 | 1.01E-05 | *pud-2.2* | -6.2158453 | 6.75731475 | 87.99319 | 2.95E-07 |
| *C14B9.2* | 1.585769416 | 8.426017217 | 56.36956614 | 3.65E-06 | *pud-3* | -7.5608201 | 3.55227436 | 86.5179463 | 6.04E-07 |
| *nhr-155* | 1.564935633 | 0.895037292 | 5.291689411 | 0.038090339 | *F19C7.4* | -2.6519646 | 1.87555426 | 48.4506276 | 8.31E-06 |
| *C15B12.2* | 1.725618408 | -0.208313316 | 5.617550174 | 0.03337976 | *F21C10.9* | -1.7243702 | 7.5006503 | 19.4322579 | 0.00065927 |
| *math-15* | 1.925889356 | 2.492509931 | 43.91658753 | 1.40E-05 | *ilys-5* | -3.1565295 | 8.48921558 | 43.8850497 | 1.42E-05 |
| *C17F4.3* | 3.086950661 | 1.582630003 | 31.43993053 | 7.57E-05 | *F22E5.1* | -2.4686868 | 3.88911229 | 68.2467369 | 1.25E-06 |
| *C18A11.1* | 2.206837704 | 5.468306706 | 65.13096651 | 1.64E-06 | *F22E5.8* | -1.5616518 | 0.88339046 | 15.5632659 | 0.00157972 |
| *C18B2.4* | 2.00843374 | 6.90395595 | 56.68358601 | 3.56E-06 | *F22H10.6* | -1.7801731 | 0.65978877 | 17.5111554 | 0.00099922 |
| *ari-1.3* | 1.627598312 | 5.353229312 | 35.97920374 | 3.90E-05 | *F23F12.3* | -3.3107152 | 2.53131975 | 74.5196227 | 7.58E-07 |
| *WBGene00016301* | 2.946993498 | 1.315690033 | 58.459745 | 2.99E-06 | *F26G1.3* | -4.8791166 | -1.1817893 | 7.4860632 | 0.0175708 |
| *C34H4.1* | 2.318081126 | 5.695757119 | 58.39062818 | 3.06E-06 | *asp-13* | -4.0944901 | 7.70866677 | 57.2264414 | 3.42E-06 |
| *C35E7.3* | 4.740516032 | -1.9847548 | 8.650684011 | 0.012925607 | *F28B4.3* | -1.7216993 | 8.55276115 | 12.8873677 | 0.00315283 |
| *C37C3.10* | 2.61519486 | 0.324451769 | 12.38855987 | 0.003599313 | *F37C4.6* | -1.6178071 | 5.11440065 | 36.1670714 | 3.78E-05 |
| *fbxc-5* | 1.866656204 | 4.146964706 | 41.35007828 | 1.91E-05 | *btb-16* | -1.5986017 | 2.86487089 | 22.8898769 | 0.00032592 |
| *fbxc-4* | 1.671492774 | 4.135734743 | 32.12007322 | 6.80E-05 | *btb-17* | -1.9400269 | 2.17595033 | 35.3671961 | 4.23E-05 |
| *fbxc-2* | 1.671492774 | 4.135734743 | 32.12007322 | 6.80E-05 | *F40A3.7* | -1.7169916 | 1.6008275 | 18.3118276 | 0.00083517 |
| *fbxc-1* | 1.866656204 | 4.146964706 | 41.35007828 | 1.91E-05 | *F41C6.6* | -1.7473843 | -0.7232564 | 8.26344939 | 0.01266942 |
| *irg-2* | 3.01810315 | 3.61451885 | 40.28272804 | 2.22E-05 | *F42A10.7* | -4.0989618 | 3.47167112 | 94.980214 | 1.83E-07 |
| *C49G7.7* | 6.05201059 | 4.988340921 | 77.71081331 | 6.08E-07 | *WBGene00018448* | -4.1510297 | 0.11326008 | 56.8260266 | 3.49E-06 |
| *C49G7.10* | 3.839922141 | 6.017976376 | 60.23798494 | 2.57E-06 | *WBGene00018449* | -2.8726861 | -1.404927 | 15.2677891 | 0.00169829 |
| *cest-32* | 1.599848656 | 6.602693912 | 71.05513728 | 9.96E-07 | *WBGene00018450* | -2.8726861 | -1.404927 | 15.2677891 | 0.00169829 |
| *C54E4.5* | 1.943583916 | 4.000726275 | 45.63134941 | 1.14E-05 | *WBGene00018451* | -2.8726861 | -1.404927 | 15.2677891 | 0.00169829 |
| *cebp-1* | 1.792720405 | 5.792744044 | 40.75303366 | 2.08E-05 | *F48E3.2* | -1.5058542 | 3.51514933 | 13.1617855 | 0.0029203 |
| *E02C12.6* | 2.301458544 | 4.41319605 | 54.92664317 | 4.21E-06 | *F48G7.5* | -3.7327092 | 3.10246002 | 18.921809 | 0.0007355 |
| *fbxa-53* | 1.841852604 | 1.562920122 | 32.08319935 | 6.84E-05 | *drd-50* | -2.0319535 | 2.73630983 | 7.37046503 | 0.01730126 |
| *numr-1* | 3.947670795 | 2.330328988 | 15.25771971 | 0.001711038 | *F49F1.5* | -3.5284393 | 5.18076245 | 24.8443624 | 0.00022834 |
| *F10G2.7* | 2.039446354 | -0.335243307 | 16.7098827 | 0.001201729 | *F49F1.7* | -1.8392857 | 3.42399762 | 18.8972777 | 0.00073506 |
| *F14F9.2* | 4.019751281 | 1.425940541 | 7.337248292 | 0.01750686 | *oac-32* | -3.8724456 | 0.10250739 | 59.8173364 | 2.63E-06 |
| *F14F9.3* | 3.605097921 | 5.030558965 | 42.24869198 | 1.73E-05 | *irg-3* | -2.0217297 | 3.01231419 | 20.6423558 | 0.00050987 |
| *F14F9.4* | 1.529630113 | 5.031963153 | 55.33651848 | 4.04E-06 | *F56A4.3* | -5.6210279 | 5.24017392 | 5.1823556 | 0.04302859 |
| *F14H12.2* | 2.40165524 | 0.108362058 | 18.04341595 | 0.000886436 | *ugt-52* | -2.4043155 | 2.65658889 | 75.2716167 | 7.15E-07 |
| *F16B4.2* | 2.282503069 | 3.747707011 | 61.69264248 | 2.21E-06 | *nspg-6* | -1.6927136 | 6.73333129 | 43.5666329 | 1.46E-05 |
| *F19B10.5* | 2.197737254 | -1.798925746 | 4.794119179 | 0.046762238 | *F56F10.1* | -1.7843392 | 7.14152217 | 35.2319766 | 4.36E-05 |
| *F19G12.3* | 4.24912946 | -1.215252815 | 22.44565734 | 0.000355065 | *F57F4.4* | -2.1743441 | 10.5911105 | 34.8911141 | 4.53E-05 |
| *WBGene00017631* | 2.423581222 | -0.621649953 | 20.5702196 | 0.000516281 | *F58B6.1* | -1.9820538 | 0.69333191 | 19.9213344 | 0.00059069 |
| *F20B6.9* | 2.486961492 | -2.138998928 | 6.701651606 | 0.022009168 | *WBGene00019208* | -2.813758 | 0.03328993 | 14.1333289 | 0.0022701 |
| *F22E5.6* | 9.573395795 | 5.396821986 | 425.2689719 | 1.54E-11 | *H20E11.2* | -2.5137732 | 2.5266613 | 29.4254296 | 0.00010354 |
| *F22E5.13* | 1.828509415 | 3.603102594 | 23.96839448 | 0.000265927 | *H20E11.3* | -1.9351109 | 3.61143917 | 32.6564845 | 6.27E-05 |
| *F28H1.1* | 2.22883438 | 3.615331197 | 41.97206874 | 1.77E-05 | *bgal-2* | -2.4801314 | 4.99652182 | 65.7198309 | 1.56E-06 |
| *sup-36* | 2.107311191 | 4.52642924 | 143.141034 | 1.52E-08 | *K01A2.4* | -3.4694629 | 0.52094923 | 25.7822289 | 0.00019144 |
| *F40B5.2* | 1.51883286 | 5.477798657 | 63.97875075 | 1.81E-06 | *txt-4* | -1.8998889 | 5.07010326 | 27.7391115 | 0.00013792 |
| *F41B4.3* | 3.615346614 | 3.461860737 | 123.103046 | 3.85E-08 | *cyp-35B2* | -3.829804 | 2.82361073 | 46.1416207 | 1.08E-05 |
| *WBGene00018325* | 2.687765023 | 1.028909879 | 12.08845462 | 0.00392798 | *cyp-35B1* | -2.8589358 | 3.85260973 | 39.5571444 | 2.42E-05 |
| *F42C5.3* | 2.303152457 | 1.261344075 | 8.62737573 | 0.011244793 | *cyp-35A5* | -2.1079849 | 4.11588672 | 32.767128 | 6.19E-05 |
| *fbxa-182* | 1.663060523 | 4.566287169 | 32.66831377 | 6.26E-05 | *WBGene00019532* | -1.7720815 | 0.76010868 | 17.5070116 | 0.00100016 |
| *F43H9.4* | 1.758296127 | 4.742190267 | 26.62397165 | 0.000166732 | *K09C4.1* | -2.4438872 | 1.55691627 | 30.2967294 | 9.01E-05 |
| *nhr-185* | 2.076631139 | -1.831372605 | 6.177130586 | 0.026816778 | *K09C4.4* | -2.4766386 | -0.6487012 | 21.4610027 | 0.00043104 |
| *sec-22* | 1.666592227 | 5.628091768 | 41.16957252 | 1.97E-05 | *cyp-35A3* | -2.2052669 | 3.8628209 | 24.4631817 | 0.00024474 |
| *F56D2.5* | 1.810294802 | 4.565570841 | 49.85779068 | 7.13E-06 | *fil-2* | -2.0213097 | -1.0591886 | 5.85271636 | 0.0317201 |
| *F57B9.3* | 5.241469691 | 4.480082359 | 98.63639022 | 1.50E-07 | *K12H4.7* | -1.5649583 | 9.33570941 | 21.9046973 | 0.0003947 |
| *spg-20* | 2.640217578 | 5.146786176 | 92.89226561 | 2.09E-07 | *trx-3* | -1.8684423 | 4.37003566 | 19.4366225 | 0.00065866 |
| *fbxc-3* | 1.671492774 | 4.135734743 | 32.12007322 | 6.80E-05 | *R05D8.9* | -3.5991876 | 2.26181716 | 17.9905266 | 0.00090236 |
| *WBGene00019379* | 1.614838826 | 1.21683023 | 4.688591732 | 0.048958932 | *R05D8.11* | -2.2481388 | -2.3399794 | 7.70282078 | 0.01631933 |
| *K09D9.1* | 3.927819451 | 5.27899087 | 57.84073254 | 3.22E-06 | *aagr-2* | -1.5484816 | 9.10134501 | 74.5432135 | 7.56E-07 |
| *oac-57* | 1.810799476 | 0.48907803 | 16.81673655 | 0.001172161 | *clec-43* | -2.9179234 | 0.62026841 | 13.00057 | 0.00305798 |
| *set-12* | 2.605707348 | 2.460486621 | 70.95177898 | 1.00E-06 | *R07C12.1* | -4.1330461 | 1.25598062 | 22.1160523 | 0.00038147 |
| *pgph-1* | 5.69976705 | -0.811223703 | 6.245800909 | 0.027405352 | *WBGene00019934* | -3.3897416 | -0.6306365 | 29.1133384 | 0.0001089 |
| *swt-7* | 1.564698118 | 6.317537292 | 64.88488315 | 1.67E-06 | *lipl-3* | -2.4313327 | -0.1741489 | 18.5603855 | 0.00079071 |
| *K11H12.3* | 2.613555428 | -0.738922532 | 7.894681922 | 0.014401753 | *T02H6.9* | -2.4387292 | -1.1798092 | 6.09467672 | 0.02772368 |
| *PDB1.1* | 1.996342443 | 6.268608759 | 63.85017754 | 1.83E-06 | *ugt-53* | -3.2950026 | 3.03128335 | 80.2539038 | 4.93E-07 |
| *R02D3.8* | 1.867979942 | 4.860121528 | 65.47091698 | 1.59E-06 | *clec-53* | -3.4611269 | 2.654718 | 28.6907519 | 0.0001178 |
| *R03H10.6* | 2.743019447 | 3.147587696 | 49.50855654 | 7.40E-06 | *T05E7.1* | -2.5627159 | 4.10349956 | 51.8233738 | 5.78E-06 |
| *R03H10.7* | 2.181861985 | 3.311313873 | 88.11216916 | 2.86E-07 | *math-38* | -2.3373361 | 0.13695564 | 16.4120819 | 0.00128878 |
| *T03F1.6* | 2.158888663 | 2.180652338 | 19.21774679 | 0.000687171 | *dach-1* | -5.4882154 | 2.09359134 | 76.6068237 | 6.51E-07 |
| *T05A8.2* | 4.726133673 | -1.247301351 | 6.48129511 | 0.030328411 | *T10B5.7* | -1.5561804 | 5.17120164 | 17.9035562 | 0.00092006 |
| *bath-46* | 1.848223821 | 3.804704102 | 64.30021186 | 1.76E-06 | *math-42* | -2.8816127 | 1.43376449 | 60.3187237 | 2.51E-06 |
| *T08E11.1* | 4.163039102 | 0.831044484 | 27.36227237 | 0.000146965 | *nep-22* | -1.5986457 | 7.77454996 | 22.8765068 | 0.00032929 |
| *WBGene00020364* | 3.104291699 | 0.864892333 | 68.39196006 | 1.24E-06 | *clec-218* | -3.3592279 | 4.5659774 | 25.4911243 | 0.00020333 |
| *fbxa-60* | 1.955810231 | 6.202807889 | 18.93158003 | 0.000733947 | *W03D8.8* | -2.6780838 | 2.3952512 | 16.9403554 | 0.00114541 |
| *WBGene00020458* | 2.564994971 | -0.101243024 | 19.71775181 | 0.000616525 | *cpr-8* | -3.7839575 | 4.56767228 | 69.5732021 | 1.15E-06 |
| *T13C5.6* | 2.939042574 | 4.884947814 | 64.75496745 | 1.72E-06 | *gba-4* | -2.9370705 | 6.39411282 | 31.7864565 | 7.23E-05 |
| *T19C3.4* | 1.865592312 | 6.255033664 | 90.99069783 | 2.36E-07 | *irld-53* | -1.5471778 | 1.47546415 | 8.49925079 | 0.01170488 |
| *T19D12.4* | 1.988246899 | 7.911277231 | 46.69163758 | 1.01E-05 | *pud-1.2* | -6.3719705 | 8.19530346 | 69.2855003 | 1.17E-06 |
| *cnp-3* | 2.295068814 | 5.659045584 | 35.37907581 | 4.27E-05 | *nspg-12* | -2.7373551 | 3.14748217 | 53.7303064 | 4.75E-06 |
| *T23F4.2* | 1.554927792 | 5.044581238 | 23.64885523 | 0.000284526 | *spp-23* | -2.4362187 | 3.31383857 | 81.2043885 | 4.61E-07 |
| *T24A6.7* | 2.40163178 | -0.266168198 | 25.61398079 | 0.000197233 | *comt-4* | -1.7715908 | 0.84371715 | 8.8760935 | 0.01033496 |
| *T24C4.4* | 2.449643483 | 3.527033946 | 24.02284965 | 0.000265395 | *Y40C7B.4* | -2.1779396 | -2.1942932 | 5.78755382 | 0.03119413 |
| *T24E12.5* | 2.889183772 | 5.567578031 | 52.53124143 | 5.46E-06 | *clec-174* | -2.9239032 | 0.8013004 | 10.6942607 | 0.00587617 |
| *W02C12.2* | 1.520310077 | 2.813453908 | 51.84296052 | 5.77E-06 | *clec-71* | -1.5970945 | 0.09730042 | 7.08662 | 0.01911642 |
| *WBGene00020981* | 2.073717231 | 4.77010592 | 14.52666577 | 0.002053426 | *tig-3* | -4.6521599 | -0.7210951 | 9.41091772 | 0.00872207 |
| *clec-125* | 1.586916181 | 0.903685314 | 11.09084676 | 0.005208427 | *daao-1* | -2.5899799 | 3.01074401 | 15.6323368 | 0.00156139 |
| *svh-11* | 3.38208759 | -0.065724196 | 68.54640822 | 1.22E-06 | *clec-210* | -3.647625 | 1.32043016 | 37.3240377 | 3.23E-05 |
| *Y22D7AR.2* | 1.532559388 | 3.591070863 | 24.98252807 | 0.000220846 | *nspg-8* | -2.700763 | 3.86688506 | 54.6238046 | 4.34E-06 |
| *clec-121* | 7.531568679 | -0.537082972 | 34.75808332 | 6.32E-05 | *ZC196.4* | -2.4622899 | 1.15638946 | 19.498511 | 0.00064582 |
| *Y32G9A.5* | 4.687897259 | -1.264834197 | 12.53214032 | 0.004373295 | *ZK6.6* | -1.5770783 | 0.44620102 | 5.46906278 | 0.03545427 |
| *Y34F4.4* | 3.164030412 | 4.023362518 | 28.53126997 | 0.000120925 | *lipl-5* | -1.8996332 | 8.55664146 | 39.3821361 | 2.46E-05 |
| *Y39A3A.4* | 1.599521171 | 2.305939639 | 16.23324342 | 0.001344549 | *ZK697.14* | -3.2973817 | -1.560108 | 11.9566871 | 0.00406073 |
| *comt-3* | 2.715400534 | 6.598764589 | 45.15433921 | 1.23E-05 | *ZK1193.2* | -1.8493172 | 5.28707459 | 110.702243 | 7.33E-08 |
| *WBGene00021520* | 2.067287175 | 3.452744376 | 26.12484869 | 0.000180346 | *WBGene00023313* | -3.0263701 | -0.5943668 | 7.62107682 | 0.01580397 |
| *Y45G5AM.3* | 1.67825863 | 5.188947274 | 97.83243207 | 1.54E-07 | *clec-80* | -1.5771508 | 5.16235698 | 65.3787132 | 1.60E-06 |
| *Y46H3A.5* | 3.494441782 | 1.678083547 | 34.55482867 | 4.77E-05 | *F11D5.7* | -3.4709064 | 0.43108884 | 18.3988765 | 0.00118052 |
| *WBGene00021737* | 7.967279284 | 0.126858891 | 43.82190107 | 2.07E-05 | *H39E23.3* | -3.7011831 | -2.1197003 | 5.77127162 | 0.03855543 |
| *fbxa-48* | 3.904010939 | -0.495975908 | 47.03282553 | 9.74E-06 | *clec-25* | -1.6675988 | -1.5068634 | 5.0046879 | 0.04281017 |
| *fbxa-66* | 4.92651975 | 2.050605095 | 161.9030179 | 7.10E-09 | *clec-36* | -3.0605428 | 1.42307704 | 55.5757326 | 3.95E-06 |
| *atln-1* | 1.784508397 | 8.002133231 | 83.01473621 | 4.05E-07 | *F23F12.13* | -2.0997154 | 1.71971866 | 53.033395 | 5.10E-06 |
| *Y54G2A.18* | 2.922432552 | 8.544046398 | 149.8892735 | 1.15E-08 | *F23G4.1* | -3.1681142 | -0.5617615 | 35.8392914 | 3.96E-05 |
| *manf-1* | 1.660475181 | 6.932348403 | 81.20710487 | 4.61E-07 | *folt-3* | -2.0122966 | -1.60453 | 10.2216727 | 0.00675558 |
| *Y54G2A.36* | 1.962808031 | 3.306100994 | 20.87396855 | 0.000486332 | *T19H5.6* | -3.3834031 | 0.5023092 | 35.547259 | 4.13E-05 |
| *Y58A7A.3* | 2.102201632 | 7.246741991 | 54.36611771 | 4.48E-06 | *F26G1.11* | -1.7181469 | 2.33543539 | 12.1073394 | 0.00389924 |
| *Y58A7A.4* | 2.721621274 | 2.864428693 | 25.19964578 | 0.000214191 | *Y70C5A.3* | -2.4327155 | 1.17595802 | 26.0329736 | 0.00018317 |
| *Y58A7A.5* | 3.878532937 | 6.307958002 | 114.5007394 | 6.14E-08 | *E02H4.7* | -2.7774133 | 3.08765504 | 27.3009455 | 0.0001485 |
| *Y71G12B.2* | 2.683103927 | 3.008501621 | 92.71944453 | 2.11E-07 | *C39B5.14* | -1.5542662 | 0.52039175 | 6.0703633 | 0.02794042 |
| *pho-9* | 1.959435086 | 4.721542868 | 18.45559964 | 0.000814119 | *F57C12.6* | -2.3572182 | -1.1120756 | 7.71724626 | 0.01528159 |
| *fbxa-25* | 3.801334181 | 0.925369207 | 52.21825928 | 5.55E-06 | *WBGene00194769* | -2.5032872 | 1.82960566 | 29.3072688 | 0.00010553 |
| *fbxa-26* | 2.338601447 | 3.393308081 | 18.78486683 | 0.000757646 | *F19C6.8* | -3.2809492 | -1.7134569 | 14.2945529 | 0.00246686 |
| *fbxa-79* | 1.512614272 | 3.262554337 | 12.22825726 | 0.003767943 | *K12B6.11* | -1.5928155 | 1.03882677 | 8.88673109 | 0.01029908 |
| *fbxa-80* | 1.644392607 | 2.668606021 | 19.35021003 | 0.000666544 | *F59D8.3* | -7.2535612 | -1.3998345 | 6.67731142 | 0.02478315 |
| *fbxa-19* | 1.568440961 | 3.495434254 | 27.44660253 | 0.000143552 | *T27A10.8* | -6.228405 | -0.5450452 | 7.53924661 | 0.01629911 |
| *fbxa-138* | 1.977540562 | 0.102670652 | 18.63622395 | 0.000777699 | *WBGene00202499* | -4.0619189 | -1.8520297 | 5.24384301 | 0.04399367 |
| *pals-17* | 1.538617016 | 3.921298308 | 11.2655558 | 0.004966705 | *W04H10.5* | -2.6797052 | 1.26306592 | 18.3649748 | 0.00149899 |
| *Y102A11A.9* | 4.180684905 | 4.113963048 | 127.3975268 | 3.12E-08 | *K08D10.14* | -2.3267024 | 2.83956028 | 99.0006248 | 1.43E-07 |
| *WBGene00022487* | 1.868559622 | 1.457398382 | 26.89861405 | 0.000157608 | *T25G12.13* | -1.5122737 | 0.58607857 | 8.17529559 | 0.01305354 |
| *WBGene00022545* | 2.476196369 | 1.765737362 | 14.1393553 | 0.002266588 | *F22E5.23* | -1.503388 | -0.6319773 | 6.44963483 | 0.02418071 |
| *sdz-35* | 9.386040671 | 5.194006589 | 276.6969041 | 2.48E-10 |  |  |  |  |  |
| *ZC239.13* | 1.559415328 | -1.22993932 | 5.069278965 | 0.041677909 |  |  |  |  |  |
| *ZC239.14* | 4.244698983 | 3.131825123 | 132.6393683 | 2.44E-08 |  |  |  |  |  |
| *ZK177.9* | 5.624555033 | -1.829198385 | 16.34835705 | 0.001793644 |  |  |  |  |  |
| *ZK418.7* | 2.758939847 | 4.52162847 | 28.77671647 | 0.000116157 |  |  |  |  |  |
| *ZK1055.6* | 2.081540882 | 5.831818346 | 75.27090091 | 7.15E-07 |  |  |  |  |  |
| *ZK1055.7* | 1.811402077 | 6.685243147 | 97.85831782 | 1.54E-07 |  |  |  |  |  |
| *F43C1.7* | 2.885664733 | 1.192857667 | 32.4087549 | 6.51E-05 |  |  |  |  |  |
| *Y75B8A.39* | 5.107167694 | 2.820637535 | 78.04923558 | 5.87E-07 |  |  |  |  |  |
| *F40F12.9* | 3.717978654 | 2.739588874 | 66.31437512 | 1.47E-06 |  |  |  |  |  |
| *F15H10.10* | 5.235160735 | -0.33845692 | 9.62764771 | 0.009633696 |  |  |  |  |  |
| *tag-229* | 1.896112605 | 4.570857601 | 141.4918748 | 1.64E-08 |  |  |  |  |  |
| *zipt-7.1* | 1.88175836 | 6.028369064 | 106.7668676 | 9.11E-08 |  |  |  |  |  |
| *C54C6.7* | 2.477645346 | 0.482027895 | 16.71458022 | 0.001200411 |  |  |  |  |  |
| *Y6G8.5* | 2.617112046 | 1.041448553 | 15.02697593 | 0.001811439 |  |  |  |  |  |
| *C30H6.12* | 4.171488611 | 1.309796264 | 44.86703544 | 1.25E-05 |  |  |  |  |  |
| *Y71G12B.32* | 2.836448266 | 1.910420671 | 41.04017036 | 1.99E-05 |  |  |  |  |  |
| *K10G6.5* | 3.457417043 | -1.035647338 | 8.226188438 | 0.012852266 |  |  |  |  |  |
| *Y82E9BL.18* | 3.198944531 | 3.346493782 | 55.66009997 | 3.92E-06 |  |  |  |  |  |
| *T08A9.13* | 2.71338215 | 0.271583677 | 29.36924204 | 0.000104478 |  |  |  |  |  |
| *F41E6.15* | 2.373891642 | -1.063545507 | 7.334508555 | 0.017488458 |  |  |  |  |  |
| *B0205.13* | 1.515126896 | 4.831128425 | 16.08378423 | 0.001400703 |  |  |  |  |  |
| *B0205.14* | 2.79936494 | 2.540165804 | 32.91824217 | 6.03E-05 |  |  |  |  |  |
| *B0303.16* | 2.114269634 | -1.383281467 | 10.70063011 | 0.005845811 |  |  |  |  |  |
| *T06E6.15* | 1.947406296 | 0.369249884 | 11.76919119 | 0.004282117 |  |  |  |  |  |
| *M01B2.13* | 2.069553104 | 2.184054334 | 20.53439818 | 0.000520096 |  |  |  |  |  |
| *F26D11.12* | 1.563567575 | 1.75089681 | 15.4891787 | 0.001608528 |  |  |  |  |  |
| *F26D11.13* | 2.464666781 | -0.330218399 | 7.953393409 | 0.014092484 |  |  |  |  |  |
| *eol-1* | 2.022979181 | 1.532676901 | 16.63456878 | 0.001223087 |  |  |  |  |  |
| *C25F9.11* | 4.218005163 | 3.955044162 | 51.98166596 | 5.78E-06 |  |  |  |  |  |
| *C25F9.12* | 4.005891488 | 1.70190023 | 34.29537995 | 4.99E-05 |  |  |  |  |  |
| *Y43F8B.15* | 4.491637299 | 2.348388004 | 94.7746936 | 1.86E-07 |  |  |  |  |  |
| *Y37H2A.14* | 2.783340351 | 6.432705491 | 39.13624896 | 2.57E-05 |  |  |  |  |  |
| *Y37H2A.13* | 3.489610038 | -1.349505799 | 13.0954824 | 0.003797372 |  |  |  |  |  |
| *F33H12.7* | 4.118074143 | 6.516671115 | 83.27451019 | 4.07E-07 |  |  |  |  |  |
| *C25F9.14* | 1.775882407 | 4.765997732 | 38.80766839 | 2.65E-05 |  |  |  |  |  |
| *C41G7.8* | 2.067260166 | 3.377893915 | 13.269386 | 0.002845656 |  |  |  |  |  |
| *C49G7.12* | 3.770726853 | 4.018235823 | 59.76032627 | 2.69E-06 |  |  |  |  |  |
| *WBGene00077629* | 5.375172669 | 1.623683703 | 200.0571932 | 1.88E-09 |  |  |  |  |  |
| *K08D8.7* | 1.792672949 | 2.821366271 | 70.94949803 | 1.00E-06 |  |  |  |  |  |
| *C33D9.13* | 3.053331718 | 4.335014185 | 127.4220113 | 3.12E-08 |  |  |  |  |  |
| *B0462.5* | 2.839454575 | 2.923324004 | 53.65397356 | 4.79E-06 |  |  |  |  |  |
| *F19B10.13* | 4.140521492 | 0.348981551 | 33.6151314 | 5.44E-05 |  |  |  |  |  |
| *ZC239.22* | 1.864013998 | 1.138255216 | 12.16562575 | 0.003829461 |  |  |  |  |  |
| *C25F9.16* | 4.013757512 | -0.806331888 | 15.20456978 | 0.001981021 |  |  |  |  |  |
| *Y43F8B.23* | 2.566821808 | 2.022983216 | 45.98763249 | 1.10E-05 |  |  |  |  |  |
| *K08D8.12* | 2.874384312 | 2.671163182 | 28.6033024 | 0.000119502 |  |  |  |  |  |
| *nlp-78* | 1.5820787 | -1.435296058 | 5.128536865 | 0.040670112 |  |  |  |  |  |
| *Y43F8B.25* | 4.189922815 | 1.068544684 | 85.00151054 | 3.53E-07 |  |  |  |  |  |
| *anr-16* | 2.512652038 | -0.873371461 | 16.33066183 | 0.001313833 |  |  |  |  |  |
| *anr-32* | 1.834720234 | 0.442717881 | 15.07670071 | 0.001780444 |  |  |  |  |  |
| *anr-42* | 2.286472377 | -1.820162001 | 10.44398575 | 0.006314313 |  |  |  |  |  |
| *linc-64* | 3.45484641 | -1.491967604 | 24.44050462 | 0.000392321 |  |  |  |  |  |
| *Y14H12A.4* | 2.150069353 | 1.976994097 | 62.37871186 | 2.08E-06 |  |  |  |  |  |
| *C54F6.18* | 3.057463384 | 0.185136721 | 12.30226609 | 0.003700803 |  |  |  |  |  |
| *C49C8.8* | 3.038353664 | -0.388031537 | 9.901030844 | 0.007456543 |  |  |  |  |  |
| *WBGene00269338* | 4.897624972 | 1.914694175 | 50.60577387 | 6.68E-06 |  |  |  |  |  |
| *C49G7.13* | 4.998120202 | -1.057552635 | 21.66769983 | 0.00041365 |  |  |  |  |  |
| *Y41C4A.32* | 8.200385535 | 9.918314264 | 637.0255194 | 1.04E-12 |  |  |  |  |  |
| *H05L14.3* | 2.759534637 | -1.08571704 | 20.79207311 | 0.000493363 |  |  |  |  |  |
| *W07G1.25* | 4.257700673 | -1.081020851 | 16.10801373 | 0.001605561 |  |  |  |  |  |
| *F19B10.14* | 2.390398616 | -0.852155223 | 6.467680355 | 0.024017131 |  |  |  |  |  |
| *WBGene00303363* | 2.243436264 | -0.062113635 | 22.47095512 | 0.000353327 |  |  |  |  |  |
| *T07H8.11* | 3.782259532 | 0.968919831 | 8.456569218 | 0.011901948 |  |  |  |  |  |
| *R193.6* | 2.805714394 | -0.68728444 | 10.39802072 | 0.006402742 |  |  |  |  |  |

Table S7

| **Up - HSP-3 only** | **Up - HSP-4 only** | **Up in both** | **Down - HSP-3 only** | **Down -HSP-4 only** | **Down in both** |
| --- | --- | --- | --- | --- | --- |
| *abf-4* | *acr-18* | *warf-1* | *col-123* | *amt-4* | *abu-9* |
| *aip-1* | *bre-1* | *WBGene00000191* | *dhs-15* | *aqp-1* | *cpr-2* |
| *cyp-14A5* | *bro-1* | *ckb-2* | *grl-27* | *asm-2* | *ech-9* |
| *dhs-9* | *cnd-1* | *gem-4* | *hsp-3* | *asm-3* | *lbp-8* |
| *WBGene00001091* | *crn-4* | *gst-38* | *pqn-91* | *asp-1* | *smf-3* |
| *far-3* | *cul-6* | *rab-11.2* | *srh-1* | *asp-2* | *spp-4* |
| *his-25* | *daf-14* | *rrf-2* | *sri-40* | *asp-6* | *vit-1* |
| *hsp-16.1* | *dnj-28* | *skr-5* | *srv-13* | *cpr-4* | *vit-3* |
| *hsp-16.2* | *dsc-4* | *sri-36* | *str-41* | *cpr-5* | *vit-4* |
| *hsp-16.11* | *fis-2* | *sri-39* | *tbh-1* | *dhs-2* | *cyp-25A1* |
| *hsp-16.41* | *fkh-3* | *sri-74* | *C10C5.4* | *dhs-14* | *WBGene00009230* |
| *hsp-16.48* | *gcy-13* | *fbxa-199* | *C26G2.2* | *gfi-1* | *oac-20* |
| *hsp-17* | *gpa-6* | *arrd-3* | *sdz-6* | *gst-10* | *clec-24* |
| *hsp-70* | *gst-9* | *tba-7* | *pals-38* | *hsp-4* | *clec-28* |
| *nas-3* | *his-17* | *tbb-6* | *bgnt-1.3* | *ins-35* | *clec-33* |
| *pqn-97* | *ins-34* | *tsp-1* | *F29D10.2* | *lys-1* | *F49C12.7* |
| *pqn-98* | *kgb-2* | *tsp-2* | *F36D1.8* | *lys-4* | *F55G11.2* |
| *srx-111* | *lec-8* | *C08E8.4* | *H25K10.1* | *lys-7* | *cyp-14A2* |
| *str-144* | *mig-21* | *fbxa-98* | *K01D12.8* | *lys-8* | *WBGene00010753* |
| *pals-26* | *nhr-3* | *fbxc-58* | *T13F3.6* | *lys-10* | *M04C9.4* |
| *WBGene00007181* | *nhr-6* | *C31H5.7* | *chil-25* | *nuc-1* | *cyp-14A4* |
| *C04F12.1* | *nhr-117* | *C32H11.9* | *clec-144* | *pcp-1* | *chil-23* |
| *rnh-1.3* | *nsf-1* | *C45B11.2* | *clec-4* | *pcp-3* | *ugt-30* |
| *fipr-23* | *sid-2* | *fbxa-141* | *nhr-234* | *pmp-5* | *T05E12.3* |
| *oac-14* | *sip-1* | *F11D11.3* | *clec-247* | *trpl-5* | *sysm-1* |
| *ifas-2* | *sir-2.3* | *F15B9.6* | *pals-30* | *spp-1* | *comt-2* |
| *F14F8.8* | *skr-3* | *F47B8.4* | *cest-2.3* | *spp-11* | *Y40H7A.10* |
| *F19B2.5* | *srh-116* | *F53B2.8* | *ZK218.4* | *spp-16* | *clec-61* |
| *F20G2.1* | *sri-67* | *mpst-2* | *ZK1025.2* | *srx-68* | *WBGene00015077* |
| *fipr-26* | *sri-70* | *cdr-4* | *C04G6.2* | *str-7* | *clec-10* |
| *gmd-2* | *srp-2* | *K10G4.3* | *C07G1.7* | *str-168* | *ilys-2* |
| *cyp-13A5* | *srp-7* | *M01G12.7* | *C09B8.4* | *tag-10* | *pud-2.1* |
| *pals-13* | *srp-8* | *M01G12.9* | *C18A11.4* | *wrt-7* | *pud-4* |
| *Y38E10A.22* | *srw-86* | *M163.5* | *C18H7.1* | *C08B6.2* | *pud-1.1* |
| *Y75B12B.3* | *str-82* | *arrd-11* | *C30G12.2* | *ugt-22* | *pud-2.2* |
| *fbxa-50* | *str-124* | *zip-6* | *C53B7.2* | *cbl-1* | *pud-3* |
| *clec-60* | *str-245* | *R12H7.4* | *D1014.6* | *hrg-7* | *F23F12.3* |
| *ZK970.7* | *syx-2* | *T23F11.6* | *D1014.7* | *ugt-23* | *F42A10.7* |
| *cyp-35A1* | *ubc-23* | *zip-10* | *oac-12* | *cest-9.1* | *WBGene00018448* |
| *C06E4.6* | *zmp-1* | *arrd-8* | *F07E5.7* | *cest-7* | *WBGene00018449* |
| *C14C6.3* | *plin-1* | *Y26D4A.3* | *F14H12.7* | *C31H5.6* | *WBGene00018450* |
| *C14C6.6* | *B0457.6* | *Y37H2A.11* | *folt-2* | *C55A1.6* | *WBGene00018451* |
| *WBGene00015761* | *C06B3.7* | *Y38H6C.9* | *F45D11.15* | *ttr-44* | *F48G7.5* |
| *C14C6.8* | *C06C3.7* | *Y54G11A.4* | *F45D11.16* | *cest-3* | *oac-32* |
| *pals-32* | *C11E4.8* | *fbxa-30* | *F47B7.4* | *clec-17* | *F56A4.3* |
| *fbxa-12* | *pals-6* | *ZK896.1* | *F49D11.3* | *F01D5.1* | *WBGene00019208* |
| *WBGene00016526* | *C23H4.6* | *B0348.2* | *F54D10.8* | *F01D5.2* | *K01A2.4* |
| *D1022.5* | *fbxa-82* | *dod-20* | *F56A4.2* | *F01D5.3* | *cyp-35A5* |
| *EEED8.12* | *C25F9.5* | *irg-1* | *H01M10.2* | *F01D5.5* | *clec-43* |
| *F13A2.2* | *dod-21* | *WBGene00015596* | *hacd-1* | *acox-1.4* | *WBGene00019934* |
| *F25A2.1* | *C34C12.4* | *WBGene00015597* | *T10B5.8* | *clec-227* | *ugt-53* |
| *math-28* | *zip-5* | *fbxa-163* | *clec-118* | *clec-57* | *clec-53* |
| *fbxb-53* | *C44H9.4* | *fbxa-165* | *clec-209* | *clec-54* | *math-38* |
| *F48G7.13* | *gale-1* | *fbxa-158* | *Y27F2A.9* | *F09B12.3* | *dach-1* |
| *K06H6.1* | *mfb-1* | *nhr-155* | *clec-70* | *F09C8.1* | *math-42* |
| *K06H6.2* | *E04D5.4* | *fbxc-5* | *plep-1* | *cpt-5* | *clec-218* |
| *K06H6.4* | *ugt-44* | *fbxc-4* | *WBGene00022209* | *stdh-2* | *pud-1.2* |
| *WBGene00019454* | *F01D5.7* | *fbxc-2* | *srt-13* | *ctsa-1.2* | *comt-4* |
| *fil-2* | *gfat-1* | *fbxc-1* | *ZK105.6* | *F14D7.6* | *clec-71* |
| *M60.7* | *F08G2.5* | *C49G7.7* | *WBGene00044503* | *cyp-13A12* | *F11D5.7* |
| *T05A8.7* | *F13A7.11* | *C49G7.10* | *C04G6.13* | *cyp-35D1* | *H39E23.3* |
| *lgc-1* | *F16B12.4* | *numr-1* | *T14G12.12* | *clec-42* | *F23G4.1* |
| *phat-5* | *F17C11.11* | *F14F9.3* | *BE0003N10.6* | *F17B5.1* | *folt-3* |
| *bath-25* | *plp-2* | *F22E5.6* | *F54D8.10* | *cth-1* | *T19H5.6* |
| *W09G12.7* | *pals-14* | *F41B4.3* | *WBGene00304827* | *asah-2* | *T27A10.8* |
| *Y41D4B.15* | *F23B2.10* | *WBGene00018325* |  | *thn-1* |  |
| *Y94H6A.10* | *F35E8.2* | *fbxa-182* |  | *thn-2* |  |
| *fbxa-35* | *WBGene00009428* | *F57B9.3* |  | *cyp-37B1* |  |
| *fbxa-36* | *cbp-3* | *fbxc-3* |  | *clec-64* |  |
| *ZK488.5* | *fbxa-144* | *K09D9.1* |  | *F35E12.6* |  |
| *ZK488.6* | *ipla-2* | *pgph-1* |  | *clec-166* |  |
| *ZK1240.1* | *F47B8.3* | *R03H10.6* |  | *clec-165* |  |
| *ZK1240.8* | *gadr-6* | *T03F1.6* |  | *clec-169* |  |
| *ZC21.10* | *fbxa-189* | *T08E11.1* |  | *argk-1* |  |
| *C01G10.17* | *F49H6.5* | *WBGene00020364* |  | *lipl-2* |  |
| *WBGene00045188* | *F52B11.2* | *cnp-3* |  | *WBGene00009838* |  |
| *fbxc-12* | *F52B11.5* | *T24E12.5* |  | *clec-31* |  |
| *Y26D4A.21* | *F54B8.4* | *WBGene00020981* |  | *clec-32* |  |
| *C39B5.14* | *F54C8.7* | *clec-121* |  | *F49C12.4* |  |
| *pals-20* | *hrg-3* | *Y34F4.4* |  | *nspg-11* |  |
| *K01A6.8* | *clec-229* | *comt-3* |  | *F49E12.10* |  |
| *WBGene00271786* | *H40L08.1* | *fbxa-48* |  | *WBGene00009991* |  |
| *H04D03.7* | *cbp-2* | *fbxa-66* |  | *lipl-1* |  |
|  | *K08D8.4* | *Y58A7A.3* |  | *nep-17* |  |
|  | *phf-33* | *Y58A7A.4* |  | *cest-13* |  |
|  | *K10D3.6* | *Y58A7A.5* |  | *F58B4.5* |  |
|  | *lgc-49* | *Y71G12B.2* |  | *WBGene00010543* |  |
|  | *sodh-1* | *fbxa-25* |  | *K06A4.7* |  |
|  | *glo-1* | *fbxa-138* |  | *K08F9.1* |  |
|  | *R09H10.2* | *Y102A11A.9* |  | *K10G4.5* |  |
|  | *swt-6* | *WBGene00022545* |  | *asah-1* |  |
|  | *WBGene00011210* | *sdz-35* |  | *mfsd-11* |  |
|  | *R10E8.8* | *ZC239.14* |  | *clec-258* |  |
|  | *allo-1* | *ZK177.9* |  | *R07E3.1* |  |
|  | *T03F6.3* | *Y75B8A.39* |  | *chil-19* |  |
|  | *T04F8.7* | *F40F12.9* |  | *chil-21* |  |
|  | *T05F1.7* | *Y6G8.5* |  | *R09H10.5* |  |
|  | *T05F1.9* | *Y71G12B.32* |  | *R11D1.7* |  |
|  | *cgt-1* | *K10G6.5* |  | *pho-8* |  |
|  | *nhr-213* | *Y82E9BL.18* |  | *T01D3.6* |  |
|  | *T06D8.9* | *T08A9.13* |  | *cest-1.2* |  |
|  | *WBGene00011607* | *F41E6.15* |  | *T03E6.8* |  |
|  | *T08D10.3* | *B0205.13* |  | *T10G3.4* |  |
|  | *T11B7.1* | *B0205.14* |  | *T16G1.4* |  |
|  | *sqst-1* | *B0303.16* |  | *T16G1.5* |  |
|  | *T14G8.3* | *M01B2.13* |  | *T16G1.6* |  |
|  | *T23F6.2* | *eol-1* |  | *T16G1.7* |  |
|  | *nmat-1* | *C25F9.11* |  | *T16G12.1* |  |
|  | *Y17D7B.2* | *C25F9.12* |  | *clec-26* |  |
|  | *arrd-7* | *Y43F8B.15* |  | *clec-34* |  |
|  | *Y38F1A.2* | *Y37H2A.14* |  | *clec-38* |  |
|  | *Y41E3.8* | *F33H12.7* |  | *W02B12.1* |  |
|  | *Y43F8B.12* | *WBGene00077629* |  | *clec-50* |  |
|  | *Y47H10A.5* | *C33D9.13* |  | *oac-53* |  |
|  | *fbxc-23* | *B0462.5* |  | *oac-54* |  |
|  | *Y56A3A.33* | *F19B10.13* |  | *Y6G8.2* |  |
|  | *Y60A3A.8* | *ZC239.22* |  | *Y32F6A.5* |  |
|  | *fbxa-89* | *Y43F8B.25* |  | *clec-8* |  |
|  | *fbxa-90* | *anr-16* |  | *vmo-1* |  |
|  | *fbxa-115* | *anr-32* |  | *Y39G8B.7* |  |
|  | *WBGene00013763* | *Y14H12A.4* |  | *Y47H9C.1* |  |
|  | *Y116F11B.9* | *C54F6.18* |  | *Y47H10A.3* |  |
|  | *fbxa-31* | *WBGene00269338* |  | *Y49E10.16* |  |
|  | *WBGene00013841* | *Y41C4A.32* |  | *Y51H4A.5* |  |
|  | *fbxa-37* | *H05L14.3* |  | *Y69H2.9* |  |
|  | *ZK836.3* | *F19B10.14* |  | *clec-27* |  |
|  | *C37A5.3* |  |  | *cest-2.1* |  |
|  | *WBGene00014979* |  |  | *ugt-6* |  |
|  | *faah-2* |  |  | *ugt-4* |  |
|  | *B0244.4* |  |  | *ZK550.2* |  |
|  | *ggtb-1* |  |  | *ZK673.1* |  |
|  | *B0403.3* |  |  | *clec-51* |  |
|  | *pdi-6* |  |  | *clec-52* |  |
|  | *B0507.6* |  |  | *C03A7.2* |  |
|  | *C06E1.1* |  |  | *C03A7.13* |  |
|  | *C14B9.2* |  |  | *clec-89* |  |
|  | *C15B12.2* |  |  | *math-10* |  |
|  | *math-15* |  |  | *C17F4.7* |  |
|  | *C17F4.3* |  |  | *prmt-6* |  |
|  | *C18A11.1* |  |  | *C23H5.8* |  |
|  | *C18B2.4* |  |  | *dod-3* |  |
|  | *ari-1.3* |  |  | *C27D9.2* |  |
|  | *WBGene00016301* |  |  | *mfsd-13.2* |  |
|  | *C34H4.1* |  |  | *clec-5* |  |
|  | *C35E7.3* |  |  | *cest-10* |  |
|  | *C37C3.10* |  |  | *abhd-3.2* |  |
|  | *irg-2* |  |  | *C46F2.1* |  |
|  | *cest-32* |  |  | *cyp-35A4* |  |
|  | *C54E4.5* |  |  | *cest-35.2* |  |
|  | *cebp-1* |  |  | *clec-86* |  |
|  | *E02C12.6* |  |  | *klo-2* |  |
|  | *fbxa-53* |  |  | *E02H9.9* |  |
|  | *F10G2.7* |  |  | *clec-7* |  |
|  | *F14F9.2* |  |  | *F11D5.5* |  |
|  | *F14F9.4* |  |  | *F19C7.4* |  |
|  | *F14H12.2* |  |  | *F21C10.9* |  |
|  | *F16B4.2* |  |  | *ilys-5* |  |
|  | *F19B10.5* |  |  | *F22E5.1* |  |
|  | *F19G12.3* |  |  | *F22E5.8* |  |
|  | *WBGene00017631* |  |  | *F22H10.6* |  |
|  | *F20B6.9* |  |  | *F26G1.3* |  |
|  | *F22E5.13* |  |  | *asp-13* |  |
|  | *F28H1.1* |  |  | *F28B4.3* |  |
|  | *sup-36* |  |  | *F37C4.6* |  |
|  | *F40B5.2* |  |  | *btb-16* |  |
|  | *F42C5.3* |  |  | *btb-17* |  |
|  | *F43H9.4* |  |  | *F40A3.7* |  |
|  | *nhr-185* |  |  | *F41C6.6* |  |
|  | *sec-22* |  |  | *F48E3.2* |  |
|  | *F56D2.5* |  |  | *drd-50* |  |
|  | *spg-20* |  |  | *F49F1.5* |  |
|  | *WBGene00019379* |  |  | *F49F1.7* |  |
|  | *oac-57* |  |  | *irg-3* |  |
|  | *set-12* |  |  | *ugt-52* |  |
|  | *swt-7* |  |  | *nspg-6* |  |
|  | *K11H12.3* |  |  | *F56F10.1* |  |
|  | *PDB1.1* |  |  | *F57F4.4* |  |
|  | *R02D3.8* |  |  | *F58B6.1* |  |
|  | *R03H10.7* |  |  | *H20E11.2* |  |
|  | *T05A8.2* |  |  | *H20E11.3* |  |
|  | *bath-46* |  |  | *bgal-2* |  |
|  | *fbxa-60* |  |  | *txt-4* |  |
|  | *WBGene00020458* |  |  | *cyp-35B2* |  |
|  | *T13C5.6* |  |  | *cyp-35B1* |  |
|  | *T19C3.4* |  |  | *WBGene00019532* |  |
|  | *T19D12.4* |  |  | *K09C4.1* |  |
|  | *T23F4.2* |  |  | *K09C4.4* |  |
|  | *T24A6.7* |  |  | *cyp-35A3* |  |
|  | *T24C4.4* |  |  | *fil-2* |  |
|  | *W02C12.2* |  |  | *K12H4.7* |  |
|  | *clec-125* |  |  | *trx-3* |  |
|  | *svh-11* |  |  | *R05D8.9* |  |
|  | *Y22D7AR.2* |  |  | *R05D8.11* |  |
|  | *Y32G9A.5* |  |  | *aagr-2* |  |
|  | *Y39A3A.4* |  |  | *R07C12.1* |  |
|  | *WBGene00021520* |  |  | *lipl-3* |  |
|  | *Y45G5AM.3* |  |  | *T02H6.9* |  |
|  | *Y46H3A.5* |  |  | *T05E7.1* |  |
|  | *WBGene00021737* |  |  | *T10B5.7* |  |
|  | *atln-1* |  |  | *nep-22* |  |
|  | *Y54G2A.18* |  |  | *W03D8.8* |  |
|  | *manf-1* |  |  | *cpr-8* |  |
|  | *Y54G2A.36* |  |  | *gba-4* |  |
|  | *pho-9* |  |  | *irld-53* |  |
|  | *fbxa-26* |  |  | *nspg-12* |  |
|  | *fbxa-79* |  |  | *spp-23* |  |
|  | *fbxa-80* |  |  | *Y40C7B.4* |  |
|  | *fbxa-19* |  |  | *clec-174* |  |
|  | *pals-17* |  |  | *tig-3* |  |
|  | *WBGene00022487* |  |  | *daao-1* |  |
|  | *ZC239.13* |  |  | *clec-210* |  |
|  | *ZK418.7* |  |  | *nspg-8* |  |
|  | *ZK1055.6* |  |  | *ZC196.4* |  |
|  | *ZK1055.7* |  |  | *ZK6.6* |  |
|  | *F43C1.7* |  |  | *lipl-5* |  |
|  | *F15H10.10* |  |  | *ZK697.14* |  |
|  | *tag-229* |  |  | *ZK1193.2* |  |
|  | *zipt-7.1* |  |  | *WBGene00023313* |  |
|  | *C54C6.7* |  |  | *clec-80* |  |
|  | *C30H6.12* |  |  | *clec-25* |  |
|  | *T06E6.15* |  |  | *clec-36* |  |
|  | *F26D11.12* |  |  | *F23F12.13* |  |
|  | *F26D11.13* |  |  | *F26G1.11* |  |
|  | *Y37H2A.13* |  |  | *Y70C5A.3* |  |
|  | *C25F9.14* |  |  | *E02H4.7* |  |
|  | *C41G7.8* |  |  | *C39B5.14* |  |
|  | *C49G7.12* |  |  | *F57C12.6* |  |
|  | *K08D8.7* |  |  | *WBGene00194769* |  |
|  | *C25F9.16* |  |  | *F19C6.8* |  |
|  | *Y43F8B.23* |  |  | *K12B6.11* |  |
|  | *K08D8.12* |  |  | *F59D8.3* |  |
|  | *nlp-78* |  |  | *WBGene00202499* |  |
|  | *anr-42* |  |  | *W04H10.5* |  |
|  | *linc-64* |  |  | *K08D10.14* |  |
|  | *C49C8.8* |  |  | *T25G12.13* |  |
|  | *C49G7.13* |  |  | *F22E5.23* |  |
|  | *W07G1.25* |  |  |  |  |
|  | *WBGene00303363* |  |  |  |  |
|  | *T07H8.11* |  |  |  |  |
|  | *R193.6* |  |  |  |  |

Table S8
